# Comparing end-tidal CO_2_, respiration volume per time (RVT), and average gray matter signal for mapping cerebrovascular reactivity amplitude and delay with breath-hold task BOLD fMRI

**DOI:** 10.1101/2022.11.28.517116

**Authors:** Kristina M. Zvolanek, Stefano Moia, Joshua N. Dean, Rachael C. Stickland, César Caballero-Gaudes, Molly G. Bright

## Abstract

Cerebrovascular reactivity (CVR), defined as the cerebral blood flow response to a vasoactive stimulus, is an imaging biomarker with demonstrated utility in a range of diseases and in typical development and aging processes. A robust and widely implemented method to map CVR involves using a breath-hold task during a BOLD fMRI scan. Recording end-tidal CO_2_ (P_ET_CO_2_) changes during the breath-hold task is recommended to be used as a reference signal for modeling CVR amplitude in standard units (%BOLD/mmHg) and CVR delay in seconds. However, obtaining reliable P_ET_CO_2_ recordings requires equipment and task compliance that may not be achievable in all settings. To address this challenge, we investigated two alternative reference signals to map CVR amplitude and delay in a lagged general linear model (lagged-GLM) framework: respiration volume per time (RVT) and average gray matter BOLD response (GM-BOLD). In 8 healthy adults with multiple scan sessions, we compare spatial agreement of CVR maps from RVT and GM-BOLD to those generated with P_ET_CO_2_. We define a threshold to determine whether a P_ET_CO_2_ recording has “sufficient” quality for CVR mapping and perform these comparisons in 16 datasets with sufficient P_ET_CO_2_ and 6 datasets with insufficient P_ET_CO_2_. When P_ET_CO_2_ quality is sufficient, both RVT and GM-BOLD produce CVR amplitude maps that are nearly identical to those from P_ET_CO_2_ (after accounting for differences in scale), with the caveat they are not in standard units to facilitate between-group comparisons. CVR delays are comparable to P_ET_CO_2_ with an RVT regressor but may be underestimated with the average GM-BOLD regressor. Importantly, when P_ET_CO_2_ quality is insufficient, RVT and GM-BOLD CVR recover reasonable CVR amplitude and delay maps, provided the participant attempted the breath-hold task. Therefore, our framework offers a solution for achieving high quality CVR maps in both retrospective and prospective studies where sufficient P_ET_CO_2_ recordings are not available and especially in populations where obtaining reliable measurements is a known challenge (e.g., children). Our results have the potential to improve the accessibility of CVR mapping and to increase the prevalence of this promising metric of vascular health.

## 1. Introduction

The regulation of cerebral blood flow (CBF) is critical to maintain proper brain function. One mechanism that allows for tight regulation of CBF is the dilation and constriction of arterioles to increase or decrease blood flow, respectively. This mechanism can be characterized by a metric called cerebrovascular reactivity (CVR), defined as the CBF response to a vasoactive stimulus. It represents the ability of the brain’s blood vessels to dilate or constrict and is thus an indicator of vascular health. CVR has gained attention in recent years as an imaging biomarker in a range of pathologies, including stroke (Krainik et al., 2005), atherosclerotic disease (Donahue et al., 2014), multiple sclerosis (Marshall et al., 2014), moyamoya disease (Mikulis et al., 2005), sickle cell anemia (Václavů et al., 2019), and brain tumors (Fierstra et al., 2018), among others. In addition, changes in CVR throughout developmental (Leung et al., 2016b) and aging (McKetton et al., 2018) processes have been reported.

CVR measurements require two components: 1) a vasoactive stimulus to elicit a change in blood flow, and 2) a measure of the CBF response. An established approach for CVR measurements involves administering carbon dioxide (CO_2_) gas via a face mask during an MRI scan (Fierstra et al., 2013; Liu et al., 2019; Sleight et al., 2021). CO_2_ acts as a vasodilator, causing a systemic increase in blood flow, and the resulting blood flow response throughout the brain can be detected by MRI. Most commonly, the blood oxygenation level-dependent (BOLD) contrast is used as a surrogate measure of CBF. Alternatively, arterial spin labeling (ASL) may be used to obtain CBF in quantitative units, but it is currently limited by its low SNR, poor temporal resolution, and modeling challenges due to altered labeling efficiency in the hypercapnic state (Pinto et al., 2021).

Other guidance for exemplar CVR measurements include normalizing the blood flow response to the CO_2_ change to make the units quantitative and accounting for regional delays in the CVR response. For quantitative CVR measurements, it is necessary to record the changes in arterial CO_2_ throughout the gas challenge (Kastrup et al., 2001; Liu et al., 2019; Sleight et al., 2021; Tancredi and Hoge, 2013). Arterial CO_2_ measurements are invasive, so end-tidal CO_2_, the partial pressure of CO_2_ at the end of an exhale, may be used as a surrogate (McSwain et al., 2010; Peebles et al., 2007). By accounting for end-tidal CO_2_ changes (typically in units of mmHg), CVR can be reported as the blood flow response per unit change of CO_2_ (%BOLD/mmHg for BOLD or ΔCBF/mmHg for ASL). Additionally, it is important to consider not only the amplitude, but also the timing, or delay, of the blood flow response (Bright et al., 2009; Chang et al., 2008; Duffin et al., 2015; Moia et al., 2020a). Variations in the response time may occur due to regional variations in the timing of arterial blood arrival and local regulation of vessel diameter (Donahue et al., 2016). Accounting for this delay is not only important to achieve accurate CVR amplitudes but also serves as a separate metric of vascular health that is sensitive to cerebrovascular pathology (Donahue et al., 2016; Leung et al., 2016a; Sam et al., 2016; Stickland et al., 2021; Thomas et al., 2014; Thrippleton et al., 2018).

While gas challenges are typically recommended for robust CVR measurements, the equipment for gas delivery is expensive and requires technical expertise to operate safely (Liu and De Vis, 2019). In addition, some participants find wearing a mask uncomfortable and may experience increased claustrophobia or anxiety, introducing uncontrolled confounding factors in the data (Urback et al., 2017). Breathing tasks, such as breath-holds (Ratnatunga and Adiseshiah, 1990), deep breaths (Bright et al., 2009; Sousa et al., 2014), and intermittent breath modulations (Liu et al., 2020), are feasible alternatives to gas challenges. By modulating endogenous CO_2_ levels, these tasks serve as vasoactive stimuli and can also produce reliable estimates of CVR. Breath-holds are a particularly common breathing modulation (Urback et al., 2017) and have been used successfully even in populations with known task-compliance challenges (Dlamini et al., 2018; Handwerker et al., 2007; Thomason et al., 2005). CVR estimates are comparable between breath-hold tasks and gas challenges (Kastrup et al., 2001; Tancredi and Hoge, 2013), with robust measurements across a range of breath-hold durations (Bright and Murphy, 2013; Magon et al., 2009).

However, obtaining accurate CVR measurements from a breath-hold task still requires reliable end-tidal CO_2_ measurements, which may not be achieved in all subjects and settings. Besides, end-tidal CO_2_ measurements require external physiological monitoring equipment (e.g., gas analyzer), which may not be available in all clinical or research imaging centers. Even in healthy adults, there are challenges in achieving successful breath-hold performance with end-tidal CO_2_ recordings. For example, in a recent study of 10 healthy adults (Moia et al., 2021), 3 subjects were excluded due to “poor performance of the breath-hold task”. There are added difficulties with cooperation in patient cohorts, particularly in those with cognitive impairments who may struggle to execute commands (Pujol et al., 1998; Schouwenaars et al., 2021). Obtaining high-quality data in younger participants also tends to be more challenging, and previous work has reported inconsistent performance of a breathing task among children and young adults (Stickland et al., 2021).

The primary complication with breath-hold data quality is obtaining end-tidal CO_2_ measurements both before and after the breath-hold, which is critical for characterizing CO_2_ changes and measuring CVR in quantitative, normalized units (Bright and Murphy, 2013; Murphy et al., 2011). This can be achieved by designing the breath-hold task with expirations both before and after the breath-hold period (Pinto et al., 2021). Unreliable estimates of these expiration end-tidal CO_2_ values may occur if the participant simply does not execute them as instructed, for example, by performing a brief inspiration instead. In addition, end-tidal CO_2_ measurements are typically acquired via a nasal cannula, which requires a participant to breathe through their nose for the duration of the experiment. Lapses in nose-breathing or variations in the pressure of exhaled air may also lead to inaccurate end-tidal values.

In this work, we aimed to find alternative strategies for mapping CVR amplitude and delay that could be used in cases where end-tidal CO_2_ measurements are unavailable or unreliable. We approached this problem in breath-hold task data using a lagged-general linear model (lagged-GLM approach) to achieve more accurate CVR amplitude estimates by accounting for regional variations in CVR delay (Moia et al., 2021, 2020a; Stickland et al., 2021). We then compared results using end-tidal CO_2_, or two alternative regressors (reference signals), in the lagged-GLM.

First, we investigated another measure of respiratory physiology, respiration volume per time (RVT) (Birn et al., 2008, 2006). RVT represents changes in both the rate and depth of breathing and is obtained by continuously measuring chest position via a pressure-sensitive belt worn around the chest or abdomen. RVT is an attractive alternative to end-tidal CO_2_ because it also captures whether the participant attempts the breath-hold task. Even if the end-tidal CO_2_ measurements do not reflect a change during the apnea period, there will be a decrease in RVT due to the pause in breathing. RVT and end-tidal CO_2_ are highly correlated, have similar overlap in the variance they explain in the BOLD signal, and consistent latencies at which they affect the BOLD signal (Chang and Glover, 2009). Additionally, a respiration belt is commonly found in most scanner set-ups, making it potentially more accessible than end-tidal CO_2_ measurements.

Second, we investigated a data-driven regressor using the average gray matter BOLD timeseries (GM-BOLD). The main advantage of the GM-BOLD signal is that no external monitoring equipment is required. Changes in the BOLD timeseries should be evident provided the participant attempted the breath-hold and achieved periods of hypercapnia (Bright and Murphy, 2013; Stickland et al., 2021). While the global BOLD signal or “refined” GM-BOLD regressors have been used in other CVR methods, including techniques that capture both amplitude and delay (Geranmayeh et al., 2015; Liu et al., 2017; Tong et al., 2011; Tong and Frederick, 2014; van Niftrik et al., 2016), our proposed approach simultaneously models other regressors (e.g., motion confounds) when searching for the optimum delay of the reference signal and outputs amplitude maps normalized to the input regressor amplitude (Moia et al., 2020a).

The aim of this work was to test if RVT or GM-BOLD timeseries can be used in a lagged-GLM framework to achieve estimates of CVR amplitude and delay that are spatially similar to those generated with the gold standard of end-tidal CO_2,_ with the caveat that these alternative CVR amplitude measurements will no longer be in the standard, normalized units (%BOLD/mmHg) that are recommended for CVR comparisons across people and sessions (Kastrup et al., 2001; Murphy et al., 2011; Pinto et al., 2021; Sleight et al., 2021). We assess the agreement between CVR amplitude and delay maps in breath-hold fMRI datasets with high-quality or “sufficient” end-tidal CO_2_ data, and in those where end-tidal CO_2_ measurements were sub-optimal or “insufficient”. We hypothesized that in a lagged-GLM framework, using RVT and GM-BOLD as reference signals would produce CVR amplitude and delay measurements that are highly correlated with those produced by high-quality end-tidal CO_2_ measurements. In cases with unreliable end-tidal CO_2_ measurements, we hypothesized that RVT or GM-BOLD timeseries could be used to recover reasonable CVR amplitude and delay maps, provided that the participant attempted the breath-hold task.

## 2. Methods

### 2.1 Participants

A subset of the imaging and physiological data used in this manuscript have been published previously (Moia et al., 2021, 2020b). The full dataset includes ten healthy subjects (5F, 24-40y at the start of the experiment) with no history of psychiatric or neurological disorders. All subjects completed ten MRI sessions, which were scheduled exactly one week apart at the same time of day. MRI scanning took place using a 3T Siemens PrismaFit scanner with a 64-channel head coil. The study was approved by the Basque Center on Cognition, Brain and Language ethics committee. Informed consent was obtained before each MRI session.

Eight of the ten subjects were included in this analysis (sub-002, sub-003, sub-004, sub-006, sub-007, sub-008, sub-009, sub-010), based on those with sufficient data quality in the same two consecutive sessions (ses-02 and ses-03). In addition, two additional sessions were included from three of the subjects (sub-006, sub-009, sub-010) to capture two consecutive sessions (ses- 07 and ses-08 for sub-006 and sub-010; ses-08 and ses-09 for sub-009) with insufficient end-tidal CO_2_ timeseries (i.e., low power in the dominant frequency range of the breath-hold task, described in greater detail in Section 2.4.1). These eight subjects have similar demographics to the complete ten (4F, 27-40 yrs).

### 2.2 Data collection

#### 2.2.1 Magnetic resonance imaging

Subjects underwent a variety of task-based and resting-state acquisitions during each MRI session, but the current study focuses on the multi-echo fMRI acquisition during a breath-hold (BH) task. The multi-echo fMRI protocol was a T2*-weighted, simultaneous multislice (multiband, or MB), gradient-echo echo planar imaging sequence provided by the Center for Magnetic Resonance Research (CMRR, Minnesota) with the following parameters: 340 volumes, TR = 1.5 s, TEs = 10.6/28.69/46.78/64.87/82.96 ms, flip angle = 70°, MB acceleration factor = 4, GRAPPA = 2, 52 slices with interleaved acquisition, partial Fourier = 6/8, FoV = 211 × 211 mm^2^, voxel size = 2.4 × 2.4 × 3 mm^3^, phase encoding = AP, bandwidth = 2470 Hz/px, LeakBlock kernel reconstruction (Cauley et al., 2014) and SENSE coil combination (Sotiropoulos et al., 2013). Prior to the fMRI acquisition, single-band reference (SBRef) images were collected for each echo time to facilitate functional realignment and masking, and a pair of spin-echo echo planar images with opposite phase-encoding (AP or PA) directions and identical volume layout (TR = 2920 ms, TE = 28.6 ms, flip angle = 70°) were acquired to estimate field distortions. For anatomical co-registration and tissue segmentation, a T1-weighted MP2RAGE (TR = 5 s, TE = 2.98 ms, TI1 = 700 ms, TI2 = 2.5 s, flip angle 1 = 4°, flip angle 2 = 5°, GRAPPA = 3, 176 slices, FoV read = 256 mm, voxel size = 1 × 1 × 1 mm 3, TA = 662 s) and a T2-weighted Turbo Spin Echo image (TR = 3.39 s, TE = 389 ms, GRAPPA = 2, 176 slices, FoV read = 256 mm, voxel size = 1 × 1 × 1 mm^3^, TA = 300 s) were acquired. All DICOM files were transformed into NIFTI files with dcm2nii and formatted into Brain Imaging Data Structure (Gorgolewski et al., 2016) with heudiconv (Halchenko et al., 2019).

#### 2.2.2 Physiological data

During scanning, expired CO_2_ and O_2_ pressures were recorded via a nasal cannula (Intersurgical) and gas analyzer (ADInstruments ML206). Chest position was measured with a respiratory effort transducer (BIOPAC) placed around the upper abdomen, on the area of highest expansion during breathing. These measurements were then transferred to a physiological monitoring system (BIOPAC MP150) that simultaneously recorded scan triggers. Physiological signals were sampled at 10 kHz, starting before and continuing after the fMRI scan to allow for shifting of regressors. Before processing, the files were converted to BIDS with phys2bids (The phys2bids developers, 2019) and the physiological signals were decimated to 40 Hz to reduce file sizes.

#### 2.2.3 Breath-hold task

The BH task paradigm included eight repetitions of a 58 s BH trial. Within each trial, there were four paced breathing cycles (1 cycle = 3 s inhale and 3 s exhale), a 20 s BH, 3 s exhalation, and 11 s of free recovery breathing (Bright and Murphy, 2013). Participants were cued with visual instructions projected through a mirror on the head coil. A 15 s resting period was appended to the start and end of the paradigm to enable shifting of physiological regressors in subsequent analysis.

Prior to the scan, subjects were instructed about the importance of exhaling through their nose both before and after the BH period. These exhalations are critical because they provide end-tidal CO_2_ measurements to estimate arterial changes in CO_2_ achieved by each BH (Bright and Murphy, 2013). If the exhale is not performed properly or the measurement is unreliable, it is not possible to obtain a standard CVR estimate in units of %BOLD/mmHg.

### 2.3 Data analysis

The MRI images and physiological data used in this study are available on OpenNeuro at <u>doi:10.18112/openneuro.ds003192.v1.0.1</u> (Moia, Uruñuela, Ferrer, & Caballero-Gaudes, 2020). All code for pre-processing of the MRI data has been prepared to be run in a Singularity container, which is publicly available at https://git.bcbl.eu/smoia/euskalibur_container. The pre-processing pipeline is available at https://github.com/smoia/EuskalIBUR_preproc. Publicly available Python scripts, peakdet (Markello & DuPre, 2020) and phys2cvr (Moia, Vigotsky, & Zvolanek, 2022), were used for processing of CO_2_ recordings and computation of CVR parameter maps. The open- source *Rapidtide* v2.2.7 toolbox (B. deB Frederick, Salo, & Drucker, 2022) was used for exploratory analysis (see Discussion Section 4.3). Additional analysis code and details about how they were implemented for this manuscript are shared in the public GitHub repository: https://github.com/BrightLab-ANVIL/Zvolanek_2022.

#### 2.3.1 MRI pre-processing

Key MRI pre-processing steps are discussed here, and more detailed information can be found in Moia and colleagues (2021). MRI pre-processing was performed with a series of custom scripts combining FSL (Jenkinson et al., 2012), AFNI (Cox, 1996), and ANTs (Tustison et al., 2014) commands. The T2-weighted image was skull-stripped and co-registered to the MP2RAGE. The MP2RAGE was segmented into gray matter (GM), white matter (WM), and cerebrospinal fluid (CSF) tissues. Then, the MP2RAGE was normalized to a resampled version (2.5 mm resolution) of the MNI152 6^th^ generation template (FSL version, 1 mm resolution) (Grabner et al., 2006). The T2-weighted image was co-registered to the skull-stripped SBRef image of the first echo. Volume realignment of the functional data was performed using the SBRef of the first echo as the reference and applying the spatial transformation to all subsequent echoes (Jenkinson et al., 2002; Jenkinson and Smith, 2001). An optimal combination of the different echoes was created with tedana (DuPre et al., 2021, 2019), which weights each echo timeseries according to the voxelwise T2* value (Posse et al., 1999). Finally, the pair of spin-echo images with reverse phase-encoding directions was used to perform field distortion correction with Topup (Andersson et al., 2003). The optimally-combined, distortion-corrected data were used as the input for CVR modeling.

#### 2.3.2 Reference signals

Three different reference signals were generated for each dataset, as depicted in Figure 1: end-tidal CO_2_ (P_ET_CO_2_), respiration volume per time (RVT), and the average gray matter BOLD signal (GM-BOLD). End-tidal peaks were identified with a peak detection algorithm and manually reviewed. Linear interpolation was performed between the end-tidal peaks to create P_ET_CO_2_ timeseries. Finally, P_ET_CO_2_ timeseries were convolved with the two-gamma variate canonical hemodynamic response function.

**Figure 1:**
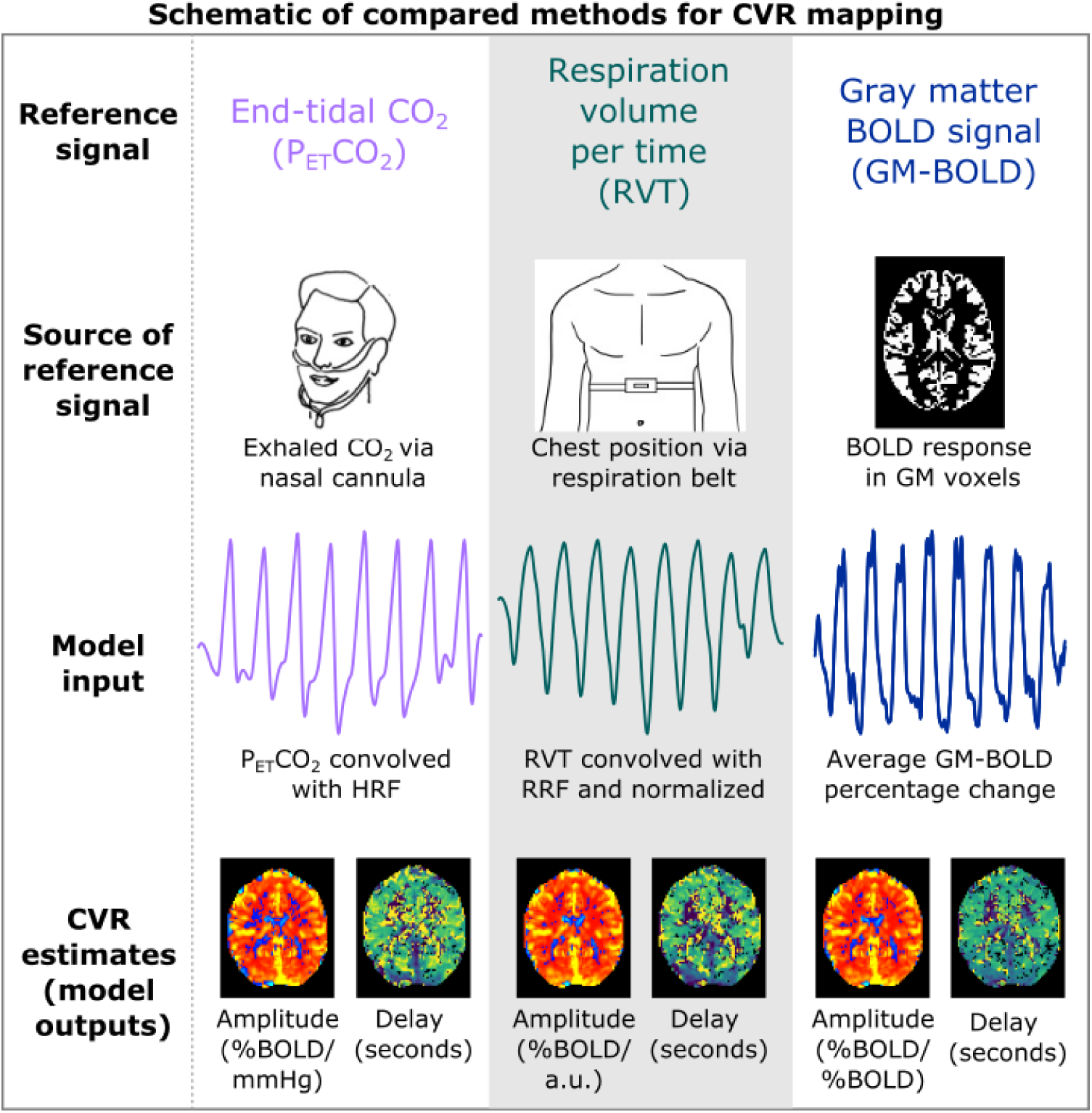
Key steps of the CVR modeling methods compared in this manuscript. Reference timeseries are generated via external recordings or the BOLD MRI data. P_ET_CO_2_ and RVT timeseries are convolved with canonical response functions. For all methods, modeling is repeated for shifted variations of each reference time signal. On a voxelwise basis, the shift that optimizes the full model R^2^ is selected. Maps of amplitude and delay are then generated using these parameters. P_ET_CO_2_ = partial pressure of end-tidal CO_2_, RVT = respiration volume per time, BOLD = blood oxygenation level dependent, GM = gray matter, HRF = hemodynamic response function, RRF = respiration response function.

Respiration recordings were processed using a custom MATLAB script. Maxima and minima in the belt trace were identified with a peak detection algorithm and manually inspected. The computation of respiration volume per time (RVT) requires alternating maxima and minima (Birn et al., 2006), but in an end-exhalation BH task, there are two consecutive minima due to exhales before and after the hold. To address this, only minima preceding the BH period were included. Linear envelopes of these maxima and minima were used to compute RVT as previously defined (Birn et al., 2006). Briefly, the difference in maxima and minima is computed at each timepoint and divided by the time between successive maxima. The RVT timeseries were then convolved with the respiration response function (RRF) (Birn et al., 2008). Importantly, all convolved RVT timeseries were z-normalized (i.e., zero mean and unit standard deviation). The normalization procedure was implemented to account for the high variability in RVT amplitudes (see Supplementary Figure S1 and Table S2). All subsequent “RVT” results refer to the convolved, normalized reference signal.

The average BOLD timeseries in GM was generated from the optimally-combined, distortion-corrected functional data with phys2cvr (Moia et al., 2022b). An eroded version of the co-registered GM mask (obtained by zeroing non-zero edge voxels within a 2.5 mm sigma Gaussian kernel with fslmaths) was used as the ROI for the average timecourse extraction. The reference signal was then expressed in signal percentage change.

#### 2.3.3 CVR amplitude and delay estimation

Voxelwise hemodynamic CVR amplitude and delay were computed using phys2cvr (Moia et al., 2022b) to implement a lagged-GLM framework that has been described previously (Moia et al., 2021, 2020a; Stickland et al., 2021). Each reference signal was considered independently from the others, but the same procedures outlined below were used for each CVR model.

First, all traces were shifted to maximize the cross-correlation with the up-sampled GM-BOLD timeseries (40 Hz to match the physiological signals). This “bulk” shift primarily accounts for measurement delay in the physiological recordings. Then, 61 shifted variants of each regressor (including the bulk shifted regressor) were created for each reference signal, in 0.3 s increments (Moia et al., 2020a). These shifts ranged ±9 s from the bulk shift. Separate GLMs were created for each shifted variant. In each case, fMRI data were modelled by a design matrix consisting of the shifted reference signal and the following nuisance regressors: Legendre polynomials up to the fourth-order, 6 realignment parameters, and their 6 temporal derivatives. Each lagged-GLM was fitted via orthogonal least squares (Moia et al., 2020a). The lagged-GLM with the maximum full model R^2^ was identified for each voxel; its corresponding shift (in seconds) determined the CVR delay, and its associated beta coefficient was extracted and rescaled to be expressed in percentage BOLD signal change (%BOLD). Therefore, the lagged-GLM generated two maps for each reference signal, as depicted in Figure 1: CVR amplitude (in units of %BOLD normalized to the amplitude of the input regressor) and CVR delay (in seconds). Delay maps were centered on the median delay across GM voxels. Both CVR amplitude and delay maps were thresholded to remove voxels at or adjacent to boundary conditions (delay = -9, -8.7, +8.7, 9 seconds) because they were considered not optimized by the lagged-GLM (Moia et al., 2020a). CVR amplitude and delay maps were normalized via nearest neighbor interpolation to the MNI152 6^th^ generation template (FSL version, 1 mm resolution) resampled to 2.5 mm resolution.

### 2.4 Data summaries and comparisons

#### 2.4.1 Determining sufficient reference signal quality

The quality of reference signals for each dataset was assessed by computing the relative power in the dominant frequency range of the BH task (0.014 to 0.020 Hz). This range is centered around 0.017 Hz, which corresponds to the 58 s BH cycle. MATLAB’s *bandpower* function was used to compute the total power between 0.014 to 0.020 Hz, as well as the total power in the signal, between 0 Hz and the Nyquist frequency (i.e., 20 Hz). Relative power was then calculated using the following equation:

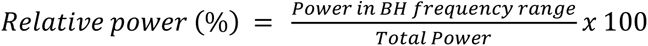

Reference signals with greater than 50% power in the BH range were deemed “sufficient”, as more than half of the signal power is in the frequency range of interest. In the time domain, this relative power threshold corresponds to reference signals with clear signal changes during each BH cycle (Figure 2). Reference signals with less than 50% power were categorized as “insufficient”.

**Figure 2:**
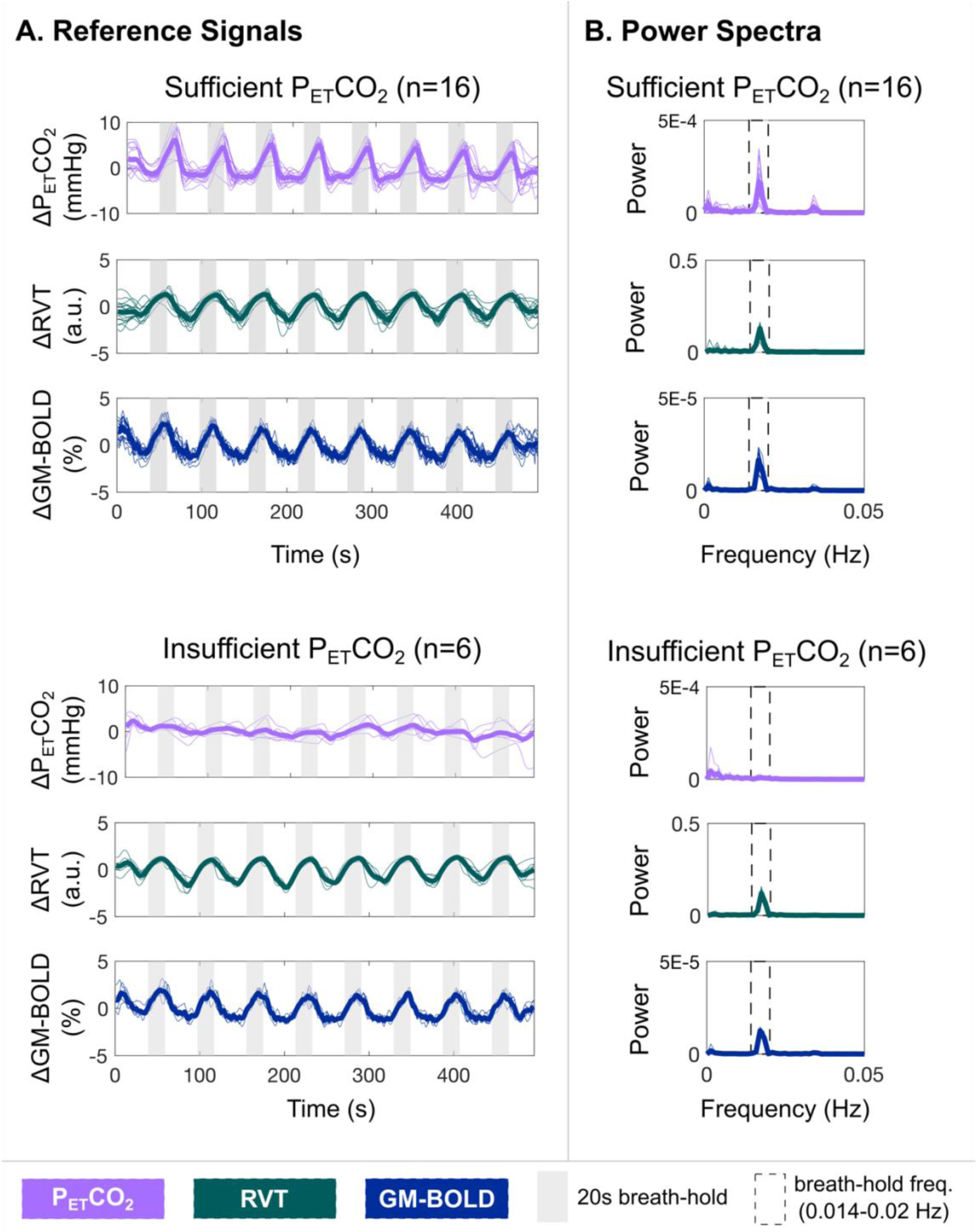
**A)** Reference signals for each dataset (thin lines) and the group average (thick lines). Gray bars indicate each 20 second breath-hold (BH) cycle. Reference signals from the three compared methods are depicted: partial pressure of end-tidal CO_2_ convolved with the hemodynamic response function (P_ET_CO_2_), respiration volume per time convolved with the respiration response function and normalized to unit variance (RVT), and average BOLD signal percentage change in gray matter (GM-BOLD). Sufficient P_ET_CO_2_ datasets (top) indicate those where the P_ET_CO_2_ timeseries has relative power >50% in the BH frequency range, while insufficient P_ET_CO_2_ datasets (bottom) indicate those where the P_ET_CO_2_ timeseries has relative power <50%. **B)** Power spectra for each dataset (thin lines) and the group average (thick lines), corresponding to the reference signals plotted in panel A. Dashed rectangles indicate the BH frequency range (0.014 to 0.020 Hz). Note that there is no peak in this range for the datasets with insufficient P_ET_CO_2_ timeseries, while a peak is visible for all other reference signals.

#### 2.4.2 Reference signal cross-correlations

Relationships between the reference signals for each dataset were assessed by computing the cross-correlation between each pair. The “bulk shifted” P_ET_CO_2_ and RVT signals were used for these comparisons, which had already been shifted to maximize the cross-correlation with the GM-BOLD signal during CVR modeling (see Section 2.3.3). The additional cross-correlation was performed to understand the relationships between signals going into the lagged-GLM and to check for any remaining offsets that may explain differences in resulting CVR maps. The GM-BOLD signal was up-sampled to 40 Hz to match the temporal resolution of the physiological signals. Using MATLAB’s *xcorr* function, cross-correlations between each pair of reference signals were computed at 0.025 s increments (i.e., 40 Hz) within a range of ±9 s. Pearson correlations (r) were transformed to Fisher’s Z values to facilitate group averaging and comparisons.

#### 2.4.3 CVR amplitude and delay values

The 98^th^ percentile of brain voxels in each CVR amplitude map (after thresholding of voxels at the boundary) was computed using the *fslstats* function in FSL. For each reference signal, the kernel density estimation of the distribution of CVR amplitude and CVR delay values was computed with MATLAB’s *ksdensity* function. Distributions were computed in gray matter using the eroded tissue mask (see section 2.3.2).

#### 2.4.4 Spatial correlations between CVR parameter maps

CVR amplitude and delay maps for each reference signal were parcellated using FSL’s Harvard-Oxford cortical atlas in MNI space (https://identifiers.org/neurovault.collection:262, <u>HarvardOxford-cort-maxprob-thr25-1mm)</u>, resampled to 2.5 mm resolution. This atlas consists of 48 cortical parcels and was further split into left and right hemispheres to generate a total of 96 cortical parcels. Then, the median CVR parameter (i.e., amplitude or delay) within each parcel was computed. The 96 median values from any two corresponding CVR parameter maps (e.g., two CVR amplitude maps) were then input to determine “spatial” correlations (i.e., at the level of the parcels).

Three different types of spatial correlations were performed:

1. **Inter-reference**: Between CVR parameter maps from different reference signals, within the same subject and session (e.g., between P_ET_CO_2_ CVR amplitude and RVT CVR amplitude for sub-002 ses-02),
2. **Inter-session**: Between CVR parameter maps from two consecutive sessions, for a given subject and reference signal (e.g., between P_ET_CO_2_ CVR amplitude maps from ses-02 and ses-03 for sub-002),
3. **Inter-quality**: Between CVR parameter maps from datasets with sufficient P_ET_CO_2_ quality and insufficient P_ET_CO_2_ quality, for a given reference signal and subject (e.g., between a sufficient P_ET_CO_2_ CVR amplitude map and an insufficient P_ET_CO_2_ CVR amplitude map for sub-006).

For all spatial correlations, the Pearson correlation coefficients were computed and transformed to Fisher’s Z. A linear model was fitted, and the beta-coefficients describing the slope were extracted. The intercept of the linear model was allowed to vary for both CVR amplitude and delay to account for potential offsets between the two inputs.

## 3. Results

In the following sections, we first describe the reference signals from all datasets in our study and distinguish those with sufficient vs. insufficient quality. Then, we show inter-reference and inter-session comparisons for datasets in which all three reference signals have sufficient quality. Next, we show inter-quality results (from two sessions) that incorporate one session with insufficient P_ET_CO_2_ quality. Finally, we present the inter-reference comparisons from only sessions with insufficient P_ET_CO_2_ quality. All comparisons are repeated for both CVR amplitude and CVR delay.

### 3.1 Reference Signals

Table 2 summarizes relative power at the BH task frequency for P_ET_CO_2_, RVT, and GM-BOLD in all datasets included in our study. This metric was used to assess data quality, and each dataset was classified as “sufficient” or “insufficient” according to the relative power in the P_ET_CO_2_ timeseries. We chose a subset of the available data, such that 16 datasets included in our study have sufficient P_ET_CO_2_ quality, and 6 datasets have insufficient P_ET_CO_2_ quality, with relative power below the 50% threshold and reaching as low as 4.33% (sub-009 ses-08). Across all datasets considered, insufficient P_ET_CO_2_ traces have 21.1±11.2% relative power (mean±stdev across subjects), while sufficient P_ET_CO_2_ traces have 68.0±6.57% relative power. Note that all RVT and GM-BOLD signals have greater than 50% relative power, with most far exceeding the threshold. Relative power in RVT and GM-BOLD signals is also generally higher than in P_ET_CO_2_, with relative power at 85.7±10.8% in RVT signals and 78.7±8.38% in GM-BOLD signals.

**Table 1:** Comparison of proposed reference signals for modeling CVR amplitude and delay

|  | **P<sub>ET</sub>CO<sub>2</sub>** | **RVT** | **GM-BOLD** |
| --- | --- | --- | --- |
| **CVR delay measurement** | Quantitative (seconds) | Quantitative (seconds) | Quantitative (seconds) |
| **CVR amplitude measurement** | Standard (%BOLD/mmHg) | Not standard, arbitrary (%BOLD/a.u.) | Not standard, unitless (%BOLD/%BOLD) |
| **Additional equipment required for acquisition** | Nasal cannula and gas monitoring system | Respiratory belt (typically comes with scanner) | N/A |
| **Head motion confounds in reference signal** | N/A | N/A | Present |
| **Sensitivity of data acquisition to participant compliance** | Participant must breathe through their nose and start and end breath-hold on exhalation | Potential bias if participant’s breathing pattern changes (e.g., stomach vs. chest or shallow vs. deep breathing), belt must be positioned correctly | N/A |

*P<sub>ET</sub>CO<sub>2</sub>* = partial pressure of end-tidal CO<sub>2</sub>, *RVT* = respiration volume per time, *GM-BOLD* = average blood oxygenation level dependent signal in gray matter.

**Table 2:**
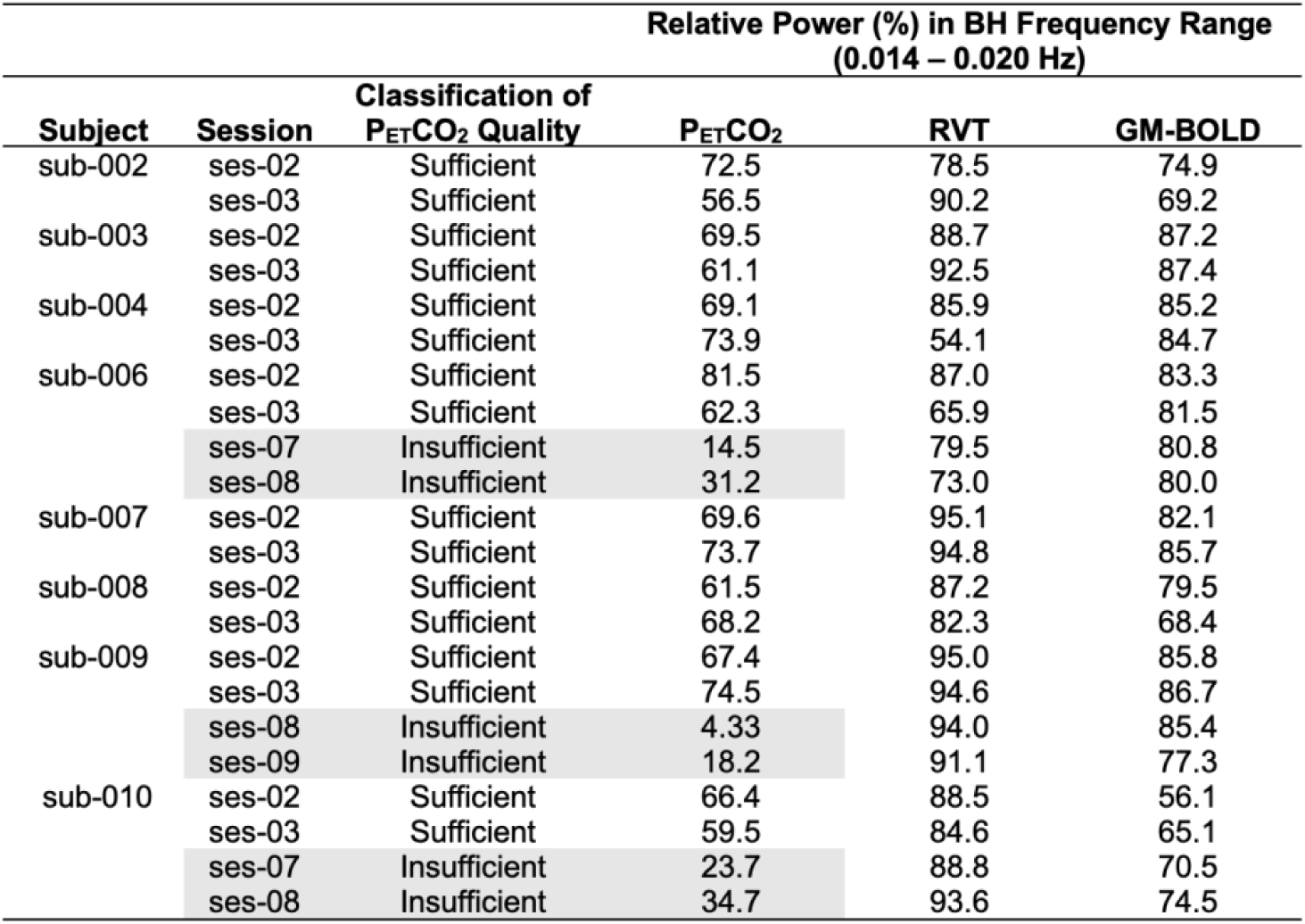
Classification of reference signals as “sufficient” or “insufficient” based on relative power in the breath-hold frequency range. “Sufficient” P_ET_CO_2_ classification is based on relative power >50%. Datasets with insufficient P_ET_CO_2_ quality are highlighted in gray.

The P_ET_CO_2_, RVT, and GM-BOLD signals for all datasets, in addition to group averages, are shown in Figure 2. Sufficient P_ET_CO_2_ datasets with P_ET_CO_2_ timeseries that have >50% relative power in the BH frequency range are plotted separately from those with insufficient P_ET_CO_2_. For sufficient P_ET_CO_2_ traces as well as all RVT and GM-BOLD traces, there are clear peaks associated with each BH cycle (indicated by the gray bars). These signal changes are expected due to periods of apnea, which increase P_ET_CO_2_ and elicit a cerebrovascular response that is detectable by BOLD fMRI. In contrast, the insufficient P_ET_CO_2_ traces lack consistent peaks for each BH cycle, and the magnitude of P_ET_CO_2_ changes is smaller. These signal characteristics likely indicate a failure to perform an exhalation before and after the BH, or exhalation through the mouth rather than the nose, which would not be captured by the nasal cannula. In these datasets, insufficient P_ET_CO_2_ traces are not due to a failure to complete the BH task, because these subjects also have clear cyclic changes in their RVT signals, indicating long durations of a stable chest position (i.e., periods of apnea).

Fig. 2B illustrates the power spectra corresponding to the reference signals in Fig. 2A. The BH frequency range is indicated by a dashed rectangle, where most of the signal power is expected. There are clear peaks within this window for sufficient P_ET_CO_2_ signals, as well as for all RVT and GM-BOLD signals. However, a peak within the BH frequency range is not evident for insufficient P_ET_CO_2_ signals, which is consistent with the lack of periodic signal changes for each BH cycle in the time domain. These power spectra also support the low relative power reported for insufficient P_ET_CO_2_ datasets in Table 2.

All reference signals are highly correlated in datasets with sufficient P_ET_CO_2_, while correlations with insufficient P_ET_CO_2_ timeseries are much lower. Relationships between each pair of reference signals were characterized by cross-correlations. These results are summarized in Supplementary Table S1. Datasets with sufficient P_ET_CO_2_ have large, positive cross-correlation amplitudes for the three reference signal comparisons (reported as mean±stdev Fisher’s Z values across subjects): P_ET_CO_2_ & RVT: 0.96±0.23, P_ET_CO_2_ & GM-BOLD: 1.19±0.22, GM-BOLD & RVT: 1.04±0.25. As expected, cross-correlations of P_ET_CO_2_ with RVT and GM-BOLD are lower in datasets with insufficient P_ET_CO_2,_ while the correlation between RVT and GM-BOLD is preserved (P_ET_CO_2_ & RVT: 0.38±0.13, P_ET_CO_2_ & GM-BOLD: 0.42±0.15, GM-BOLD & RVT: 1.20±0.22).

### 3.2 Sufficient P_ET_CO_2_ Datasets: CVR Amplitude Comparisons

#### 3.2.1 Inter-reference comparisons

CVR amplitude maps are spatially similar for all reference signals, after accounting for differences in scale, in datasets with sufficient P_ET_CO_2_ quality. Fig. 3 shows delay-optimized CVR amplitude maps generated by each reference signal. For each CVR map, the 98^th^ percentile of CVR amplitude across all brain voxels was computed (Supplementary Table S3), and this magnitude was used as the positive and negative limits of the color scale. With this scaling method, the CVR amplitude maps look nearly identical, though there are small differences particularly in voxel clusters throughout WM and CSF regions. The same relative spatial patterns are observed in all maps: higher amplitudes in cortical GM, lower amplitudes in WM, and negative amplitudes in CSF-filled regions. However, it is important to draw attention to the fact that the absolute magnitude of these CVR amplitudes is different between methods. For example, the 98^th^ percentile CVR amplitudes are 0.78±0.22 %BOLD/mmHg for P_ET_CO_2_ CVR, 1.99±0.45 %BOLD/a.u. for RVT CVR, and 2.31±0.21 %BOLD/%BOLD for GM-BOLD CVR.

**Figure 3:**
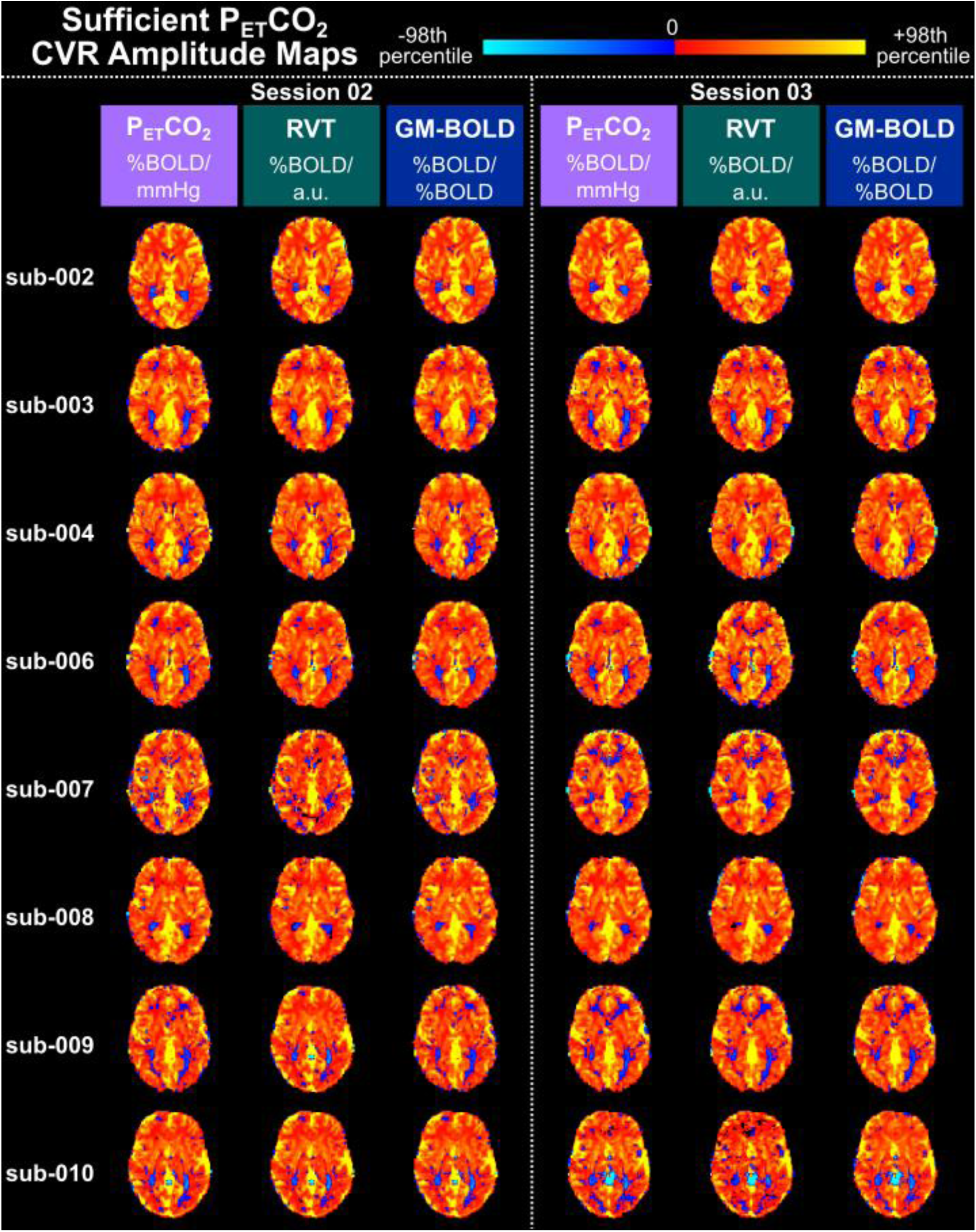
Delay-optimized CVR amplitude maps for all 16 datasets with sufficient P_ET_CO_2_ quality transformed to the MNI152 6th generation template space. For each subject, maps from session 02 are shown on the left and maps from session 03 are shown on the right. A single axial slice of the CVR map from each reference signal is shown in each column. Each CVR map is plotted on a separate color scale. The 98^th^ percentile CVR amplitude value across all voxels was computed for each map (see Table S2 for the magnitudes) and used as the positive and negative limits of the color scale. Voxels with delays at the boundary conditions have been removed. Note the different units of CVR amplitude for each reference signal.

As expected from the qualitative similarity of the CVR amplitude maps, the distributions of CVR amplitude are similar across GM voxels for each method, though they span a different range of values. Fig. 4 displays the distribution of CVR amplitude in GM for all datasets with sufficient P_ET_CO_2_. For all reference signals, the distributions of CVR amplitude are consistent both within and between subjects. Note that it may not be appropriate to interpret the range of the CVR amplitude distributions, because only P_ET_CO_2_ CVR amplitude is in meaningful units. Normalization of the RVT signal is critical to achieving these similarities in CVR amplitude, as the amplitude of the RVT measurement itself is arbitrary, with high variability even between two sessions of the same subject (see Supplementary Figure S1 and Table S2). Supplementary Figure S2 shows the distribution of CVR amplitudes without normalizing RVT and illustrates the impact on the resulting unscaled amplitude maps.

**Figure 4:**
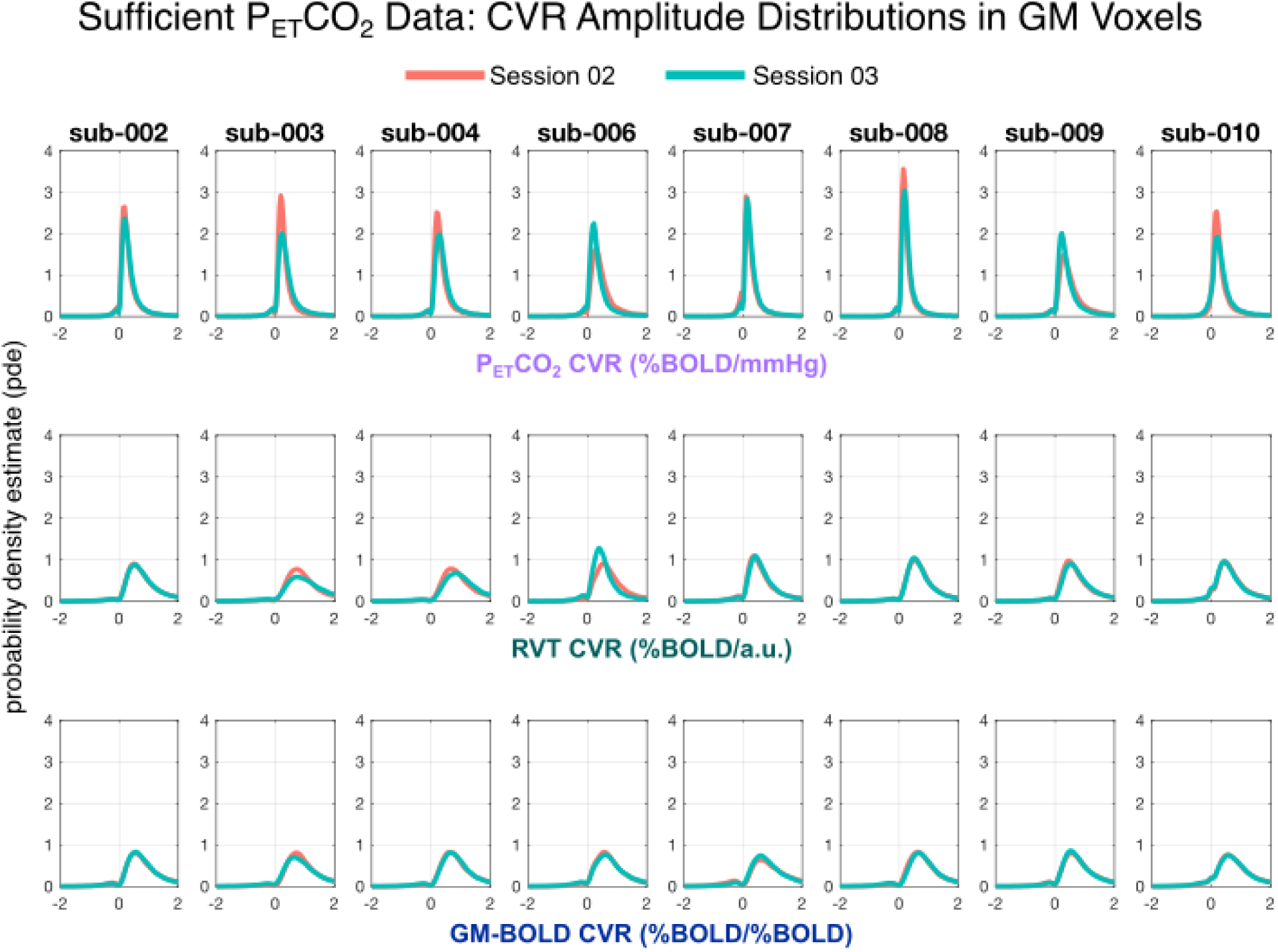
Distributions of CVR amplitude across grey matter (GM) voxels in all sufficient P_ET_CO_2_ datasets. For each subject, distributions from session 02 are plotted in orange and session 03 are plotted in teal. Each row shows the distribution of CVR amplitude for a different reference signal, with P_ET_CO_2_ CVR on top, RVT CVR in the middle, and GM-BOLD CVR on the bottom. Note that skewness of the P_ET_CO_2_ CVR distributions is different from those of the RVT CVR and GM-BOLD CVR because of the range of the plots (from -2 to +2) which matches closer to the 98% percentiles of the latter.

CVR amplitudes from each reference signal are highly correlated in datasets with sufficient P_ET_CO_2_ quality. Fig. 5A shows the spatial correlations between CVR amplitude values generated by each reference signal (inter-reference correlations) and a visual comparison of these spatial correlations from session-to-session for each subject. These comparisons are based on CVR amplitudes from cortical parcels in the Harvard-Oxford atlas. The correlation coefficients, Fisher’s Z transformed correlations, and slopes for the lines-of-best fit are also summarized in Supplementary Table S4. All group average inter-reference spatial correlations are significantly different from zero (P_ET_CO_2_ & RVT: Z=2.08, p<0.001; P_ET_CO_2_ & GM-BOLD: Z=2.26, p<0.001; GM-BOLD & RVT: Z=2.15, p<0.001). There is no significant difference between the strength of the CVR amplitude spatial correlations for each pairwise comparison between reference signals, based on a t-test adjusted for non-independent correlations (Howell, 2010) (P_ET_CO_2_ & RVT vs. P_ET_CO_2_ & GM-BOLD: T(13)=0.75, p=0.47; P_ET_CO_2_ & GM-BOLD vs. GM-BOLD & RVT: T(13)=0.40, p=0.70; P_ET_CO_2_ & RVT vs. GM-BOLD & RVT: T(13)=0.35, p=0.73).

**Figure 5:**
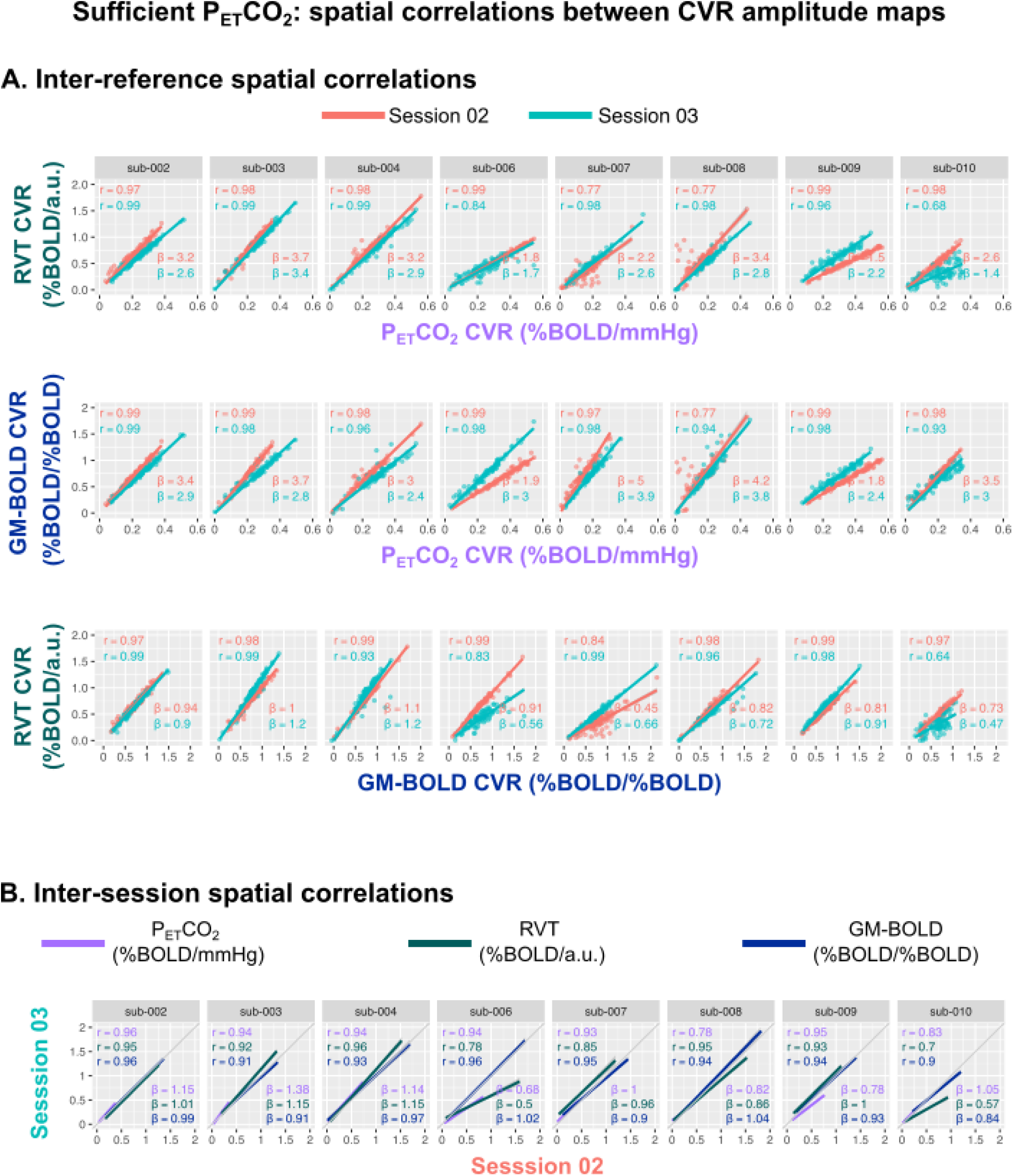
**A)** Inter-reference spatial correlations between P_ET_CO_2_, RVT, and GM-BOLD CVR amplitude maps, for each subject and session (summarized in Supplementary Table S4). Each of the three pairwise comparisons are plotted in a different row. **B)** Inter-session spatial correlations between CVR amplitude maps from the same reference signal between two consecutive sessions (summarized in Supplementary Table S5). The unity line (y=x) is plotted in gray for reference. All **c**orrelations were computed using the median CVR amplitude in 96 cortical parcels, identified from the Harvard-Oxford cortical atlas and separated by hemisphere. Each dot in a sub-plot represents the median CVR amplitude in one cortical parcel. Lines-of-best-fit are shown between each pair of CVR amplitude maps. Pearson correlation coefficients (r) are listed in the top left corner and slopes for the lines-of-best-fit (β) are displayed in the bottom right corner.

There is variability in the slope of the relationship between CVR amplitudes, with the best reliability between P_ET_CO_2_ and RVT. In general, RVT and GM-BOLD CVR amplitudes are 2-3 times larger than for P_ET_CO_2_ (average slopes of 2.36±0.71 for P_ET_CO_2_ & RVT, 2.91±0.23 for P_ET_CO_2_ & GM-BOLD). However, the magnitudes may not be meaningful due to the arbitrary units in RVT and GM-BOLD CVR. The reliability of these slopes was assessed with an intraclass correlation (using a two-way random effects model of absolute agreement), with the following results: ICC(2,1)=0.62 for P_ET_CO_2_ & RVT, ICC(2,1)=0.44 for P_ET_CO_2_ and GM-BOLD, and ICC(2,1)=0.41 for GM-BOLD and RVT. Thus, there is good reliability for P_ET_CO_2_ & RVT CVR amplitudes, and fair reliability for the other inter-reference relationships. However, these estimates may be limited by the small number of repeated measurements and subjects.

#### 3.2.2 Inter-session comparisons

For all reference signals, the resulting CVR amplitude maps are highly similar between sessions, provided that there was sufficient P_ET_CO_2_ data in each subject. Inter-session spatial correlations were similar for each reference signal with no significant differences in average Fisher’s Z across subjects (P_ET_CO_2_: Z=1.62, RVT: Z=1.50, GM-BOLD: Z=1.75). The inter-session spatial correlations are depicted in Fig. 5B and summarized in Supplementary Table S5. There is nearly a 1:1 relationship in the CVR amplitude maps between consecutive sessions for each reference signal (average slopes: P_ET_CO_2_=1.00±0.23, RVT=0.91±0.26, GM-BOLD=0.95±0.07). Excluding the outlier of sub-006, the average slope for RVT increases to 0.97±0.21.

### 3.3 Sufficient P_ET_CO_2_ Datasets: CVR Delay Comparisons

#### 3.3.1 Inter-reference comparisons

The CVR delay maps generated by P_ET_CO_2_ and RVT reference signals show similar spatial variation, while GM-BOLD delay maps have smaller delay magnitudes and reduced contrast, among datasets with sufficient P_ET_CO_2_ quality. Fig. 6 displays CVR delay maps generated by each reference signal. Since CVR delay is expressed in quantitative units of seconds for all reference signals, CVR delay maps are centered around the GM median to fairly compare between reference signals. In general, P_ET_CO_2_ and RVT delay maps characterize more extreme relative delays than GM-BOLD delay maps (indicated by more yellow and violet voxels throughout P_ET_CO_2_ and RVT maps).

**Figure 6:**
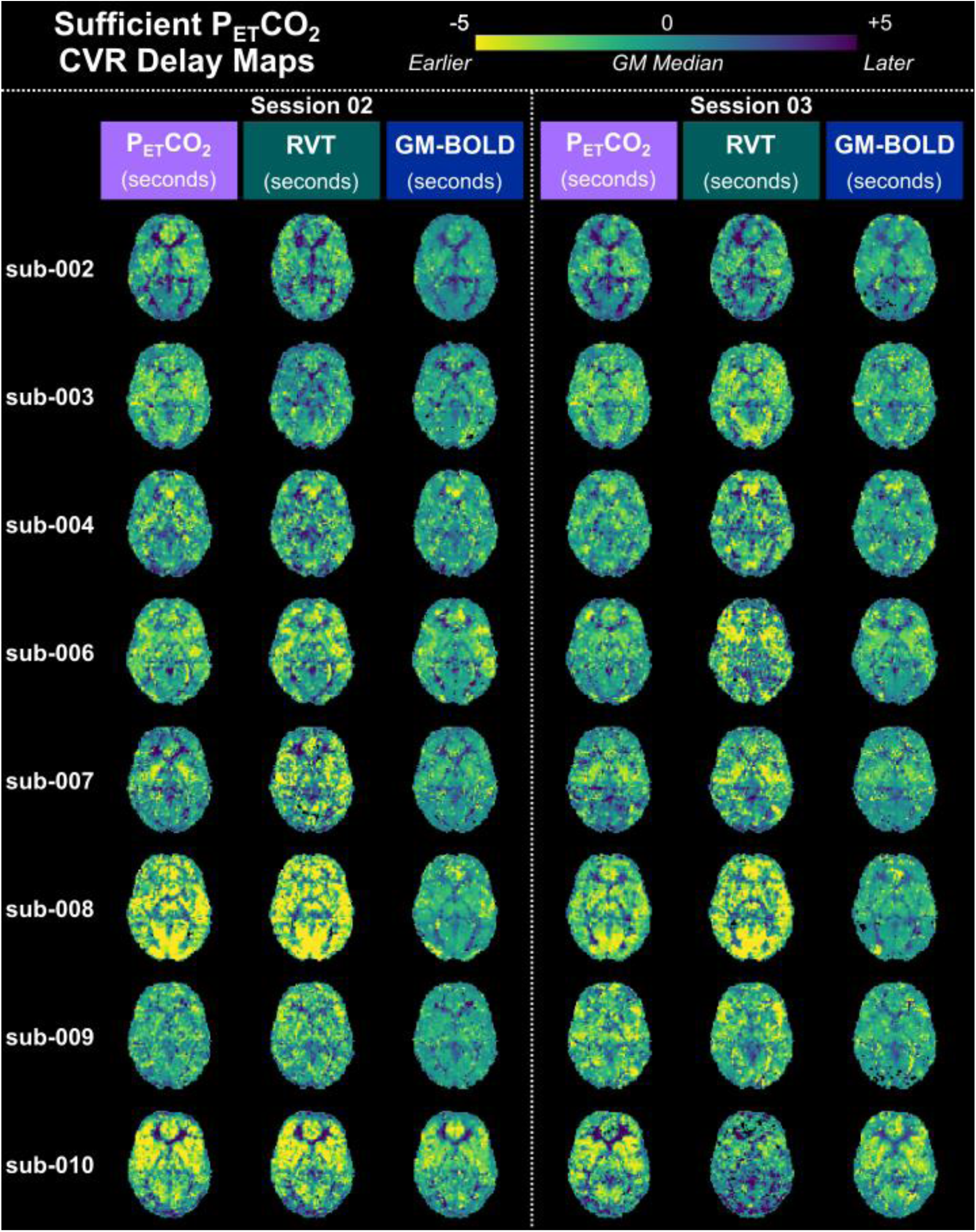
CVR delay maps for all datasets with sufficient P_ET_CO_2_ quality, transformed to the MNI152 6th generation template space. For each subject, maps from session 02 are shown on the left and maps from session 03 are shown on the right. A central, axial slice of the CVR delay map from each reference signal is shown in each column. CVR delay maps have been normalized to the median delay in grey matter (GM). Voxels at boundary conditions (absolute delay = +/- 8.7s, 9s) have also been removed. Negative values indicate regions with earlier hemodynamic responses relative to median delay in GM, while positive values indicate those with later responses.

The distributions of CVR delay for each reference signal (Fig. 7) support the observation that that P_ET_CO_2_ and RVT CVR delay maps show similar spatial variation while there is reduced contrast in GM-BOLD delay maps. The shape of P_ET_CO_2_ and RVT delay distributions are generally similar: both are slightly right skewed and centered just below 0 seconds. On the other hand, GM-BOLD delay distributions are narrower and zero-centered, with a high proportion of voxels exhibiting delay values near 0 seconds. In addition, the GM-BOLD delay distributions are less smooth, with several small peaks apparent for some datasets (e.g., sub-003 ses-02, indicated by the orange trace). Finally, P_ET_CO_2_ and RVT distributions are more variable between subjects, while GM-BOLD distributions have a relatively consistent shape.

**Figure 7:**
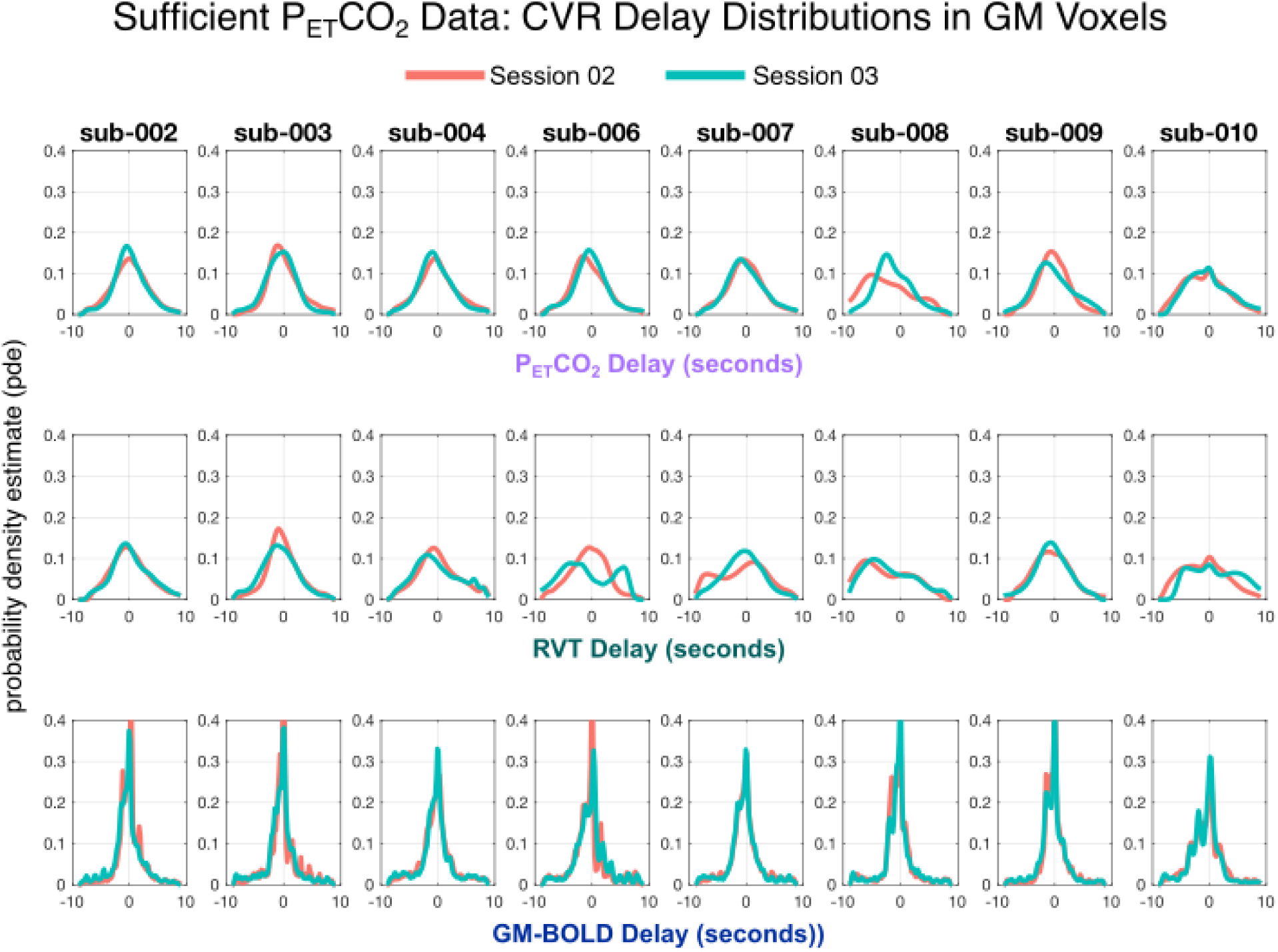
Distributions of CVR delay for each reference signal in all datasets with sufficient P_ET_CO_2_ quality. CVR delay values have been normalized to the median delay in grey matter. For each subject, distributions from session 02 are plotted in orange and session 03 are plotted in teal. Each row shows the distribution of CVR delay for a different reference signal, with P_ET_CO_2_ delay on top, RVT delay in the middle, and GM-BOLD delay on the bottom.

The slopes of inter-reference relationships (Fig. 8A) further illustrate the narrower range of delays observed with the GM-BOLD reference signal (Figs 6 and 7). P_ET_CO_2_ and RVT delay values are nearly proportional, with an average slope of 0.93±0.35. Excluding the outlier of sub- 002 ses-02, the average slope becomes 0.99±-0.06. However, as demonstrated in the maps, GM-BOLD delay values tend to underestimate delay relative to P_ET_CO_2_ and RVT. See middle and bottom rows of Fig. 8A, respectively, and note the switch in axes; this manifests as slopes >1 for GM-BOLD with P_ET_CO_2_ and slopes <1 for GM-BOLD with RVT.

**Figure 8:**
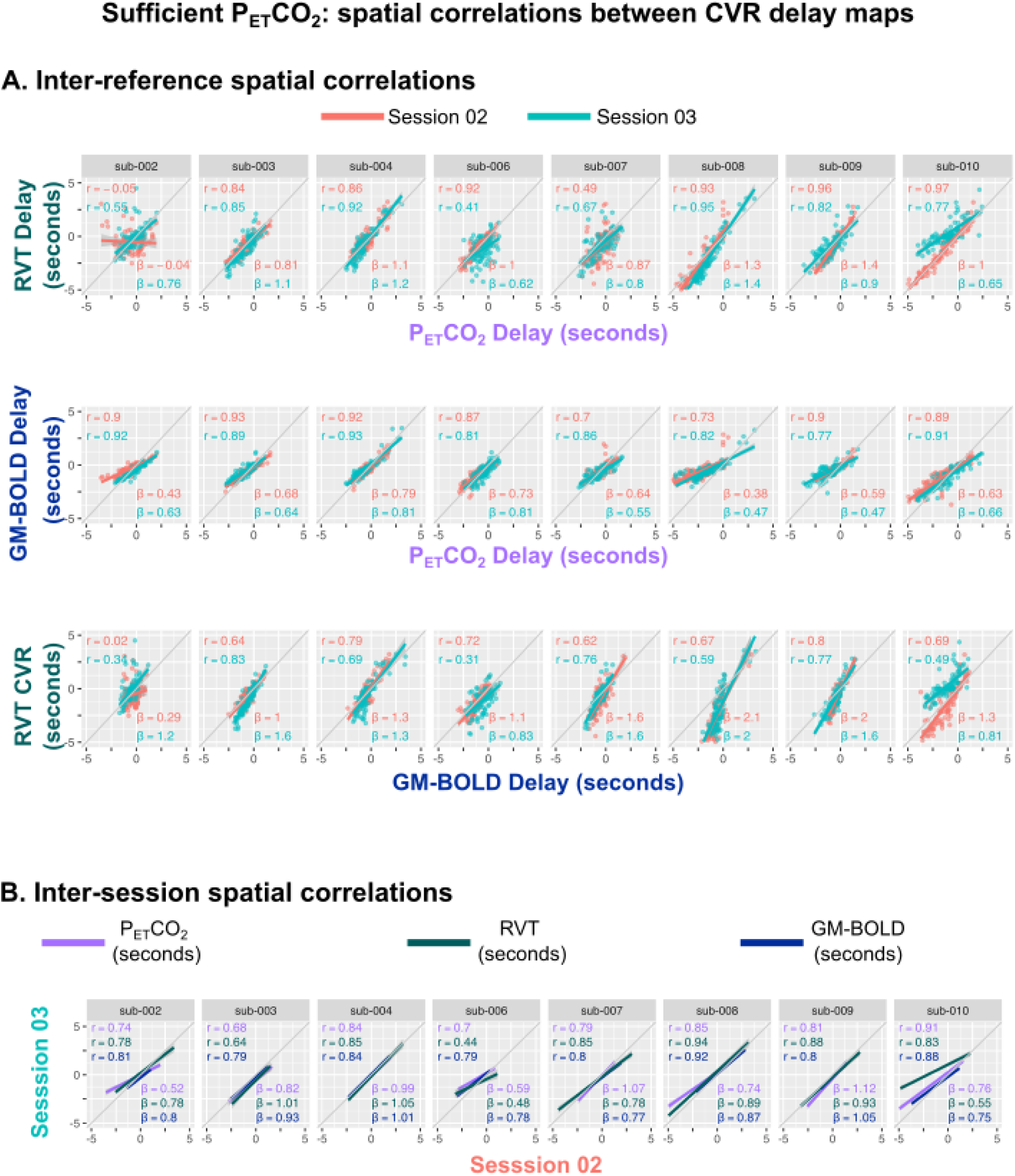
**A)** Inter-reference spatial correlations between P_ET_CO_2_, RVT, and GM-BOLD CVR delay maps, for each subject and session (summarized in Supplementary Table S6). Each of the three pairwise comparisons are plotted in a different row. The unity line (y=x) is plotted in gray for reference. **B)** Inter- session spatial correlations between CVR delay maps from the same reference signal between two consecutive sessions (summarized in Supplementary Table S7). All correlations were computed using the median CVR delay in 96 cortical parcels, identified from the Harvard-Oxford cortical atlas and separated by hemisphere. Each dot in a sub-plot represents the median CVR delay in one cortical parcel. Lines-of-best- fit are shown between each pair of CVR delay maps. Pearson correlation coefficients (r) are listed in the top left corner and slopes for the lines-of-best-fit (β) are displayed in the bottom right corner.

When comparing maps of CVR delay across the three reference signals (Fig. 8A), we see significant spatial correlations for all comparisons, although they are weaker on average compared to the spatial correlations of CVR amplitude (Fig. 5A). The corresponding spatial correlation coefficients, Fisher Z transformed correlations, and slopes for the lines-of-best fit are summarized in Supplementary Table S6. All group average inter-reference spatial correlations are significantly different from zero (P_ET_CO_2_ & RVT: Z=1.18, p<0.001; P_ET_CO_2_ & GM-BOLD: Z=1.35, p<0.001; GM-BOLD & RVT: Z=1.09, p<0.001). There were no significant differences in the average spatial correlations between each pair of reference signals, based on a t-test adjusted for non-independent correlations (Howell, 2010) (P_ET_CO_2_ & RVT vs. P_ET_CO_2_ & GM-BOLD: T(13)=0.59, p=0.57; P_ET_CO_2_ & GM-BOLD vs. GM-BOLD & RVT: T(13)=1.02, p=0.33; P_ET_CO_2_ & RVT vs. GM-BOLD & RVT: T(13)=0.43, p=0.67). However, there are two datasets with RVT delay values that are poorly correlated with other reference signals (sub-002 ses-02 and sub-006 ses- 03). These poor correlations are supported by low cross-correlations between the RVT and GM-BOLD reference signals in each dataset (0.46 and 0.64, respectively, Table S1), which may drive poor optimization in the lagged-GLM and cause more voxels to be near boundary conditions. Local variations in BOLD signal features may introduce regional differences in the success of optimization, overall leading to poor spatial agreement between RVT delay maps with other reference signals. This is consistent with the CVR delay maps shown in Fig. 6 for these datasets.

The slope of the relationship between CVR delay values from a given pair of reference signals is generally more consistent compared to CVR amplitudes, consistent with the common quantitative units (seconds) of CVR delay achieved with all three methods. This is demonstrated by the inter-subject consistency of slopes for each best-fit line in Fig. 8A. In addition, the inter-session reliability of the slopes was assessed with an intraclass correlation (using a two-way random effects model of absolute agreement). There is good reliability between P_ET_CO_2_ and GM-BOLD delays (ICC(2,1)=0.73) and between GM-BOLD and RVT delays (ICC(2,1)=0.63). However, there is poor reliability between P_ET_CO_2_ and RVT delays (ICC(2,1)=0.35). As with CVR amplitude, these ICC estimates may be limited by the small number of repeated measurements and subjects.

#### 3.3.2 Inter-session comparisons

CVR delay maps for each reference signal are also highly spatially correlated between two consecutive sessions, provided the P_ET_CO_2_ quality was sufficient. These inter-session spatial correlations for CVR delay are summarized in Fig. 8B and Supplementary Table S7. There were no significant differences in average Fisher’s Z across subjects (P_ET_CO_2_: Z=1.11, RVT: Z=1.14, GM-BOLD: Z=1.21). The average slope between delays from consecutive sessions is also similar for each reference signal (average slope for P_ET_CO_2_: 0.83±0.22, RVT: 0.81±0.21, GM-BOLD: 0.87±0.11).

### 3.4 Insufficient P_ET_CO_2_ Datasets

#### 3.4.1 Inter-quality comparisons

As described in Section 3.1, a total of 6 datasets were identified as having insufficient P_ET_CO_2_ quality, based on the relative power content at the BH task frequency. Fig. 9 shows the reference signals, power spectra, and resulting CVR maps from datasets with sufficient and insufficien t P_ET_CO_2_ quality within the same example subject (inter-quality comparison). Not surprisingly, the CVR amplitude and delay maps generated by an insufficient P_ET_CO_2_ timeseries do not show physiologically plausible spatial variations (Fig. 9B). Despite the insufficient task-related information within the P_ET_CO_2_ timeseries, the RVT and GM-BOLD timeseries still demonstrate modulations consistent with the 8 cycles of the BH task and clear peaks in their power spectra. Therefore, consistent with our hypothesis, the resulting RVT and GM-BOLD CVR parameter maps are comparable to those from the dataset with sufficient P_ET_CO_2_ quality.

**Figure 9:**
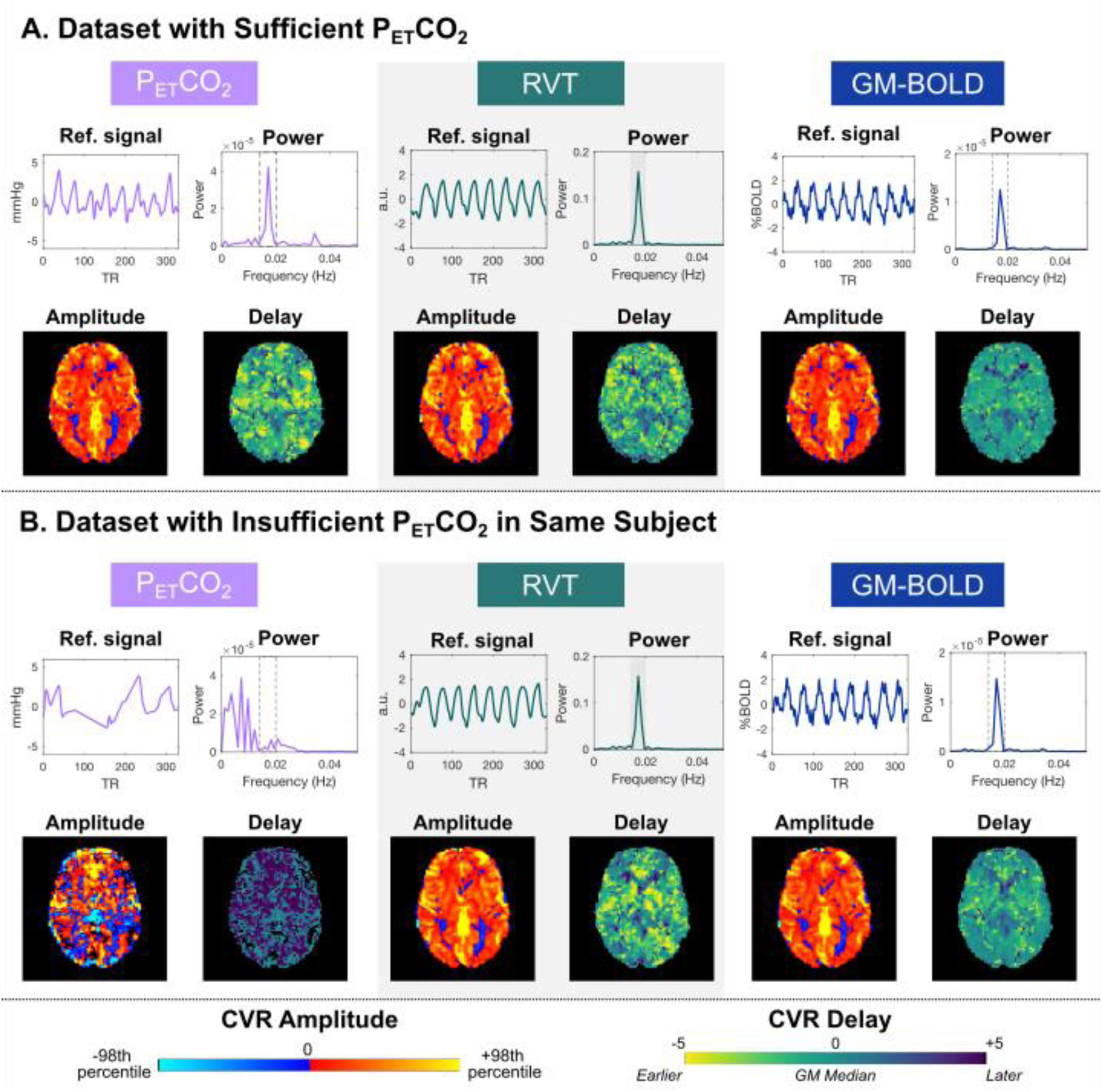
Example reference signals, power spectra, and CVR maps for two datasets in the same subject (sub-009) with A) sufficient P_ET_CO_2_ quality (ses-02) and B) insufficient P_ET_CO_2_ quality (ses-08). The insufficient P_ET_CO_2_ timeseries can be distinguished by the absence of a peak in the power spectrum at the breath-hold task frequency (0.014 to 0.020 Hz, indicated by dashed lines). CVR amplitude and delay maps are comparable between the two datasets, for all reference signals except insufficient P_ET_CO_2_. Note that the RVT timeseries and power spectra are plotted on different scales for visualization purposes. CVR maps are scaled to 98^th^ percentile values, which can be found in Table S2. Also note that only the P_ET_CO_2_ CVR amplitude map is in quantitative units (%BOLD/mmHg), compared to RVT CVR (%BOLD/a.u.) and GM-BOLD CVR (%BOLD/%BOLD).

The inter-quality spatial correlations between CVR parameter maps from insufficient and sufficient quality datasets support the qualitative observations in Fig. 9, in that the maps generated by RVT and GM-BOLD timeseries recover spatial information that is lost by those from the insufficient P_ET_CO_2_ trace. For each reference signal’s CVR map from an insufficient P_ET_CO_2_ session, a spatial correlation was performed with the respective parameter map from the first sufficient P_ET_CO_2_ session (ses-02) in the same subject. Table 4 summarizes the Fisher’s Z transformed spatial correlation coefficients and the slope of the best-fit line between these data. When the reference signal is “insufficient P_ET_CO_2_”, the average spatial correlations with a map computed using sufficient P_ET_CO_2_ data acquired in a different scan session are not significant for either CVR amplitude (Z=1.11±0.53) or CVR delay (Z=0.25±0.53), using Z_crit_=1.13 for N=6 at alpha=0.05. In contrast, the inter-session spatial correlations for RVT and GM-BOLD CVR amplitude and delay maps are significant between sufficient and insufficient datasets. This is expected, since the categorization of “sufficient” datasets was based on P_ET_CO_2_ quality, with RVT and GM-BOLD signals surpassing the relative power criterion in all datasets.

**Table 4:** Inter-quality spatial correlations between each reference signal’s CVR map from an insufficient P_ET_CO_2_ quality dataset and the corresponding CVR map from a sufficient P_ET_CO_2_ dataset

| Subject | Insufficient Session | Sufficient Session | Inter-Quality Spatial Correlations |  |  |  |  |  |  |  |  |  |  |  |
| --- | --- | --- | --- | --- | --- | --- | --- | --- | --- | --- | --- | --- | --- | --- |
|  |  |  | CVR Amplitude |  |  |  |  |  | CVR Delay |  |  |  |  |  |
| | | | $P_{ET}CO_2$ | | RVT | | GM-BOLD | | $P_{ET}CO_2$ | | RVT | | GM-BOLD | |
| | | | $\beta$ | Z | $\beta$ | Z | $\beta$ | Z | $\beta$ | Z | $\beta$ | Z | $\beta$ | Z |
| sub-006 | ses-07 | ses-02 | 1.30 | 1.73 | 1.18 | 1.83 | 1.00 | 1.88 | 0.43 | 0.54 | 0.67 | 0.94 | 0.60 | 0.90 |
|  | ses-08 | ses-02 | 0.98 | 1.22 | 1.04 | 1.55 | 1.01 | 1.70 | 0.44 | 0.46 | 0.62 | 1.07 | 0.77 | 1.23 |
| sub-009 | ses-08 | ses-02 | 0.06 | 0.26 | 1.02 | 1.64 | 1.01 | 1.76 | -0.82 | 0.56 | 0.79 | 1.29 | 1.09 | 1.42 |
|  | ses-09 | ses-02 | 0.55 | 1.42 | 0.85 | 1.70 | 0.98 | 1.75 | 0.51 | 0.78 | 0.79 | 1.02 | 0.54 | 0.95 |
| sub-010 | ses-07 | ses-02 | 0.98 | 0.74 | 1.05 | 1.44 | 0.93 | 1.66 | 0.29 | 0.71 | 0.75 | 1.49 | 0.79 | 1.20 |
|  | ses-08 | ses-02 | 1.15 | 1.30 | 1.12 | 1.38 | 1.19 | 1.67 | 0.63 | 1.17 | 0.68 | 1.25 | 0.81 | 1.59 |
| Average |  |  | 0.84 | 1.11 | 1.04 | 1.59* | 1.02 | 1.74* | 0.25 | 0.70 | 0.72 | 1.18* | 0.77 | 1.21* |
| StDev |  |  | 0.46 | 0.53 | 0.11 | 0.17 | 0.09 | 0.08 | 0.53 | 0.26 | 0.07 | 0.21 | 0.19 | 0.26 |

$\beta$ = coefficient of slope for best-fit line to correlation. Z = Fisher's Z transformation of correlation coefficient.
\*Indicates Fisher's Z is significantly different from 0 at $\alpha = 0.05$ (critical Z = 1.13 for N = 6).

However, it is important to note the differences in P_ET_CO_2_ CVR maps are not as dramatic for all datasets with insufficient P_ET_CO_2_ quality. These maps are presented in Supplementary Figure S3. Specifically, amplitude maps from some insufficient P_ET_CO_2_ traces have reasonable quality, while the delay maps remain noisy. For example, the CVR amplitude maps obtained with insufficient P_ET_CO_2_ are similar to those obtained with RVT and GM-BOLD for sub-006 ses-07, sub-009 ses-09, and sub-010 ses-08. These datasets also have higher inter-quality spatial correlations, as indicated by the Fisher’s Z values in Table 4. The relative power in the insufficient P_ET_CO_2_ signals for these three datasets (Table 2) far exceeds the relative power of 4.33% in the example case highlighted in Fig. 9, indicating that there may have been some sufficient BH trials to generate reasonably good CVR amplitude maps. While some insufficient P_ET_CO_2_ CVR amplitude maps are similar, the CVR delay maps still have noticeable regional differences (e.g., more negative delays and reduced tissue contrast), though less extreme than shown in Fig. 9.

#### 3.4.2 Inter-reference comparisons

Similarly, the inter-reference spatial correlations within each insufficient P_ET_CO_2_ dataset demonstrate the corrupted CVR amplitude and CVR delay maps generated by the P_ET_CO_2_ traces. These results can be found in Supplementary Table S9. Correlations of P_ET_CO_2_ CVR amplitude with RVT and GM-BOLD CVR amplitude are expectedly lower (Z=1.44±0.73 and Z=1.41±0.69, respectively) compared to those between GM-BOLD and RVT (Z=2.56±0.13), which still have sufficient power at the task frequency. This difference is especially apparent in sub-009 ses-08 and sub-010 ses-07.

The same pattern of low spatial correlations with results derived from P_ET_CO_2_ is evident in the CVR delay values (Z=0.75±0.38 for correlation of delays with insufficient P_ET_CO_2_ CVR and RVT; Z=0.66±0.37 with insufficient P_ET_CO_2_ CVR and GM-BOLD, and Z=1.340±0.22 with RVT and GM-BOLD). In addition, the average spatial correlations for CVR delay are all considerably lower than those for CVR amplitude of the same pairwise comparison.

## 4. Discussion

In this study, we tested whether RVT or GM-BOLD can be used in a lagged-GLM framework to achieve estimates of CVR amplitude and delay that are spatially correlated with estimates from P_ET_CO_2_. We tested this in breath-hold data in healthy adults, including datasets where P_ET_CO_2_, RVT, and GM-BOLD reference signals had sufficient power (>50%) at the task frequency, and datasets where only the P_ET_CO_2_ timeseries had insufficient power. We found that in datasets with sufficient quality, all reference signals are highly correlated. Correspondingly, CVR amplitude maps are spatially similar for all reference signals, after accounting for differences in scale . However, both RVT and GM-BOLD CVR amplitudes are not in standard CVR units of %BOLD/mmHg. Regarding CVR delay, the maps generated by P_ET_CO_2_ and RVT show similar spatial variation, while GM-BOLD delay maps have a smaller range and reduced contrast. Finally, when P_ET_CO_2_ is insufficient, RVT and GM-BOLD can be used to recover spatially similar CVR amplitude and delay maps, provided that the participant attempted the breath-hold task. We explore each of these findings in further detail in the following sections.

### 4.1 Reference signals are highly correlated in breath-hold data with sufficient P_ET_CO_2_ quality

The high cross-correlation amplitudes observed between P_ET_CO_2_, RVT, and GM-BOLD signals are expected and consistent with previous reports in the literature. Each of these signals captures the physiological processes occurring during a breath-hold, marked by a cessation of breathing, increased arterial CO_2_ concentration, increased CBF, and an increased BOLD signal that eventually returns to baseline (Bright et al., 2009; Kastrup et al., 1999; Thomason et al., 2005). P_ET_CO_2_ and RVT have separately been shown to correlate with the resting-state BOLD timeseries (Birn et al., 2008, 2006; Wise et al., 2004). Additionally, Chang et al. demonstrated that P_ET_CO_2_ and RVT (convolved with the respiration response function) are highly correlated and account for similar spatial and temporal variations in the resting-state BOLD signal (Chang and Glover, 2009).

In breath-hold data, these cross-correlations are magnified due to the alternating periods of task and rest, which lead to large coupled amplitude fluctuations in P_ET_CO_2_, RVT, and GM-BOLD that are approximately sinusoidal at the task frequency (Pinto et al., 2021). These quasi-sinusoidal variations are critical to our approach for determining sufficient P_ET_CO_2_ based on relative power at the task frequency. While this strategy can be easily implemented to quality check P_ET_CO_2_ recordings, it requires periodic breathing modulation and thus cannot easily be translated to evaluate the quality of natural P_ET_CO_2_ fluctuations in resting-state data.

The reference signals we considered are not exhaustive. The near-sinusoidal fluctuations in the BOLD response during a quasi-periodic breath-hold task can be modeled using a Fourier series, with a sine-cosine pair at the task frequency and additional harmonics, to estimate both CVR amplitude and delay (Lipp et al., 2015; Murphy et al., 2011; Pinto et al., 2016; van Niftrik et al., 2016). Additionally, many studies use different variations of a global BOLD signal to model CVR, rather than a respiratory-derived signal, due to the known influence of arterial CO_2_ fluctuations on the BOLD signal (Geranmayeh et al., 2015; Liu et al., 2017; Tong et al., 2011; Tong and Frederick, 2014; van Niftrik et al., 2016). As we have demonstrated with GM-BOLD, there are clear breath-hold effects in the average BOLD response, leading to CVR measurements that are comparable to those derived from P_ET_CO_2_.

### 4.2 CVR amplitude maps are comparable between reference signals, but RVT and GM-BOLD amplitudes are not in standard CVR units

Based on the high cross-correlations between input reference signals, it is not surprising that the resulting CVR amplitude maps are also highly correlated. In fact, CVR maps from each reference signal look nearly identical when scaled to the 98^th^ percentile CVR amplitude. Regardless of the method used to model CVR, this visualization scaling approach may facilitate qualitative comparisons of CVR maps, longitudinally, between cohorts, and between protocols. Our CVR visualization approach also indicates the method used to model CVR may not be critical for qualitative comparisons, which is consistent with the current ethos regarding the “multiverse” of analysis pipelines in the functional neuroimaging community (Botvinik-Nezer et al., 2020; Dafflon et al., 2022; Steegen et al., 2016; Taylor et al., 2022).

Despite the qualitative similarities between CVR maps, there are important differences in the absolute magnitudes of CVR amplitude. Both RVT and GM-BOLD CVR are not in standard CVR units, which is an important caveat, particularly for comparing CVR between cohorts or with literature values. In these cases, it is still best to use P_ET_CO_2_ as a reference signal, because the resulting CVR amplitude in units of %BOLD/mmHg is physiologically meaningful. There is also between-subject variability in the slope of the relationship between CVR amplitudes, likely driven in part by the arbitrary units of RVT and GM-BOLD CVR. Overall, RVT CVR had the most reliable relationship with P_ET_CO_2_ CVR amplitude, suggesting that this might be a better alternative than GM-BOLD to capture differences in CVR amplitude.

However, our results indicate that RVT and GM-BOLD would still be useful in many cases, such as making relative comparisons between brain regions within a subject and identifying focal pathology. In addition, the CVR maps for RVT and GM-BOLD were consistent between scan sessions. With these steady measurements, it could be possible to compare longitudinally within a subject, provided that a breath-hold task is used to induce modulations and there is sufficient power at the task frequency.

We observed that normalizing the RVT timeseries before inputting it to the lagged-GLM is critical to achieve reasonable CVR amplitude values. RVT is reported in arbitrary units (a.u.) because the magnitude of RVT varies across experimental setups and is sensitive to the tightness of the respiration belt and its placement on the body (i.e., chest vs. abdomen). Thus, there is high variability in the scale of RVT fluctuations across datasets (Supplementary Fig. S1 and Table S2). The resulting CVR amplitude maps are impacted by this variability because they are scaled to the amplitude of the reference signal. If RVT is not normalized, there are large differences in the range of amplitude values, which could be misleading if CVR maps are plotted on a fixed scale (Supplementary Fig. S2).

### 4.3 CVR delay maps are comparable for P_ET_CO_2_ and RVT, but GM-BOLD may underestimate delay variability

In datasets with sufficient P_ET_CO_2_ quality, RVT and GM-BOLD both produce delay maps that are highly correlated with those from P_ET_CO_2_, but the delay magnitudes tend to be smaller when GM-BOLD is the reference signal. This is evident in the narrower distributions of GM-BOLD delay values (Fig. 7) and in the biased slopes from inter-reference spatial correlations with P_ET_CO_2_ and RVT delays (Fig. 8A). Thus, GM-BOLD may underestimate the true delay value, particularly for voxels with larger absolute P_ET_CO_2_ delays.

There are a few potential reasons for the narrower distribution of GM-BOLD delay values. We normalized delay maps to the GM median to compare between reference signals and participants. Many GM voxels will be well-characterized by the average BOLD timeseries and have similar delay values that are reduced to zero after this spatial normalization step. Additionally, the GM-BOLD signal (after the T2-weighted combination of the echoes) might be more affected by motion-related effects than other reference signals (Moia et al., 2021). For example, peaks or slow drifts in the GM-BOLD timeseries due to head motion could bias the optimum delay estimated for a given voxel. More likely, the GM-BOLD signal is “blurring” the breath-hold response due to the wide variation in relative timing across the brain (Tong et al., 2019). This has been addressed previously with the concept of making a “refined” or “dynamic” global signal regressor that accounts for voxel-specific variations in delay to recover a source signal (Erdoğan et al., 2016; Frederick et al., 2012; Tong and Frederick, 2014). Our approach using the average response across GM voxels is well-established but more simplistic and may have restricted the sensitivity to a wider range of delays. An average signal from the cerebellum (Donahue et al., 2016; Liu et al., 2021), sagittal sinus (Pillai and Mikulis, 2015; van Niftrik et al., 2016), or other small ROIs (Erdoğan et al., 2016) could also be used to mitigate this issue. However, the cerebellum is sensitive to noise (Diedrichsen et al., 2010; van der Zwaag et al., 2015) and these ROIs are arbitrary for CVR analysis.

To address limitations attributed to the GM-BOLD regressor, we performed a post hoc exploratory analysis to compare CVR delays using a “refined” GM-BOLD approach. The refined GM-BOLD regressor used in this analysis was generated by *Rapidtide* v2.2.7, a data-driven algorithm that uses the refined GM-BOLD timeseries as a regressor, for which it iteratively considers a voxel-by-voxel fit across a range of temporal offsets using a cross-correlation method (Frederick et al., 2012, 2016). We considered a temporal range of ±9 s with 0.3 s increments to match the lagged-GLM (specific command options are detailed in Table S11, and we refer the reader to the *Rapidtide* documentation (Frederick et al., 2022a) to explore more in-depth details about the settings). This algorithm further differs from the lagged-GLM processing method by also temporally smoothing the average GM-BOLD response with a band-pass filter (0.009-0.15 Hz) and “despeckling” using a spatial median filter to correct erroneous time delays due to autocorrelation in the probe regressor (Frederick, 2017). Additionally, motion parameters and Legendre polynomials are regressed from the data before the cross-correlation fit, in contrast to being included in the lagged-GLM. Fig. 10 shows a comparison between the original GM-BOLD approach and a refined GM-BOLD approach for a representative subject (sub-008). Results for all subjects with sufficient P_ET_CO_2_ data quality can be found in Supplementary (Figures S4-S7, Table S10). The refined GM-BOLD regressor is similar to the GM-BOLD time series yet smoother, with high frequencies removed (Fig. 10A).

**Figure 10:**
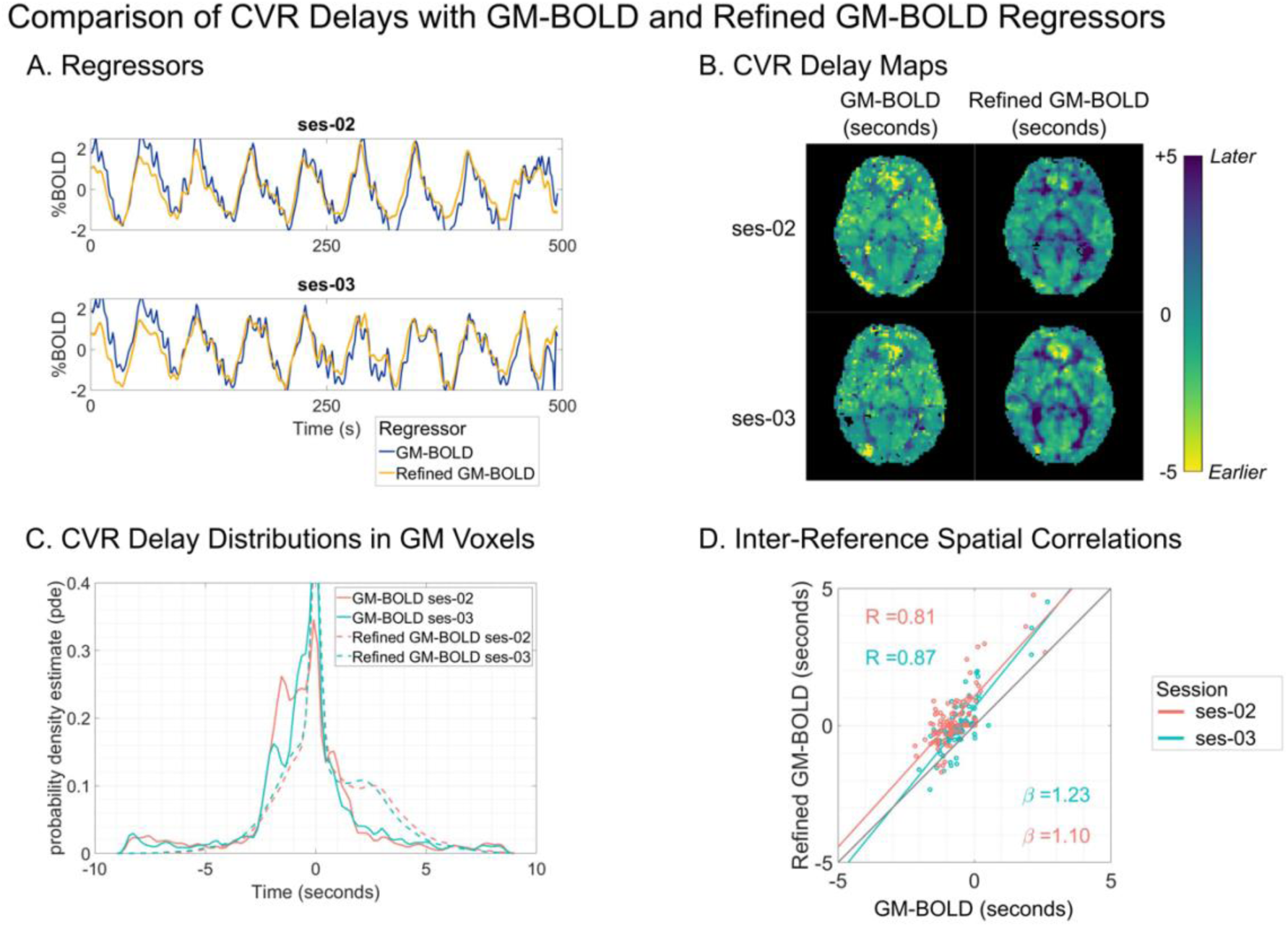
**A)** Reference signal for sub-008 from GM-BOLD (blue) and Refined GM-BOLD (yellow) in ses- 02 (top) and ses-03 (bottom). **B)** CVR delay map for sub-008, transformed to MNI space. Maps from ses- 02 are shown on the top row, and maps from ses-03 are shown on the bottom row. An axial slice from two compared methods is shown in each column: GM-BOLD (left) and Refined GM-BOLD (right). CVR delay maps using the GM-BOLD approach have been normalized to the GM median delay with voxels at boundary conditions (absolute delay = +/- 8.7s, 9s) removed. Refined GM-BOLD delay maps are re-centered to 0s and exclude voxels where the similarity function failed (Frederick et al., 2022a). **C)** CVR delay distribution for sub-008 across GM-BOLD (solid line) and Refined-GM (dashed line) from ses-02 (orange) and ses-03 (teal). **D)** Inter-reference spatial correlation between GM-BOLD and Refined GM-BOLD delay maps for sub-008 in ses-02 (orange) and ses-03 (teal) with respective best-fit-lines and an identity line (black) for comparison. Each point represents the median delay value in one of the 96 cortical parcels from the Harvard-Oxford cortical atlas. Correlation coefficient (R) for each session is listed on the top left, and the slopes for the lines-of-best-fit (/beta) for each session are listed on the bottom right.

CVR delay maps generated using the refined GM-BOLD approach depict greater visual contrast between gray matter and white matter in comparison to the CVR delay maps generated with the average GM-BOLD approach (Fig. 10B). Furthermore, the distribution of delays generated from a CVR delay map using the refined GM-BOLD approach show a skewness towards larger positive delays (Fig. 10C). The delay values from both methods are highly correlated and the slopes of the spatial correlations are greater than 1, indicating that the refined GM-BOLD approach depicts more extreme delays across most of the cortex in comparison to the GM-BOLD approach (Fig. 10D). Thus, using a refined GM-BOLD timeseries as a regressor may partially compensate for the smaller distribution in delays attributed to the lagged-GLM with a standard GM-BOLD timeseries.

### 4.4 When P_ET_CO_2_ quality is insufficient, maps of CVR amplitude and delay can be achieved with RVT or GM-BOLD as reference signals

We have demonstrated that in breath-hold fMRI data, if a participant attempts the task but P_ET_CO_2_ quality is poor, RVT or GM-BOLD can be used to create CVR amplitude and delay maps. Based on comparisons with sufficient P_ET_CO_2_ quality data, RVT seems the best alternative to generate CVR amplitude and delay maps that are highly correlated and have consistent relationships with those obtained with P_ET_CO_2_ measurements. In addition, RVT still generates CVR measurements that are normalized to a respiratory-derived measure. If opting for a global signal like GM-BOLD, it would be best to use a refined GM-BOLD regressor to account for potential under-estimation of CVR delay.

We also proposed a method to define a “sufficient” P_ET_CO_2_ trace for CVR mapping, using a relative power threshold >50% at the breath-hold task frequency. However, this threshold is slightly arbitrary and may need to be adjusted for specific cases, with a holistic evaluation of reference signals and their resulting CVR amplitude and delay maps. In fact, some of the datasets with insufficient P_ET_CO_2_ still showed reasonably good CVR amplitude maps (Fig. S3). However, the corresponding CVR delay maps are less similar to those generated by sufficient quality timeseries and should give cause for caution when interpreting the CVR amplitude maps, due to potential mis-fitting of the reference signal. For example, there are several regions of negative CVR amplitudes in the map for sub-010 ses-07 (indicated by blue voxels in the corresponding map of Fig.S3), which resemble the vascular “steal” phenomenon and could be mis-characterized as pathology (Conklin et al., 2010; Poublanc et al., 2013; Sam et al., 2016). Therefore, insufficient P_ET_CO_2_ CVR maps should be interpreted carefully, particularly in clinical cases.

Although these are promising results to recover CVR maps retrospectively or in low resource settings, we still recommend trying to obtain sufficient P_ET_CO_2_ estimates from a breathing modulation for the highest quality CVR maps. There are suggestions throughout the literature on how to implement robust breath-hold tasks (Bright and Murphy, 2013; Murphy et al., 2011; Pinto et al., 2021; Scouten and Schwarzbauer, 2008; Urback et al., 2017). In brief, it is strongly recommended to incorporate a training session before the scan to ensure that participants understand and comply with task instructions (Kannurpatti et al., 2010; Magon et al., 2009; Zacà et al., 2014). Monitoring respiratory signals throughout the task is also encouraged to ensure quality of the recording and assess task performance (Bulte and Wartolowska, 2017). In addition, cueing strategies (e.g., text, symbolic, or auditory) should be carefully considered to make instructions intuitive for the target population. Lastly, other breathing tasks might be more feasible than a breath-hold, such as intermittent breath modulation (Liu et al., 2020) or paced deep breathing (Bright et al., 2009; Sousa et al., 2014; Stickland et al., 2021). With these alternative methods, a similar approach to determine relative power at the task frequency could still be implemented, though the limitations of extending our findings to other breathing modulations are discussed in *Section 4.7*.

### 4.5 Potential impacts and examples of utility

The use of alternative reference signals to generate CVR amplitude and delay maps has a range of potential impacts. The framework proposed here using RVT or GM-BOLD reference signals makes prospective CVR mapping accessible to any imaging centers that lack the equipment and personnel necessary to monitor and post-process respiratory gas recordings. A respiration belt should be integrated with most scanning set-ups, and the GM-BOLD signal requires no additional monitoring. All lagged-GLM regression analyses, with the exception of the RVT computation, are based on open-source software (peakdet, phys2cvr, and rapidtide) to facilitate the modeling steps for future applications. In addition, these findings present the opportunity to retrospectively generate CVR maps in breath-hold data where P_ET_CO_2_ data was not collected or had insufficient quality.

Potentially most impactful, a method to acquire robust CVR amplitude and delay maps even in datasets with insufficient P_ET_CO_2_ quality has important implications for populations where it may be difficult to obtain reliable end-tidal measurements. This includes children, where previous work has demonstrated reasonable task compliance but poor P_ET_CO_2_ quality, either due to mouth breathing or failure to perform end-exhales. It also includes aging cohorts and clinical populations (both pediatric and adults), who may similarly have difficulty following the steps needed for sufficient quality P_ET_CO_2_ (Handwerker et al., 2007; Thomason et al., 2005).

Overall, improved accessibility to CVR mapping can increase the prevalence of this informative metric of vascular health. Several reviews have described the utility of CVR mapping for understanding disease mechanisms and as a biomarker to triage patients for therapeutic interventions and track the efficacy of these interventions (Blair et al., 2015; Gupta et al., 2012; Juttukonda and Donahue, 2019; Pillai and Mikulis, 2015; Sleight et al., 2021; Smeeing et al., 2016). Aside from clinical populations, CVR mapping is also recommended in healthy cohorts to isolate differences in the BOLD response that may be due to differences in vascular rather than neural processes (Handwerker et al., 2007; Thomason et al., 2007; Tsvetanov et al., 2015).

### 4.6 Alternative approaches to address compliance challenges in CVR mapping

While we ideally recommend using a breathing task and respiratory-derived signal for CVR mapping, alternative methods without end-tidal CO_2_ recordings or in resting-state have been proposed to address the challenges associated with breathing tasks. These methods are reviewed in detail by Pinto et al., 2021, but we discuss key comparisons. As described in *Section 4.4*, the *Rapidtide* algorithm can generate a probe regressor from the global BOLD signal or another reference timeseries and use temporal cross-correlation with each voxel timeseries to determine maximum correlations and corresponding time delays (Frederick et al., 2016) (Tong and Frederick, 2014). The correlation metrics are surrogates for CVR, although the outputs are not in the standard CVR units (%BOLD/mmHg) that allow for comparison across subjects, particularly if the global signal is used as the probe. With a P_ET_CO_2_ probe, the *Rapidtide* outputs could potentially be modified to obtain CVR amplitude in normalized units.

From resting-state data, the global BOLD signal can be bandpass filtered to approximate arterial CO_2_ fluctuations and used as a regressor to estimate CVR (Liu et al., 2017). In addition, resting-state metrics such as the amplitude of low frequency fluctuations (ALFF) or fractional ALFF (fALFF) have demonstrated high correlations with CVR derived from CO_2_ challenges (Di et al., 2013; Golestani et al., 2016; Kazan et al., 2016). However, this relationship is controversial (Moia et al., 2022a). For instance, Moia et al., 2022 shows, in the same dataset used in this study, that resting-state metrics (RSFA, ALFF, fALFF) have highly variable inter-subject relationships with breath-hold CVR measures.

Additionally, the lagged-GLM has been previously performed with P_ET_CO_2_ from resting-state data, but the delay optimization procedure was less successful, leading to noisy estimates of CVR amplitude and delay (Stickland et al., 2021). This was hypothesized to be due to the smaller fluctuations in the resting-state signal relative to a breathing task and the confounds of low-frequency oscillations from neural activity and other physiological processes that may disrupt the optimization procedure (Caballero-Gaudes and Reynolds, 2017; Liu, 2016; Murphy et al., 2013). The increased BOLD sensitivity and data quality associated with the multi-echo acquisition in our study also helps to improve the CVR estimates (Moia et al., 2021). Similar results would be expected for alternative resting-state reference signals.

Although these alternative methods provide some insight into cerebrovascular physiology, each is missing a key characteristic of robust CVR measurements. Namely, none of these methods simultaneously use a breathing modulation to challenge the vascular system, normalize the BOLD changes to a reference signal that accounts for variability in breathing task performance, and correct for regional delays in CVR response time (Stickland et al., 2021).

### 4.7 Limitations and future work

The comparisons laid out in this study are valid only for breath-hold task fMRI data in healthy individuals. The alternative reference signals considered here may not be highly correlated with P_ET_CO_2_ in paced deep breathing tasks and may have insufficient variability in resting state to produce reasonable CVR maps. Using a paced hyperventilation task to induce hypocapnia, Vogt and colleagues (2011) found that RVT convolved with the respiration response function was less strongly correlated to BOLD signal changes than P_ET_CO_2_ convolved with an empirically derived response function. They conjectured that the uncoupling of the signals was due to the higher rate of CO_2_ reduction during hyperventilation relative to the slower rate of metabolic CO_2_ production, which is captured by the P_ET_CO_2_ regressor but not in the canonical respiration response (Vogt et al., 2011). This could potentially be addressed with optimization of the respiration response function for hypocapnia.

Furthermore, to achieve RVT or GM-BOLD CVR maps that are comparable to those from P_ET_CO_2_, the participant must attempt the breath-hold task with repeated periods of apnea. These changes in chest position are necessary to generate an RVT signal that has sufficient power at the task frequency. Similarly, the periods of apnea are required to induce a rise in arterial CO_2_ levels and the successive increase in CBF detected by the GM-BOLD signal. Achieving this level of task compliance could still be difficult in some cohorts, although there is a breadth of literature demonstrating successful use of breath-hold tasks (Pinto et al., 2021; Urback et al., 2017).

In participants with cerebrovascular pathology, careful consideration should be given to the reference signal and lagged-GLM parameters. For example, the average gray matter signal might be biased by regions with atypical perfusion dynamics. This could be addressed by averaging across normal-appearing tissue, or by using global signal refinement procedures as described in *Section 4.4*. Hemodynamic delays are also longer in many pathologies, such as steno-occlusive, small vessel disease, and dementia (Atwi et al., 2019; Duffin et al., 2015; Hartkamp et al., 2012; Holmes et al., 2020; McKetton et al., 2019; Thrippleton et al., 2018). The delay range used in the lagged-GLM step should be extended to reflect those that are physiologically plausible for a given condition. Using a lagged-GLM approach, delays in the range of ±9 seconds are consistently reported in healthy individuals (Bright et al., 2009; Donahue et al., 2016; Moia et al., 2021, 2020a; Sousa et al., 2014; Stickland et al., 2021), while in a case of unilateral moyamoya, delays exceeded 10 seconds in the affected hemisphere (Stickland et al., 2021).

Our choice to use a multi-echo fMRI acquisition rather than a more commonly used single echo protocol may also limit the generalizability of our findings. In fact, the GM-BOLD signal used in our study benefits from the boost in SNR achieved from the optimal combination of 5 echoes (Cohen and Wang, 2019; Moia et al., 2021). If a multi-echo fMRI approach is not feasible, spatial smoothing or cortical parcellation could be used as alternatives to boost SNR at the cost of spatial definition. Results from CVR mapping will also be sensitive to the quality of the input fMRI data, from acquisition to the pre-processing and denoising steps applied (Caballero-Gaudes and Reynolds, 2017). As with all fMRI acquisitions, we recommend mitigating motion confounds during the scan and modelling these noise sources in the lagged-GLM (Moia et al., 2021).

CVR map quality and accuracy could be further improved by refining the response functions used to model the effects of P_ET_CO_2_ and respiration fluctuations on the BOLD signal. We assumed canonical response functions for the HRF (Friston et al., 1998) and RRF (Birn et al., 2008) used to model P_ET_CO_2_ and RVT, respectively. While we have accounted for regional variations in the timing of these responses, we have not incorporated flexible response shapes. Spatial heterogeneity in the amplitude and timing of BOLD responses to respiratory variation is apparent in resting-state data, with notable differences between primary sensory regions and frontoparietal regions (Chen et al., 2020; Pinto et al., 2017). The inclusion of temporal and derivative basis sets in the lagged-GLM as described by Chen at al., 2020 may better account for this variability. In addition, response functions have been shown to vary between subjects and even between sessions from the same subject (Kassinopoulos and Mitsis, 2019). Kassinopoulos and Mitsis (2019) proposed a framework to estimate subject-specific response functions by using a combination of optimization techniques to estimate parameters of the double gamma functions, which could also be implemented to generate more accurate reference signals. Similarly, they could be estimated from the subject-specific global or GM-BOLD signals (Falahpour et al., 2013), but importantly the estimated response functions should be tested in other datasets to avoid circularity. Regardless of the approach, differences in response functions are especially important to consider in cohorts that may have atypical hemodynamics, as in older adults and cerebrovascular pathology (D’Esposito et al., 2003). Future work should re-evaluate consistency between CVR maps with region-specific and/or subject-specific response functions.

We encourage collaboration among stakeholders in the CVR community and suggest integration among the existing approaches that aim to address the feasibility of physiological modeling and CVR mapping. For example, a refined GM-BOLD regressor could be extracted from existing algorithms such as Rapidtide (Frederick et al., 2022b) or seeCVR (Bhogal, 2022) and incorporated into the lagged-GLM. Alternatively, standard implementations of CVR modeling algorithms, including Rapidtide, seeCVR, and quantiphyse (Craig et al., 2022), could be modified to input RVT as a reference signal not already supported. Machine learning may also be a promising tool to address challenges with reference signal quality. For example, Agrawal et al., 2022 successfully used the respiratory waveform in resting-state data to predict CO_2_ and derive P_ET_CO_2_ using a fully convolutional neural network (Agrawal et al., 2022). However, their method does not maintain P_ET_CO_2_ in quantitative units of mmHg either, which would be preferred for modeling CVR amplitude. A separate study proposed two deep learning architectures (again a convolutional neural network and a fully connected single-unit network) to reconstruct respiratory variation signals from the fMRI data itself (Salas et al., 2021). Future work could adapt these models to predict a “sufficient” P_ET_CO_2_ trace from insufficient P_ET_CO_2_ data, from a respiration trace, or from the fMRI data in the context of a breath-hold task. This would be especially promising if the algorithm is able to scale the resulting P_ET_CO_2_ signal in standard units (i.e., %BOLD/mmHg).

## 5. Conclusion

End-tidal CO_2_ (P_ET_CO_2_) is commonly used as a reference signal to facilitate modeling of cerebrovascular reactivity (CVR) in BOLD fMRI data, but the P_ET_CO_2_ recordings may be unavailable or unreliable in many settings. We demonstrate that respiration volume per time (RVT) or the average gray matter BOLD response during a breath-hold task can be used in a lagged general linear model framework to obtain estimates of CVR amplitude and delay. Furthermore, CVR maps from these reference signals have good spatial agreement with those from the gold standard reference of P_ET_CO_2._ In datasets with sub-optimal or “insufficient” P_ET_CO_2_ recordings, RVT and GM-BOLD can also be used to recover reasonable CVR amplitude and delay maps, provided that the participant achieved periods of apnea during the breath-hold task. This framework offers a solution to obtain non-quantitative CVR amplitude and quantitative delay maps when reliable P_ET_CO_2_ recordings are unavailable due to limitations in resources or participant compliance.

## Acknowledgments

The authors would like to thank Andrew Vigotsky for his feedback on the visualizations in this manuscript.

## CRediT Author Contribution Statement

**Kristina Zvolanek:** Conceptualization, Methodology, Software, Formal Analysis, Data Curation, Writing – Original Draft, Writing – Revising and Editing, Visualization, Project Administration, Funding Acquisition. **Stefano Moia:** Conceptualization, Methodology, Software, Formal Analysis, Resources, Data Curation, Writing – Revising and Editing, Funding Acquisition. **Joshua Dean:** Software, Formal Analysis, Writing – Original Draft, Writing – Revising and Editing, Visualization. **Rachael Stickland:** Software, Methodology, Writing – Revising and Editing. **César Caballero-Gaudes:** Conceptualization, Methodology, Resources, Writing – Revising and Editing, Funding Acquisition. **Molly Bright:** Conceptualization, Methodology, Resources, Writing – Revising and Editing, Supervision, Project Administration, Funding Acquisition.

## References

Agrawal, V., Zhong, X.Z., Chen, J.J., Chen, J., 2022. Generating dynamic carbon-dioxide from the respiratory-volume time series: A feasibility study using neural networks. bioRxiv 2022.07.11.499585. https://doi.org/10.1101/2022.07.11.499585

Andersson, J.L.R., Skare, S., Ashburner, J., 2003. How to correct susceptibility distortions in spin-echo echo-planar images: application to diffusion tensor imaging. Neuroimage 20, 870–888. https://doi.org/10.1016/S1053-8119(03)00336-7

Atwi, S., Shao, H., Crane, D.E., da Costa, L., Aviv, R.I., Mikulis, D.J., Black, S.E., MacIntosh, B.J., 2019. BOLD-based cerebrovascular reactivity vascular transfer function isolates amplitude and timing responses to better characterize cerebral small vessel disease. NMR Biomed. 32, e4064. https://doi.org/10.1002/NBM.4064

Bhogal, A.A., 2022. abhogal-lab/seeVR: 1.5. https://doi.org/10.5281/ZENODO.6532362

Birn, R.M., Diamond, J.B., Smith, M.A., Bandettini, P.A., 2006. Separating respiratory-variation-related fluctuations from neuronal-activity-related fluctuations in fMRI. Neuroimage 31, 1536–1548. https://doi.org/10.1016/j.neuroimage.2006.02.048

Birn, R.M., Smith, M.A., Jones, T.B., Bandettini, P.A., 2008. The respiration response function: The temporal dynamics of fMRI signal fluctuations related to changes in respiration. Neuroimage 40, 644–654. https://doi.org/10.1016/j.neuroimage.2007.11.059

Blair, G.W., Doubal, F.N., Thrippleton, M.J., Marshall, I., Wardlaw, J.M., 2015. Magnetic resonance imaging for assessment of cerebrovascular reactivity in cerebral small vessel disease: A systematic review. J. Cereb. Blood Flow Metab. 36, 833–841. https://doi.org/10.1177/0271678X16631756/ASSET/IMAGES/LARGE/10.1177_0271678X16631756-FIG1.JPEG

Botvinik-Nezer, R., Holzmeister, F., Camerer, C.F., Dreber, A., Huber, J., Johannesson, M., Kirchler, M., Iwanir, R., Mumford, J.A., Adcock, R.A., Avesani, P., Baczkowski, B.M., Bajracharya, A., Bakst, L., Ball, S., Barilari, M., Bault, N., Beaton, D., Beitner, J., Benoit, R.G., Berkers, R.M.W.J., Bhanji, J.P., Biswal, B.B., Bobadilla-Suarez, S., Bortolini, T., Bottenhorn, K.L., Bowring, A., Braem, S., Brooks, H.R., Brudner, E.G., Calderon, C.B., Camilleri, J.A., Castrellon, J.J., Cecchetti, L., Cieslik, E.C., Cole, Z.J., Collignon, O., Cox, R.W., Cunningham, W.A., Czoschke, S., Dadi, K., Davis, C.P., Luca, A. De, Delgado, M.R., Demetriou, L., Dennison, J.B., Di, X., Dickie, E.W., Dobryakova, E., Donnat, C.L., Dukart, J., Duncan, N.W., Durnez, J., Eed, A., Eickhoff, S.B., Erhart, A., Fontanesi, L., Fricke, G.M., Fu, S., Galván, A., Gau, R., Genon, S., Glatard, T., Glerean, E., Goeman, J.J., Golowin, S.A.E., González-García, C., Gorgolewski, K.J., Grady, C.L., Green, M.A., Guassi Moreira, J.F., Guest, O., Hakimi, S., Hamilton, J.P., Hancock, R., Handjaras, G., Harry, B.B., Hawco, C., Herholz, P., Herman, G., Heunis, S., Hoffstaedter, F., Hogeveen, J., Holmes, S., Hu, C.P., Huettel, S.A., Hughes, M.E., Iacovella, V., Iordan, A.D., Isager, P.M., Isik, A.I., Jahn, A., Johnson, M.R., Johnstone, T., Joseph, M.J.E., Juliano, A.C., Kable, J.W., Kassinopoulos, M., Koba, C., Kong, X.Z., Koscik, T.R., Kucukboyaci, N.E., Kuhl, B.A., Kupek, S., Laird, A.R., Lamm, C., Langner, R., Lauharatanahirun, N., Lee, H., Lee, S., Leemans, A., Leo, A., Lesage, E., Li, F., Li, M.Y.C., Lim, P.C., Lintz, E.N., Liphardt, S.W., Losecaat Vermeer, A.B., Love, B.C., Mack, M.L., Malpica, N., Marins, T., Maumet, C., McDonald, K., McGuire, J.T., Melero, H., Méndez Leal, A.S., Meyer, B., Meyer, K.N., Mihai, G., Mitsis, G.D., Moll, J., Nielson, D.M., Nilsonne, G., Notter, M.P., Olivetti, E., Onicas, A.I., Papale, P., Patil, K.R., Peelle, J.E., Pérez, A., Pischedda, D., Poline, J.B., Prystauka, Y., Ray, S., Reuter-Lorenz, P.A., Reynolds, R.C., Ricciardi, E., Rieck, J.R., Rodriguez-Thompson, A.M., Romyn, A., Salo, T., Samanez-Larkin, G.R., Sanz-Morales, E., Schlichting, M.L., Schultz, D.H., Shen, Q., Sheridan, M.A., Silvers, J.A., Skagerlund, K., Smith, A., Smith, D. V., Sokol-Hessner, P., Steinkamp, S.R., Tashjian, S.M., Thirion, B., Thorp, J.N., Tinghög, G., Tisdall, L., Tompson, S.H., Toro-Serey, C., Torre Tresols, J.J., Tozzi, L., Truong, V., Turella, L., van ‘t Veer, A.E., Verguts, T., Vettel, J.M., Vijayarajah, S., Vo, K., Wall, M.B., Weeda, W.D., Weis, S., White, D.J., Wisniewski, D., Xifra-Porxas, A., Yearling, E.A., Yoon, S., Yuan, R., Yuen, K.S.L., Zhang, L., Zhang, X., Zosky, J.E., Nichols, T.E., Poldrack, R.A., Schonberg, T., 2020. Variability in the analysis of a single neuroimaging dataset by many teams. Nat. 2020 5827810 582, 84–88. https://doi.org/10.1038/s41586-020-2314-9

Bright, M.G., Bulte, D.P., Jezzard, P., Duyn, J.H., 2009. Characterization of regional heterogeneity in cerebrovascular reactivity dynamics using novel hypocapnia task and BOLD fMRI. Neuroimage 48, 166–175. https://doi.org/10.1016/j.neuroimage.2009.05.026

Bright, M.G., Murphy, K., 2013. Reliable quantification of BOLD fMRI cerebrovascular reactivity despite poor breath-hold performance. Neuroimage. https://doi.org/10.1016/j.neuroimage.2013.07.007

Bulte, D., Wartolowska, K., 2017. Monitoring cardiac and respiratory physiology during FMRI. Neuroimage 154, 81–91. https://doi.org/10.1016/j.neuroimage.2016.12.001

Caballero-Gaudes, C., Reynolds, R.C., 2017. Methods for cleaning the BOLD fMRI signal. Neuroimage 154, 128–149. https://doi.org/10.1016/j.neuroimage.2016.12.018

Cauley, S.F., Polimeni, J.R., Bhat, H., Wald, L.L., Setsompop, K., 2014. Interslice leakage artifact reduction technique for simultaneous multislice acquisitions. Magn. Reson. Med. 72, 93–102. https://doi.org/10.1002/MRM.24898

Chang, C., Glover, G.H., 2009. Relationship between respiration, end-tidal CO2, and BOLD signals in resting-state fMRI. Neuroimage 47, 1381–1393. https://doi.org/10.1016/j.neuroimage.2009.04.048

Chang, C., Thomason, M.E., Glover, G.H., 2008. Mapping and correction of vascular hemodynamic latency in the BOLD signal. Neuroimage 43, 90–102. https://doi.org/10.1016/J.NEUROIMAGE.2008.06.030

Chen, J.E., Lewis, L.D., Chang, C., Tian, Q., Fultz, N.E., Ohringer, N.A., Rosen, B.R., Polimeni, J.R., 2020. Resting-state “physiological networks.” Neuroimage 213, 116707. https://doi.org/10.1016/j.neuroimage.2020.116707

Cohen, A.D., Wang, Y., 2019. Improving the Assessment of Breath-Holding Induced Cerebral Vascular Reactivity Using a Multiband Multi-echo ASL/BOLD Sequence. Sci. Rep. 9, 1–12. https://doi.org/10.1038/s41598-019-41199-w

Conklin, J., Fierstra, J., Crawley, A.P., Han, J.S., Poublanc, J., Mandell, D.M., Silver, F.L., Tymianski, M., Fisher, J.A., Mikulis, D.J., 2010. Impaired cerebrovascular reactivity with steal phenomenon is associated with increased diffusion in white matter of patients with moyamoya disease. Stroke 41, 1610–1616. https://doi.org/10.1161/STROKEAHA.110.579540/FORMAT/EPUB

Cox, R.W., 1996. AFNI: Software for Analysis and Visualization of Functional Magnetic Resonance Neuroimages. Comput. Biomed. Res. 29, 162–173. https://doi.org/10.1006/CBMR.1996.0014

Craig, M., Irving, B., Chappell, M., Croal, P., Zhao, M., 2022. physimals/quantiphysev0.9.9 [WWW Document]. URL https://quantiphyse.readthedocs.io/en/latest/index.html (accessed 10.10.22).

D’Esposito, M., Deouell, L.Y., Gazzaley, A., 2003. Alterations in the BOLD fMRI signal with ageing and disease: a challenge for neuroimaging. Nat. Rev. Neurosci. 2003 411 4, 863–872. https://doi.org/10.1038/nrn1246

Dafflon, J., F. Da Costa, P., Váša, F., Monti, R.P., Bzdok, D., Hellyer, P.J., Turkheimer, F., Smallwood, J., Jones, E., Leech, R., 2022. A guided multiverse study of neuroimaging analyses. Nat. Commun. 2022 131 13, 1–13. https://doi.org/10.1038/s41467-022-31347-8

Di, X., Kim, E.H., Huang, C.C., Tsai, S.J., Lin, C.P., Biswal, B.B., 2013. The influence of the amplitude of low-frequency fluctuations on resting-state functional connectivity. Front. Hum. Neurosci. 0, 118. https://doi.org/10.3389/FNHUM.2013.00118/BIBTEX

Diedrichsen, J., Verstynen, T., Schlerf, J., Wiestler, T., 2010. Advances in functional imaging of the human cerebellum. Curr. Opin. Neurol. 23, 382–387. https://doi.org/10.1097/WCO.0B013E32833BE837

Dlamini, N., Shah-Basak, P., Leung, J., Kirkham, F., Shroff, M., Kassner, A., Robertson, A., Dirks, P., Westmacott, R., De Veber, G., Logan, W., 2018. Breath-hold blood oxygen level-dependent MRI: A tool for the assessment of cerebrovascular reserve in children with moyamoya disease. Am. J. Neuroradiol. 39, 1717–1723. https://doi.org/10.3174/ajnr.A5739

Donahue, M.J., Dethrage, L.M., Faraco, C.C., Jordan, L.C., Clemmons, P., Singer, R., Mocco, J., Shyr, Y., Desai, A., O’Duffy, A., Riebau, D., Hermann, L., Connors, J., Kirshner, H., Strother, M.K., 2014. Routine clinical evaluation of cerebrovascular reserve capacity using carbogen in patients with intracranial stenosis. Stroke 45, 2335–41. https://doi.org/10.1161/STROKEAHA.114.005975

Donahue, M.J., Strother, M.K., Lindsey, K.P., Hocke, L.M., Tong, Y., Frederick, B.D.B., 2016. Time delay processing of hypercapnic fMRI allows quantitative parameterization of cerebrovascular reactivity and blood flow delays. J. Cereb. Blood Flow Metab. 36, 1767– 1779. https://doi.org/10.1177/0271678X15608643

Duffin, J., Sobczyk, O., Crawley, A.P., Poublanc, J., Mikulis, D.J., Fisher, J.A., 2015. The dynamics of cerebrovascular reactivity shown with transfer function analysis. Neuroimage 114, 207–216. https://doi.org/10.1016/j.neuroimage.2015.04.029

DuPre, E., Salo, T., Ahmed, Z., Bandettini, P.A., Bottenhorn, K.L., Caballero-Gaudes, C., Dowdle, L.T., Gonzalez-Castillo, J., Heunis, S., Kundu, P., Laird, A.R., Markello, R., Markiewicz, C.J., Moia, S., Staden, I., Teves, J.B., Uruñuela, E., Vaziri-Pashkam, M., Whitaker, K., Handwerker, D.A., 2021. TE-dependent analysis of multi-echo fMRI with *tedana*. J. Open Source Softw. 6, 3669. https://doi.org/10.21105/JOSS.03669

DuPre, E., Salo, T., Markello, R., Kundu, P., Whitaker, K., Handwerker, D., 2019. ME-ICA/tedana: 0.0.6. https://doi.org/10.5281/ZENODO.2558498

Erdoğan, S.B., Tong, Y., Hocke, L.M., Lindsey, K.P., deB Frederick, B., 2016. Correcting for blood arrival time in global mean regression enhances functional connectivity analysis of resting state fMRI-BOLD signals. Front. Hum. Neurosci. 10, 311. https://doi.org/10.3389/FNHUM.2016.00311/BIBTEX

Falahpour, M., Refai, H., Bodurka, J., 2013. Subject specific BOLD fMRI respiratory and cardiac response functions obtained from global signal. Neuroimage 72, 252–264. https://doi.org/10.1016/j.neuroimage.2013.01.050

Fierstra, J., Sobczyk, O., Battisti-Charbonney, A., Mandell, D.M., Poublanc, J., Crawley, A.P., Mikulis, D.J., Duffin, J., Fisher, J.A., 2013. Measuring cerebrovascular reactivity: What stimulus to use? J. Physiol. 591, 5809–5821. https://doi.org/10.1113/jphysiol.2013.259150

Fierstra, J., van Niftrik, C., Piccirelli, M., Bozinov, O., Pangalu, A., Krayenbühl, N., Valavanis, A., Weller, M., Regli, L., 2018. Diffuse gliomas exhibit whole brain impaired cerebrovascular reactivity. Magn. Reson. Imaging 45, 78–83. https://doi.org/10.1016/J.MRI.2017.09.017

Frederick, B. de B., Nickerson, L.D., Tong, Y., 2012. Physiological denoising of BOLD fMRI data using Regressor Interpolation at Progressive Time Delays (RIPTiDe) processing of concurrent fMRI and near-infrared spectroscopy (NIRS). Neuroimage 60, 1913–1923. https://doi.org/10.1016/J.NEUROIMAGE.2012.01.140

Frederick, B. deB., Salo, T., Drucker, D.M., 2016. Rapidtide. https://doi.org/10.5281/zenodo.814990

Frederick, B. deB, 2017. bbfrederick/rapidtide: December 2017 checkpoint release. https://doi.org/10.5281/ZENODO.1119128

Frederick, B. deB, Salo, T., Drucker, D.M., 2022a. bbfrederick/rapidtide: Version 2.2.7 - 6/29/22 checkpoint. https://doi.org/10.5281/ZENODO.6780450

Frederick, B. deB, Salo, T., Drucker, D.M., 2022b. bbfrederick/rapidtide: Version 2.2.8.1 - 8/29/22 deployment bug fix. https://doi.org/10.5281/ZENODO.7032879

Friston, K.J., Fletcher, P., Josephs, O., Holmes, A., Rugg, M.D., Turner, R., 1998. Event-related fMRI: Characterizing differential responses. Neuroimage 7, 30–40. https://doi.org/10.1006/nimg.1997.0306

Geranmayeh, F., Wise, R.J.S., Leech, R., Murphy, K., 2015. Measuring Vascular Reactivity With Breath-Holds After Stroke: A Method to Aid Interpretation of Group-Level BOLD Signal Changes in Longitudinal fMRI Studies. Hum Brain Mapp 36, 1755–1771. https://doi.org/10.1002/hbm.22735

Golestani, A.M., Wei, L.L., Chen, J.J., 2016. Quantitative mapping of cerebrovascular reactivity using resting-state BOLD fMRI: Validation in healthy adults. Neuroimage 138, 147–163. https://doi.org/10.1016/j.neuroimage.2016.05.025

Gorgolewski, K.J., Auer, T., Calhoun, V.D., Craddock, R.C., Das, S., Duff, E.P., Flandin, G., Ghosh, S.S., Glatard, T., Halchenko, Y.O., Handwerker, D.A., Hanke, M., Keator, D., Li, X., Michael, Z., Maumet, C., Nichols, B.N., Nichols, T.E., Pellman, J., Poline, J.B., Rokem, A., Schaefer, G., Sochat, V., Triplett, W., Turner, J.A., Varoquaux, G., Poldrack, R.A., 2016. The brain imaging data structure, a format for organizing and describing outputs of neuroimaging experiments. Sci. Data 2016 31 3, 1–9. https://doi.org/10.1038/sdata.2016.44

Grabner, G., Janke, A.L., Budge, M.M., Smith, D., Pruessner, J., Collins, D.L., 2006. Symmetric atlasing and model based segmentation: An application to the hippocampus in older adults. Lect. Notes Comput. Sci. (including Subser. Lect. Notes Artif. Intell. Lect. Notes Bioinformatics) 4191 LNCS-II, 58–66. https://doi.org/10.1007/11866763_8/COVER

Gupta, A., Chazen, J.L., Hartman, M., Delgado, D., Anumula, N., Shao, H., Mazumdar, M., Segal, A.Z., Kamel, H., Leifer, D., Sanelli, P.C., 2012. Cerebrovascular reserve and stroke risk in patients with carotid stenosis or occlusion: A systematic review and meta-analysis. Stroke 43, 2884–2891. https://doi.org/10.1161/STROKEAHA.112.663716/-/DC1

Halchenko, Y., Goncalves, M., Castello, M.V. di O., Ghosh, S., Hanke, M., Dae, Salo, T., Kent, J., Amlien, I., Brett, M., Tilley, S., Markiewicz, C., Gorgolewski, C., pvelasco, Kim, S., Stadler, J., Kaczmarzyk, J., Lukas, D.C., lee, john, Lurie, D., Pellman, J., Braun, H., Melo, B., Poldrack, B., Nichols, T., Schiffler, B., Szczepanik, M., Carlin, J., Feingold, F., Kahn, A., 2019. nipy/heudiconv v0.6.0. https://doi.org/10.5281/ZENODO.3579455

Handwerker, D.A., Gazzaley, A., Inglis, B.A., D’Esposito, M., 2007. Reducing vascular variability of fMRI data across aging populations using a breathholding task. Hum. Brain Mapp. 28, 846–859. https://doi.org/10.1002/hbm.20307

Hartkamp, N.S., Hendrikse, J., Van Der Worp, H.B., De Borst, G.J., Bokkers, R.P.H., 2012. Time course of vascular reactivity using repeated phase-contrast MR angiography in patients with carotid artery stenosis. Stroke 43, 553–556. https://doi.org/10.1161/STROKEAHA.111.637314

Holmes, K.R., Tang-Wai, D., Sam, K., Mcketton, L., Poublanc, J., Crawley, A.P., Sobczyk, O., Cohn, M., Duffin, J., Tartaglia, M.C., Black, S.E., Fisher, J.A., Wasserman, B., Mikulis, D.J., 2020. Slowed Temporal and Parietal Cerebrovascular Response in Patients with Alzheimer’s Disease. Can. J. Neurol. Sci. 47, 366–373. https://doi.org/10.1017/CJN.2020.30

Howell, D.C., 2010. Testing the Difference Between Two Independent rs, in: Statistical Methods for Psychology. pp. 275–276.

Jenkinson, M., Bannister, P., Brady, M., Smith, S., 2002. Improved Optimization for the Robust and Accurate Linear Registration and Motion Correction of Brain Images. Neuroimage 17, 825–841. https://doi.org/10.1006/NIMG.2002.1132

Jenkinson, M., Beckmann, C.F., Behrens, T.E.J., Woolrich, M.W., Smith, S.M., 2012. FSL. Neuroimage 62, 782–790. https://doi.org/10.1016/J.NEUROIMAGE.2011.09.015

Jenkinson, M., Smith, S., 2001. A global optimisation method for robust affine registration of brain images. Med. Image Anal. 5, 143–156. https://doi.org/10.1016/S1361-8415(01)00036-6

Juttukonda, M.R., Donahue, M.J., 2019. Neuroimaging of vascular reserve in patients with cerebrovascular diseases. Neuroimage. https://doi.org/10.1016/j.neuroimage.2017.10.015

Kannurpatti, S.S., Motes, M.A., Rypma, B., Biswal, B.B., 2010. Neural and vascular variability and the fMRI-BOLD response in normal aging. Magn. Reson. Imaging 28, 466–476. https://doi.org/10.1016/j.mri.2009.12.007

Kassinopoulos, M., Mitsis, G.D., 2019. Identification of physiological response functions to correct for fluctuations in resting-state fMRI related to heart rate and respiration. Neuroimage 202, 116150. https://doi.org/10.1016/j.neuroimage.2019.116150

Kastrup, A., Krüger, G., Neumann-Haefelin, T., Moseley, M.E., 2001. Assessment of cerebrovascular reactivity with functional magnetic resonance imaging: Comparison of CO2 and breath holding. Magn. Reson. Imaging 19, 13–20. https://doi.org/10.1016/S0730-725X(01)00227-2

Kastrup, A., Li, T.-Q., Glover, G.H., Moseley, M.E., 1999. Cerebral Blood Flow-Related Signal Changes during Breath-Holding. AJNR Am J Neuroradiol 20, 1233–1238.

Kazan, S.M., Mohammadi, S., Callaghan, M.F., Flandin, G., Huber, L., Leech, R., Kennerley, A., Windischberger, C., Weiskopf, N., 2016. Vascular autorescaling of fMRI (VasA fMRI) improves sensitivity of population studies: A pilot study. Neuroimage 124, 794–805. https://doi.org/10.1016/J.NEUROIMAGE.2015.09.033

Krainik, A., Hund-Georgiadis, M., Zysset, S., Yves Von Cramon, ; D, 2005. Regional Impairment of Cerebrovascular Reactivity and BOLD Signal in Adults After Stroke. https://doi.org/10.1161/01.STR.0000166178.40973.a7

Leung, J., Duffin, J., Fisher, J.A., Kassner, A., 2016a. MRI-based cerebrovascular reactivity using transfer function analysis reveals temporal group differences between patients with sickle cell disease and healthy controls. NeuroImage Clin. 12, 624–630. https://doi.org/10.1016/j.nicl.2016.09.009

Leung, J., Kosinski, P.D., Croal, P.L., Kassner, A., 2016b. Developmental trajectories of cerebrovascular reactivity in healthy children and young adults assessed with magnetic resonance imaging. J. Physiol. 594, 2681–2689. https://doi.org/10.1113/JP271056

Lipp, I., Murphy, K., Caseras, X., Wise, R.G., 2015. Agreement and repeatability of vascular reactivity estimates based on a breath-hold task and a resting state scan. Neuroimage 113, 387–396. https://doi.org/10.1016/J.NEUROIMAGE.2015.03.004

Liu, P., De Vis, J.B., 2019. Cerebrovascular reactivity (CVR) MRI with CO2 challenge: A technical review. Neuroimage 187, 104–115. https://doi.org/10.1016/J.NEUROIMAGE.2018.03.047

Liu, P., De Vis, J.B., Lu, H., 2019. Cerebrovascular reactivity (CVR) MRI with CO2 challenge: A technical review. Neuroimage 187, 104–115. https://doi.org/10.1016/j.neuroimage.2018.03.047

Liu, P., Li, Y., Pinho, M., Park, D.C., Welch, B.G., Lu, H., 2017. Cerebrovascular reactivity mapping without gas challenges. Neuroimage 146, 320–326. https://doi.org/10.1016/j.neuroimage.2016.11.054

Liu, P., Liu, G., Pinho, M.C., Lin, Z., Thomas, B.P., Rundle, M., Park, D.C., Huang, J., Welch, B.G., Lu, H., 2021. Cerebrovascular reactivity Mapping using resting-state bold functiona MRI in healthy adults and patients with moyamoya disease. Radiology 299, 419–425. https://doi.org/10.1148/RADIOL.2021203568/ASSET/IMAGES/LARGE/RADIOL.2021203568.FIG6.JPEG

Liu, P., Xu, C., Lin, Z., Sur, S., Li, Y., Yasar, S., Rosenberg, P., Albert, M., Lu, H., 2020. Cerebrovascular reactivity mapping using intermittent breath modulation. Neuroimage 215, 116787. https://doi.org/10.1016/j.neuroimage.2020.116787

Liu, T.T., 2016. Noise contributions to the fMRI signal: An overview. Neuroimage 143, 141–151. https://doi.org/10.1016/J.NEUROIMAGE.2016.09.008

Magon, S., Basso, G., Farace, P., Ricciardi, G.K., Beltramello, A., Sbarbati, A., 2009. Reproducibility of BOLD signal change induced by breath holding. Neuroimage 45, 702– 712. https://doi.org/10.1016/j.neuroimage.2008.12.059

Marshall, O., Lu, H., Brisset, J.C., Xu, F., Liu, P., Herbert, J., Grossman, R.I., Ge, Y., 2014. Impaired cerebrovascular reactivity in multiple sclerosis. JAMA Neurol. 71, 1275–1281. https://doi.org/10.1001/jamaneurol.2014.1668

McKetton, L., Sobczyk, O., Duffin, J., Poublanc, J., Sam, K., Crawley, A.P., Venkatraghavan, L., Fisher, J.A., Mikulis, D.J., 2018. The aging brain and cerebrovascular reactivity. Neuroimage 181, 132–141. https://doi.org/10.1016/j.neuroimage.2018.07.007

McKetton, L., Venkatraghavan, L., Rosen, C., Mandell, D.M., Sam, K., Sobczyk, O., Poublanc, J., Gray, E., Crawley, A., Duffin, J., Fisher, J.A., Mikulis, D., 2019. Improved white matter cerebrovascular reactivity after revascularization in patients with steno-occlusive disease. Am. J. Neuroradiol. 40, 45–50. https://doi.org/10.3174/ajnr.A5912

McSwain, S.D., Hamel, D.S., Smith, P.B., Gentile, M.A., Srinivasan, S., Meliones, J.N., Cheifetz, I.M., 2010. End-tidal and arterial carbon dioxide measurements correlate across all levels of physiologic dead space. Respir. Care 55, 288–293.

Mikulis, D.J., Krolczyk, G., Desal, H., Logan, W., DeVeber, G., Dirks, P., Tymianski, M., Crawley, A., Vesely, A., Kassner, A., Preiss, D., Somogyi, R., Fisher, J.A., 2005. Preoperative and postoperative mapping of cerebrovascular reactivity in moyamoya disease by using blood oxygen level-dependent magnetic resonance imaging. J. Neurosurg. 103, 347–355. https://doi.org/10.3171/jns.2005.103.2.0347

Moia, S., Chen, G., Urunuela, E., Stickland, R., Termenon, M., Caballero-Gaudes, C., Bright, M., 2022a. Resting state fluctuations in BOLD fMRI might not systematically reflect measures of cerebrovascular physiology between or within subjects, in: International Society of Magnetic Resonance in Medicine (ISMRM) 31st Annual Meeting & Exhibition. London, England, UK.

Moia, S., Stickland, R.C., Ayyagari, A., Termenon, M., Caballero-Gaudes, C., Bright, M.G., 2020a. Voxelwise optimization of hemodynamic lags to improve regional CVR estimates in breath-hold fMRI. Proc. Annu. Int. Conf. IEEE Eng. Med. Biol. Soc. EMBS 2020-July, 1489–1492. https://doi.org/10.1109/EMBC44109.2020.9176225

Moia, S., Termenon, M., Uruñuela, E., Chen, G., Stickland, R.C., Bright, M.G., Caballero-Gaudes, C., 2021. ICA-based denoising strategies in breath-hold induced cerebrovascular reactivity mapping with multi echo BOLD fMRI. Neuroimage 233, 117914. https://doi.org/10.1016/J.NEUROIMAGE.2021.117914

Moia, S., Uruñuela, E., Ferrer, V., Caballero-Gaudes, C., 2020b. EuskalIBUR. OpenNeuro. https://doi.org/10.18112/OPENNEURO.DS003192.V1.0.1

Moia, S., Vigotsky, A.D., Zvolanek, K.M., 2022b. physiopy/phys2cvr: A tool to compute Cerebrovascular Reactivity maps and associated lag maps. https://doi.org/10.5281/ZENODO.7336002

Murphy, K., Birn, R.M., Bandettini, P.A., 2013. Resting-state fMRI confounds and cleanup. Neuroimage 80, 349–359. https://doi.org/10.1016/j.neuroimage.2013.04.001

Murphy, K., Harris, A.D., Wise, R.G., 2011. Robustly measuring vascular reactivity differences with breath-hold: Normalising stimulus-evoked and resting state BOLD fMRI data. Neuroimage 54, 369–379. https://doi.org/10.1016/j.neuroimage.2010.07.059

Peebles, K., Celi, L., McGrattan, K., Murrell, C., Thomas, K., Ainslie, P.N., 2007. Human cerebrovascular and ventilatory CO2 reactivity to end-tidal, arterial and internal jugular vein PCO2. J. Physiol. 584, 347–357. https://doi.org/10.1113/JPHYSIOL.2007.137075

Pillai, J.J., Mikulis, D.J., 2015. Cerebrovascular reactivity mapping: An evolving standard for clinical functional imaging. Am. J. Neuroradiol. https://doi.org/10.3174/ajnr.A3941

Pinto, J., Bright, M.G., Bulte, D.P., Figueiredo, P., 2021. Cerebrovascular Reactivity Mapping Without Gas Challenges: A Methodological Guide. Front. Physiol. 11, 1711. https://doi.org/10.3389/fphys.2020.608475

Pinto, J., Jorge, J., Sousa, I., Vilela, P., Figueiredo, P., 2016. Fourier modeling of the BOLD response to a breath-hold task: Optimization and reproducibility. Neuroimage 135, 223– 231. https://doi.org/10.1016/J.NEUROIMAGE.2016.02.037

Pinto, J., Nunes, S., Bianciardi, M., Dias, A., Silveira, L.M., Wald, L.L., Figueiredo, P., 2017. Improved 7 Tesla resting-state fMRI connectivity measurements by cluster-based modeling of respiratory volume and heart rate effects. Neuroimage 153, 262. https://doi.org/10.1016/J.NEUROIMAGE.2017.04.009

Posse, S., Wiese, S., Gembris, D., Mathiak, K., Kessler, C., Grosse-Ruyken, M.-L., Elghahwagi, B., Richards, T., Dager, S.R., Kiselev, V.G., 1999. Enhancement of BOLD-Contrast Sensitivity by Single-Shot Multi-Echo Functional MR Imaging. Magn Reson Med 42, 87–97. https://doi.org/10.1002/(SICI)1522-2594(199907)42:1

Poublanc, J., Show, J., Daniel, H., Mandell, M., Conklin, J., Stainsby, J.A., Arnold, J., David, F., Mikulis, J., Crawley, A.P., 2013. Vascular Steal Explains Early Paradoxical Blood Oxygen Level-Dependent Cerebrovascular Response in Brain Regions with Delayed Arterial Transit Times. Cerebrovasc. Dis. Extra 3, 55–64. https://doi.org/10.1159/000348841

Pujol, J., Conesa, G., Deus, J., López-Obarrio, L., Isamat, F., Capdevila, A., 1998. Clinical application of functional magnetic resonance imaging in presurgical identification of the central sulcus. J. Neurosurg. 88, 863–869. https://doi.org/10.3171/JNS.1998.88.5.0863

Ratnatunga, C., Adiseshiah, M., 1990. Increase in middle cerebral artery velocity on breath holding: A simplified test of cerebral perfusion reserve. Eur. J. Vasc. Surg. 4, 519–523. https://doi.org/10.1016/S0950-821X(05)80795-9

Salas, J.A., Bayrak, R.G., Huo, Y., Chang, C., 2021. Reconstruction of respiratory variation signals from fMRI data. Neuroimage 225, 117459. https://doi.org/10.1016/j.neuroimage.2020.117459

Sam, K., Conklin, J., Holmes, K.R., Sobczyk, O., Poublanc, J., Crawley, A.P., Mandell, D.M., Venkatraghavan, L., Duffin, J., Fisher, J.A., Black, S.E., Mikulis, D.J., 2016. Impaired dynamic cerebrovascular response to hypercapnia predicts development of white matter hyperintensities. NeuroImage Clin. 11, 796–801. https://doi.org/10.1016/J.NICL.2016.05.008

Schouwenaars, I.T., De Dreu, M.J., Rutten, G.J.M., Ramsey, N.F., Jansma, J.M., 2021. A functional MRI study of presurgical cognitive deficits in glioma patients. Neuro-Oncology Pract. 8, 81–90. https://doi.org/10.1093/NOP/NPAA059

Scouten, A., Schwarzbauer, C., 2008. Paced respiration with end-expiration technique offers superior BOLD signal repeatability for breath-hold studies. Neuroimage 43, 250–257. https://doi.org/10.1016/j.neuroimage.2008.03.052

Sleight, E., Stringer, M.S., Marshall, I., Wardlaw, J.M., Thrippleton, M.J., 2021. Cerebrovascular Reactivity Measurement Using Magnetic Resonance Imaging: A Systematic Review. Front. Physiol. https://doi.org/10.3389/fphys.2021.643468

Smeeing, D.P.J., Hendrikse, J., Petersen, E.T., Donahue, M.J., de Vis, J.B., 2016. Arterial Spin Labeling and Blood Oxygen Level-Dependent MRI Cerebrovascular Reactivity in Cerebrovascular Disease: A Systematic Review and Meta-Analysis. Cerebrovasc. Dis. 42, 288–307. https://doi.org/10.1159/000446081

Sotiropoulos, S.N., Moeller, S., Jbabdi, S., Xu, J., Andersson, J.L., Auerbach, E.J., Yacoub, E., Feinberg, D., Setsompop, K., Wald, L.L., Behrens, T.E.J., Ugurbil, K., Lenglet, C., 2013. Effects of image reconstruction on fiber orientation mapping from multichannel diffusion MRI: Reducing the noise floor using SENSE. Magn. Reson. Med. 70, 1682–1689. https://doi.org/10.1002/MRM.24623

Sousa, I., Vilela, P., Figueiredo, P., 2014. Reproducibility of hypocapnic cerebrovascular reactivity measurements using BOLD fMRI in combination with a paced deep breathing task. Neuroimage 98, 31–41. https://doi.org/10.1016/j.neuroimage.2014.04.049

Steegen, S., Tuerlinckx, F., Gelman, A., Vanpaemel, W., 2016. Increasing Transparency Through a Multiverse Analysis. Perspect. Psychol. Sci. 11, 702–712. https://doi.org/10.1177/1745691616658637/ASSET/IMAGES/LARGE/10.1177_1745691616658637-FIG2.JPEG

Stickland, R.C., Zvolanek, K.M., Moia, S., Ayyagari, A., Caballero-Gaudes, C., Bright, M.G., 2021. A practical modification to a resting state fMRI protocol for improved characterization of cerebrovascular function. Neuroimage 239, 118306. https://doi.org/10.1016/j.neuroimage.2021.118306

Tancredi, F.B., Hoge, R.D., 2013. Comparison of cerebral vascular reactivity measures obtained using breath-holding and CO2 inhalation. J. Cereb. Blood Flow Metab. 33, 1066–74. https://doi.org/10.1038/jcbfm.2013.48

Taylor, P.A., Reynolds, R.C., Calhoun, V., Gonzalez-Castillo, J., Handwerker, D.A., Bandettini, P.A., Mejia, A.F., Chen, G., 2022. Highlight Results, Don’t Hide Them: Enhance interpretation, reduce biases and improve reproducibility. bioRxiv 2022.10.26.513929. https://doi.org/10.1101/2022.10.26.513929

The phys2bids developers, Alcalá, D., Ayyagari, A., Bright, M., Ferrer, V., Gaudes, C.C., Hayashi, S., Markello, R., Moia, S., Stickland, R., Uruñuela, E., Zvolanek, K., 2019. physiopy/phys2bids: BIDS formatting of physiological recordings. https://doi.org/10.5281/ZENODO.3586045

Thomas, B.P., Liu, P., Park, D.C., van Osch, M.J.P., Lu, H., 2014. Cerebrovascular reactivity in the brain white matter: magnitude, temporal characteristics, and age effects. J. Cereb. Blood Flow Metab. 34, 242–7. https://doi.org/10.1038/jcbfm.2013.194

Thomason, M.E., Burrows, B.E., Gabrieli, J.D.E., Glover, G.H., 2005. Breath holding reveals differences in fMRI BOLD signal in children and adults. Neuroimage 25, 824–837. https://doi.org/10.1016/j.neuroimage.2004.12.026

Thomason, M.E., Foland, L.C., Glover, G.H., 2007. Calibration of BOLD fMRI using breath holding reduces group variance during a cognitive task. Hum. Brain Mapp. 28, 59–68. https://doi.org/10.1002/HBM.20241

Thrippleton, M.J., Shi, Y., Blair, G., Hamilton, I., Waiter, G., Schwarzbauer, C., Pernet, C., Andrews, P.J.D., Marshall, I., Doubal, F., Wardlaw, J.M., 2018. Cerebrovascular reactivity measurement in cerebral small vessel disease: Rationale and reproducibility of a protocol for MRI acquisition and image processing. Int. J. Stroke 13, 195–206. https://doi.org/10.1177/1747493017730740

Tong, Y., Bergethon, P.R., Frederick, B. de B., 2011. An improved method for mapping cerebrovascular reserve using concurrent fMRI and near-infrared spectroscopy with Regressor Interpolation at Progressive Time Delays (RIPTiDe). Neuroimage 56, 2047– 2057. https://doi.org/10.1016/J.NEUROIMAGE.2011.03.071

Tong, Y., Frederick, B. deB., 2014. Tracking cerebral blood flow in BOLD fMRI using recursively generated regressors. Hum. Brain Mapp. 35, 5471. https://doi.org/10.1002/HBM.22564

Tong, Y., Hocke, L.M., Frederick, B.B., 2019. Low Frequency Systemic Hemodynamic “Noise” in Resting State BOLD fMRI: Characteristics, Causes, Implications, Mitigation Strategies, and Applications. Front. Neurosci. 0, 787. https://doi.org/10.3389/FNINS.2019.00787

Tsvetanov, K.A., Henson, R.N.A., Tyler, L.K., Davis, S.W., Shafto, M.A., Taylor, J.R., Williams, N., Cam-CAN, Rowe, J.B., 2015. The effect of ageing on fMRI: Correction for the confounding effects of vascular reactivity evaluated by joint fMRI and MEG in 335 adults. Hum. Brain Mapp. 36, 2248–2269. https://doi.org/10.1002/HBM.22768

Tustison, N.J., Cook, P.A., Klein, A., Song, G., Das, S.R., Duda, J.T., Kandel, B.M., van Strien, N., Stone, J.R., Gee, J.C., Avants, B.B., 2014. Large-scale evaluation of ANTs and FreeSurfer cortical thickness measurements. Neuroimage 99, 166–179. https://doi.org/10.1016/J.NEUROIMAGE.2014.05.044

Urback, A.L., MacIntosh, B.J., Goldstein, B.I., 2017. Cerebrovascular reactivity measured by functional magnetic resonance imaging during breath-hold challenge: A systematic review. Neurosci. Biobehav. Rev. https://doi.org/10.1016/j.neubiorev.2017.05.003

Václavů, L., Meynart, B.N., Mutsaerts, H.J.M.M., Petersen, E.T., Majoie, C.B.L.M., Vanbavel, E.T., Wood, J.C., Nederveen, A.J., Biemond, B.J., 2019. Hemodynamic provocation with acetazolamide shows impaired cerebrovascular reserve in adults with sickle cell disease. Haematologica 104, 690–699. https://doi.org/10.3324/haematol.2018.206094

van der Zwaag, W., Jorge, J., Butticaz, D., Gruetter, R., 2015. Physiological noise in human cerebellar fMRI. Magn. Reson. Mater. Physics, Biol. Med. 28, 485–492. https://doi.org/10.1007/S10334-015-0483-6/FIGURES/3

van Niftrik, C.H.B., Piccirelli, M., Bozinov, O., Pangalu, A., Valavanis, A., Regli, L., Fierstra, J., 2016. Fine tuning breath-hold-based cerebrovascular reactivity analysis models. Brain Behav. 6, 1–13. https://doi.org/10.1002/brb3.426

Vogt, K.M., Ibinson, J.W., Schmalbrock, P., Small, R.H., 2011. Comparison between end-tidal CO2 and respiration volume per time for detecting BOLD signal fluctuations during paced hyperventilation. Magn. Reson. Imaging 29, 1186–1194. https://doi.org/10.1016/J.MRI.2011.07.011

Wise, R.G., Ide, K., Poulin, M.J., Tracey, I., 2004. Resting fluctuations in arterial carbon dioxide induce significant low frequency variations in BOLD signal. Neuroimage 21, 1652–1664. https://doi.org/10.1016/j.neuroimage.2003.11.025

Zacà, D., Jovicich, J., Nadar, S.R., Voyvodic, J.T., Pillai, J.J., 2014. Cerebrovascular reactivity mapping in patients with low grade gliomas undergoing presurgical sensorimotor mapping with BOLD fMRI. J. Magn. Reson. Imaging 40, 383–390. https://doi.org/10.1002/jmri.24406

