## Supplementary Material for "Comparing end-tidal CO_2_, respiration volume per time (RVT), and average gray matter signal for mapping cerebrovascular reactivity amplitude and delay with breath-hold task BOLD fMRI"

### 1. Reference Signals

**Table S1: Maximum cross-correlations between each pair of reference signals, when timeseries were shifted across a lag range from ±9 seconds**

| Sufficient $P_{ET}CO_2$ Datasets | | | | | | | |
|---|---|---|---|---|---|---|---|
| | | $P_{ET}CO_2$ & RVT | | $P_{ET}CO_2$ & GM-BOLD | | GM-BOLD & RVT | |
| Subject | Session | r | Z | r | Z | r | Z |
| sub-002 | ses-02 | 0.66 | 0.79 | 0.84 | 1.23 | 0.46 | 0.50 |
| | ses-03 | 0.74 | 0.95 | 0.84 | 1.22 | 0.69 | 0.85 |
| sub-003 | ses-02 | 0.83 | 1.18 | 0.88 | 1.39 | 0.89 | 1.43 |
| | ses-03 | 0.81 | 1.13 | 0.85 | 1.27 | 0.77 | 1.01 |
| sub-004 | ses-02 | 0.77 | 1.03 | 0.90 | 1.45 | 0.81 | 1.13 |
| | ses-03 | 0.69 | 0.85 | 0.91 | 1.50 | 0.75 | 0.98 |
| sub-006 | ses-02 | 0.58 | 0.66 | 0.71 | 0.90 | 0.80 | 1.10 |
| | ses-03 | 0.52 | 0.58 | 0.81 | 1.12 | 0.64 | 0.75 |
| sub-007 | ses-02 | 0.82 | 1.15 | 0.87 | 1.31 | 0.86 | 1.31 |
| | ses-03 | 0.87 | 1.34 | 0.90 | 1.46 | 0.89 | 1.44 |
| sub-008 | ses-02 | 0.69 | 0.84 | 0.68 | 0.83 | 0.77 | 1.02 |
| | ses-03 | 0.81 | 1.13 | 0.85 | 1.25 | 0.73 | 0.92 |
| sub-009 | ses-02 | 0.83 | 1.17 | 0.82 | 1.14 | 0.86 | 1.30 |
| | ses-03 | 0.82 | 1.15 | 0.81 | 1.14 | 0.78 | 1.04 |
| sub-010 | ses-02 | 0.58 | 0.67 | 0.68 | 0.83 | 0.79 | 1.06 |
| | ses-03 | 0.63 | 0.73 | 0.73 | 0.93 | 0.71 | 0.88 |
| Average | | | 0.96* | | 1.19* | | 1.04* |
| StDev | | | 0.23 | | 0.22 | | 0.25 |

| Insufficient $P_{ET}CO_2$ Datasets | | | | | | | |
|---|---|---|---|---|---|---|---|
| | | $P_{ET}CO_2$ & RVT | | $P_{ET}CO_2$ & GM-BOLD | | GM-BOLD & RVT | |
| Subject | Session | r | Z | r | Z | r | Z |
| sub-006 | ses-07 | 0.45 | 0.48 | 0.51 | 0.56 | 0.75 | 0.98 |
| | ses-08 | 0.22 | 0.23 | 0.49 | 0.54 | 0.71 | 0.90 |
| sub-009 | ses-08 | 0.26 | 0.27 | 0.19 | 0.20 | 0.90 | 1.47 |
| | ses-09 | 0.37 | 0.39 | 0.31 | 0.33 | 0.87 | 1.33 |
| sub-010 | ses-07 | 0.36 | 0.37 | 0.35 | 0.36 | 0.84 | 1.22 |
| | ses-08 | 0.52 | 0.58 | 0.49 | 0.54 | 0.86 | 1.30 |
| Average | | | 0.38 | | 0.42 | | 1.20* |
| StDev | | | 0.13 | | 0.15 | | 0.22 |

*The maximum Pearson correlation coefficient (r) transformed to Fisher's Z (Z), are reported for datasets with sufficient $P_{ET}CO_2$ quality (top) and insufficient $P_{ET}CO_2$ quality (bottom). Sufficient $P_{ET}CO_2$ quality is*

*determined by >50% relative power in the breath-hold task frequency range. *indicates average Fisher's Z is significant at alpha = 0.05 (critical Z = 0.54 for N = 16; critical Z = 1.13 for N = 8).*

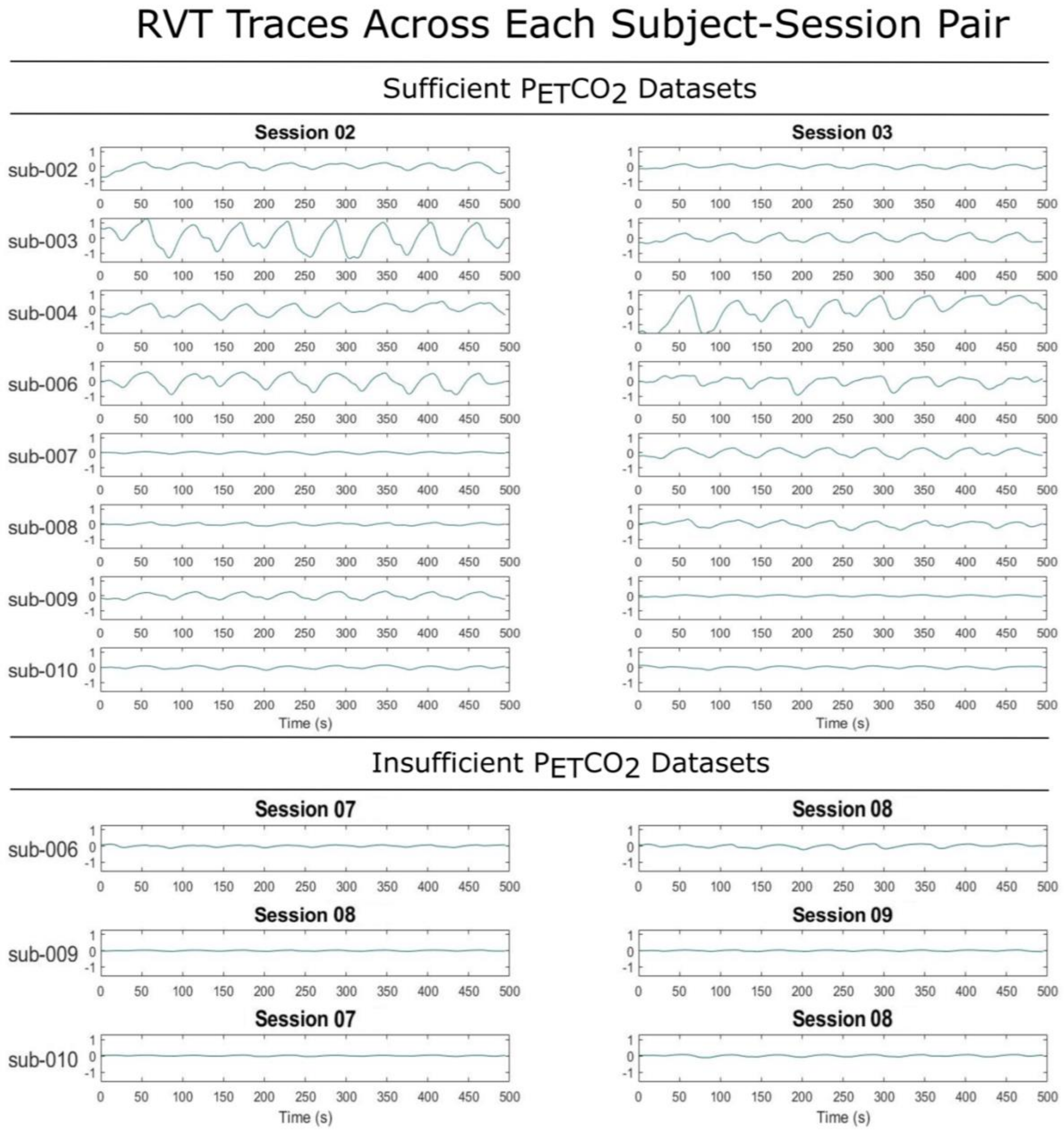

***Figure S1:*** *RVT convolved with the respiration response function for all datasets. RVT signals from datasets with sufficient $P_{ET}CO_2$ are plotted on top, while those with insufficient $P_{ET}CO_2$ are plotted on the bottom. Note the variation in amplitude between datasets*

***Table S2:*** *Maximum peak amplitude from baseline of the RVT reference signal for each subject-session pair. Baseline is considered the average RVT signal over the eight paced breathing sections per subject-session pair, and the peak amplitude is the average across the eight peaks following breath holds per subject-session pair.*

| **Sufficient $P_{ET}CO_2$ Datasets** | | |
|---|---|---|
| Subject | Session | ΔRVT (a.u.) |
| sub-002 | ses-02 | 0.36 |
| | ses-03 | 0.23 |
| sub-003 | ses-02 | 1.60 |
| | ses-03 | 0.51 |
| sub-004 | ses-02 | 0.64 |
| | ses-03 | 1.21 |
| sub-006 | ses-02 | 0.90 |
| | ses-03 | 0.49 |
| sub-007 | ses-02 | 0.12 |
| | ses-03 | 0.54 |
| sub-008 | ses-02 | 0.16 |
| | ses-03 | 0.34 |
| sub-009 | ses-02 | 0.42 |
| | ses-03 | 0.11 |
| sub-010 | ses-02 | 0.18 |
| | ses-03 | 0.13 |
| Average | | 0.50 |
| StDev | | 0.42 |
| **Insufficient $P_{ET}CO_2$ Datasets** | | |
| Subject | Session | ΔRVT (a.u.) |
| sub-006 | ses-07 | 0.11 |
| | ses-08 | 0.17 |
| sub-009 | ses-08 | 0.05 |
| | ses-09 | 0.07 |
| sub-010 | ses-07 | 0.05 |
| | ses-08 | 0.10 |
| Average | | 0.09 |
| StDev | | 0.05 |

### 2. Sufficient $P_{ET}CO_2$ Data: CVR Amplitude

***Table S3:*** *98th percentile CVR amplitude value for each reference signal in datasets with sufficient* $P_{ET}CO_2$

| 98th Percentile CVR Amplitude | | | | |
|---|---|---|---|---|
| **Subject** | **Session** | **$P_{ET}CO_2$ (%BOLD/mmHg)** | **RVT (%BOLD/a.u.)** | **GM-BOLD (%BOLD/%BOLD)** |
| sub-002 | ses-02 | 0.70 | 2.19 | 2.41 |
| | ses-03 | 0.81 | 2.19 | 2.40 |
| sub-003 | ses-02 | 0.62 | 2.35 | 2.29 |
| | ses-03 | 0.90 | 3.03 | 2.53 |
| sub-004 | ses-02 | 0.68 | 2.22 | 2.09 |
| | ses-03 | 0.84 | 2.48 | 2.09 |
| sub-006 | ses-02 | 1.12 | 2.00 | 2.20 |
| | ses-03 | 0.75 | 1.22 | 2.25 |
| sub-007 | ses-02 | 0.55 | 1.48 | 2.68 |
| | ses-03 | 0.65 | 1.70 | 2.57 |
| sub-008 | ses-02 | 0.45 | 1.61 | 1.91 |
| | ses-03 | 0.54 | 1.59 | 2.05 |
| sub-009 | ses-02 | 1.31 | 1.95 | 2.45 |
| | ses-03 | 0.98 | 2.22 | 2.36 |
| sub-010 | ses-02 | 0.72 | 1.95 | 2.47 |
| | ses-03 | 0.81 | 1.61 | 2.28 |
| Average | | 0.78 | 1.99 | 2.31 |
| StDev | | 0.22 | 0.45 | 0.21 |

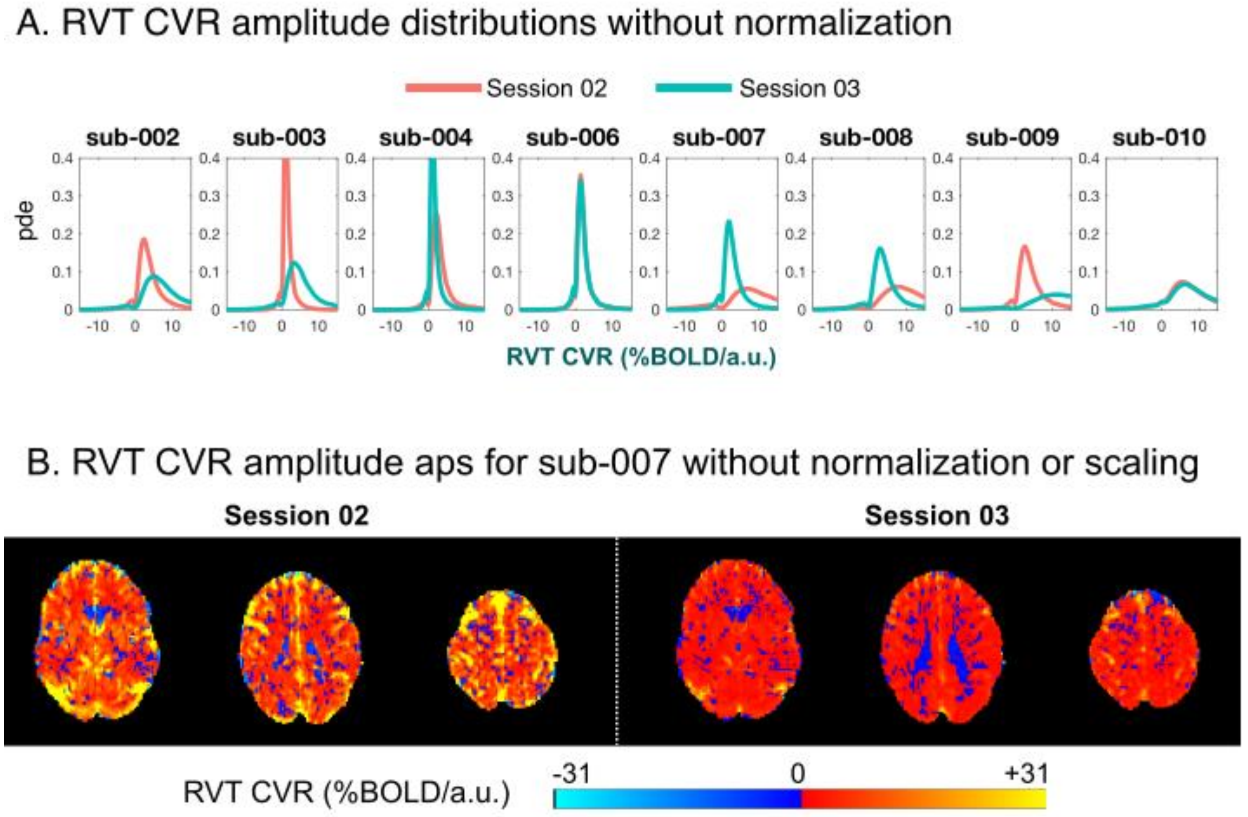

***Figure S2: A)*** *RVT CVR amplitude distributions for all sufficient $P_{ET}CO_2$ datasets, without normalizing RVT regressors to unit variance. There is notable variability in the range of amplitudes, both within and between subjects.* ***B)*** *RVT CVR amplitude maps for an example subject (sub-007) with large variability in RVT amplitudes between two sessions. Because CVR amplitude is scaled to the amplitude of the reference signal, variability in the RVT reference signal translates to variability in the RVT CVR amplitude. Both maps are plotted using the 98$^{th}$ percentile CVR amplitude value for session 02 as the limits of the color scale (30.87 %BOLD/a.u.). The 98$^{th}$ percentile CVR amplitude for session 03 is much smaller (7.93% BOLD/a.u.), and when plotted on the same scale, appears to lose contrast captured by the session 02 map.*

***Table S4:*** *Inter-reference spatial correlations between CVR amplitude maps: relationship between amplitude values from each reference signal in datasets with sufficient $P_{ET}CO_2$*

| | | Inter-Reference Spatial Correlations: CVR Amplitude | | | | | | | | | | | |
|---|---|---|---|---|---|---|---|---|---|---|---|---|---|
| | | $P_{ET}CO_2$ & RVT | | | | $P_{ET}CO_2$ & GM-BOLD | | | | GM-BOLD & RVT | | | |
| Subject | Session | β | Int | r | Z | β | Int | r | Z | β | Int | r | Z |
| sub-002 | ses-02 | 2.66 | 0.07 | 0.98 | 2.28 | 3.22 | 0.04 | 0.99 | 2.93 | 0.82 | 0.04 | 0.98 | 2.26 |
| | ses-03 | 2.56 | 0.02 | 0.99 | 2.65 | 2.76 | 0.04 | 0.99 | 2.84 | 0.92 | -0.02 | 0.99 | 2.62 |
| sub-003 | ses-02 | 3.66 | 0.01 | 0.98 | 2.27 | 3.75 | -0.02 | 0.99 | 2.70 | 0.97 | 0.03 | 0.98 | 2.38 |
| | ses-03 | 3.16 | 0.06 | 0.99 | 2.68 | 2.52 | 0.10 | 0.98 | 2.22 | 1.22 | -0.04 | 0.99 | 2.44 |
| sub-004 | ses-02 | 2.91 | 0.07 | 0.98 | 2.26 | 2.86 | 0.04 | 0.98 | 2.29 | 1.01 | 0.03 | 0.99 | 2.97 |
| | ses-03 | 2.93 | -0.03 | 0.99 | 2.57 | 2.41 | 0.01 | 0.96 | 1.94 | 1.10 | 0.05 | 0.93 | 1.67 |
| sub-006 | ses-02 | 1.79 | -0.01 | 0.99 | 2.99 | 1.96 | -0.01 | 0.99 | 2.70 | 0.90 | 0.01 | 0.99 | 2.85 |
| | ses-03 | 1.31 | 0.08 | 0.84 | 1.23 | 2.86 | 0.03 | 0.98 | 2.41 | 0.44 | 0.08 | 0.83 | 1.18 |
| sub-007 | ses-02 | 1.94 | 0.05 | 0.77 | 1.03 | 4.26 | 0.12 | 0.97 | 2.06 | 0.48 | -0.03 | 0.84 | 1.21 |
| | ses-03 | 2.58 | 0.00 | 0.98 | 2.25 | 3.76 | 0.02 | 0.98 | 2.25 | 0.68 | -0.02 | 0.99 | 2.77 |
| sub-008 | ses-02 | 2.49 | 0.18 | 0.77 | 1.02 | 2.97 | 0.22 | 0.77 | 1.02 | 0.82 | 0.00 | 0.98 | 2.36 |
| | ses-03 | 2.73 | 0.01 | 0.98 | 2.21 | 3.82 | -0.01 | 0.94 | 1.74 | 0.66 | 0.05 | 0.96 | 1.91 |
| sub-009 | ses-02 | 1.47 | 0.00 | 0.99 | 2.75 | 1.79 | 0.02 | 0.99 | 2.78 | 0.81 | 0.00 | 0.99 | 2.64 |
| | ses-03 | 1.87 | 0.09 | 0.96 | 1.93 | 2.13 | 0.07 | 0.98 | 2.23 | 0.88 | 0.03 | 0.98 | 2.37 |
| sub-010 | ses-02 | 2.55 | 0.01 | 0.98 | 2.30 | 3.20 | 0.07 | 0.98 | 2.35 | 0.78 | -0.03 | 0.97 | 2.08 |
| | ses-03 | 1.12 | 0.07 | 0.68 | 0.82 | 2.23 | 0.18 | 0.93 | 1.66 | 0.45 | 0.02 | 0.64 | 0.76 |
| | Average | 2.36 | 0.04 | | 2.08* | 2.91 | 0.06 | | 2.26* | 0.81 | 0.01 | | 2.15* |
| | StDev | 0.71 | 0.05 | | 0.68 | 0.73 | 0.07 | | 0.50 | 0.23 | 0.03 | | 0.65 |

*β = coefficient of slope for best-fit line to correlation. Int = intercept for best-fit line to correlation. r = Pearson correlation coefficient. Z = Fisher's Z transformation of r. *indicates average Fisher's Z is significant at alpha = 0.05 (critical Z = 0.54 for N = 16).*

***Table S5:*** *Inter-session spatial correlations between CVR amplitude maps: relationship between two consecutive datasets with sufficient $P_{ET}CO_2$*

| | | Inter-Session Spatial Correlations: CVR Amplitude | | | | | | | | | | | |
|---|---|---|---|---|---|---|---|---|---|---|---|---|---|
| Subject | Sessions | $P_{ET}CO_2$ | | | | RVT | | | | GM-BOLD | | | |
| | | β | Int | r | Z | β | Int | r | Z | β | Int | r | Z |
| sub-002 | ses-02 & ses-03 | 1.15 | 0.00 | 0.96 | 1.96 | 1.09 | -0.05 | 0.95 | 1.84 | 0.99 | 0.00 | 0.96 | 2.01 |
| sub-003 | ses-02 & ses-03 | 1.38 | -0.03 | 0.94 | 1.78 | 1.15 | -0.01 | 0.92 | 1.60 | 0.91 | 0.05 | 0.91 | 1.56 |
| sub-004 | ses-02 & ses-03 | 1.14 | 0.03 | 0.94 | 1.74 | 1.15 | -0.02 | 0.96 | 1.89 | 0.97 | 0.05 | 0.93 | 1.66 |
| sub-006 | ses-02 & ses-03 | 0.68 | -0.01 | 0.94 | 1.74 | 0.48 | 0.09 | 0.78 | 1.03 | 1.02 | 0.00 | 0.96 | 1.93 |
| sub-007 | ses-02 & ses-03 | 1.00 | 0.04 | 0.93 | 1.64 | 0.96 | 0.17 | 0.85 | 1.26 | 0.90 | 0.07 | 0.95 | 1.87 |
| sub-008 | ses-02 & ses-03 | 0.82 | 0.07 | 0.78 | 1.04 | 0.86 | 0.06 | 0.95 | 1.80 | 1.04 | 0.03 | 0.94 | 1.71 |
| sub-009 | ses-02 & ses-03 | 0.78 | 0.01 | 0.95 | 1.87 | 1.00 | 0.09 | 0.93 | 1.69 | 0.93 | 0.07 | 0.94 | 1.78 |
| sub-010 | ses-02 & ses-03 | 1.05 | 0.01 | 0.83 | 1.19 | 0.57 | 0.02 | 0.70 | 0.86 | 0.84 | 0.08 | 0.90 | 1.46 |
| | Average | 1.00 | 0.01 | | 1.62* | 0.91 | 0.04 | | 1.50* | 0.95 | 0.04 | | 1.75* |
| | StDev | 0.23 | -0.03 | | 0.33 | 0.26 | 0.07 | | 0.39 | 0.07 | 0.03 | | 0.19 |

*β = coefficient of slope for best-fit line to correlation. Int = intercept for best-fit line to correlation. r = square of Pearson correlation coefficient. Z = Fisher's Z transformation of r. *indicates average Fisher's Z is significant at alpha = 0.05 (critical Z = 0.88 for N = 8).*

### 3. Sufficient $P_{ET}CO_2$ Data: CVR Delay

***Table S6:*** *Inter-reference spatial correlations between CVR delay maps: relationship between delay values from each reference signal in datasets with sufficient* $P_{ET}CO_2$

| Inter-Reference Spatial Correlations: CVR Delay | | | | | | | | | | | | | |
|---|---|---|---|---|---|---|---|---|---|---|---|---|---|
| | | $P_{ET}CO_2$ & RVT | | | | $P_{ET}CO_2$ & GM-BOLD | | | | GM-BOLD & RVT | | | |
| **Subject** | **Session** | β | Int | r | Z | β | Int | r | Z | β | Int | r | Z |
| sub-002 | ses-02 | 0.01 | -0.45 | 0.01 | 0.01 | 0.43 | -0.07 | 0.90 | 1.48 | 0.42 | -0.33 | 0.21 | 0.21 |
| | ses-03 | 0.76 | 0.03 | 0.55 | 0.62 | 0.63 | -0.36 | 0.92 | 1.61 | 1.18 | 0.45 | 0.58 | 0.67 |
| sub-003 | ses-02 | 0.81 | -0.16 | 0.84 | 1.23 | 0.68 | -0.15 | 0.93 | 1.67 | 1.05 | -0.05 | 0.80 | 1.10 |
| | ses-03 | 1.05 | -0.16 | 0.85 | 1.24 | 0.64 | -0.05 | 0.89 | 1.45 | 1.58 | -0.12 | 0.91 | 1.53 |
| sub-004 | ses-02 | 1.07 | 0.10 | 0.86 | 1.28 | 0.79 | -0.07 | 0.92 | 1.59 | 1.30 | 0.16 | 0.89 | 1.42 |
| | ses-03 | 1.19 | 0.13 | 0.92 | 1.57 | 0.81 | 0.09 | 0.93 | 1.67 | 1.25 | -0.10 | 0.83 | 1.20 |
| sub-006 | ses-02 | 1.01 | 0.06 | 0.92 | 1.60 | 0.73 | -0.17 | 0.87 | 1.33 | 1.11 | 0.02 | 0.85 | 1.25 |
| | ses-03 | 0.58 | -0.69 | 0.40 | 0.42 | 0.81 | -0.25 | 0.81 | 1.11 | 0.78 | -0.47 | 0.54 | 0.60 |
| sub-007 | ses-02 | 0.87 | -0.48 | 0.49 | 0.53 | 0.64 | -0.17 | 0.70 | 0.87 | 1.55 | -0.16 | 0.79 | 1.06 |
| | ses-03 | 0.80 | -0.41 | 0.67 | 0.81 | 0.55 | -0.32 | 0.86 | 1.30 | 1.63 | 0.17 | 0.87 | 1.34 |
| sub-008 | ses-02 | 1.27 | 0.39 | 0.93 | 1.69 | 0.38 | 0.30 | 0.73 | 0.94 | 2.12 | -1.45 | 0.82 | 1.15 |
| | ses-03 | 1.42 | 0.04 | 0.95 | 1.88 | 0.47 | 0.27 | 0.82 | 1.16 | 2.00 | -1.30 | 0.77 | 1.01 |
| sub-009 | ses-02 | 1.38 | 0.02 | 0.96 | 1.89 | 0.59 | -0.01 | 0.90 | 1.48 | 1.99 | -0.09 | 0.89 | 1.44 |
| | ses-03 | 0.90 | -0.04 | 0.82 | 1.15 | 0.47 | -0.08 | 0.77 | 1.02 | 1.58 | -0.08 | 0.88 | 1.37 |
| sub-010 | ses-02 | 1.03 | 0.16 | 0.97 | 2.02 | 0.63 | -0.19 | 0.89 | 1.40 | 1.25 | -0.01 | 0.83 | 1.19 |
| | ses-03 | 0.65 | 0.95 | 0.77 | 1.02 | 0.66 | -0.46 | 0.91 | 1.53 | 0.81 | 1.20 | 0.70 | 0.86 |
| | Average | 0.93 | -0.03 | | 1.18* | 0.62 | -0.11 | | 1.35* | 1.35 | -0.14 | | 1.09* |
| | StDev | 0.35 | 0.38 | | 0.58 | 0.13 | 0.21 | | 0.26 | 0.47 | 0.61 | | 0.35 |

*β = coefficient of slope for best-fit line to correlation. Int = intercept for best-fit line to correlation. r = Pearson correlation coefficient. Z = Fisher's Z transformation of r. *indicates average Fisher's Z is significant at alpha = 0.05 (critical Z = 0.54 for N = 16).*

***Table S7:*** *Inter-session spatial correlations between CVR delay maps: relationship between two consecutive datasets with sufficient* $P_{ET}CO_2$

| Inter-Session Spatial Correlations: CVR Delay | | | | | | | | | | | | | |
|---|---|---|---|---|---|---|---|---|---|---|---|---|---|
| Subject | Sessions | $P_{ET}CO_2$ | | | | RVT | | | | GM-BOLD | | | |
| | | β | Int | r | Z | β | Int | r | Z | β | Int | r | Z |
| sub-002 | ses-02 & ses-03 | 0.52 | 0.00 | 0.74 | 0.95 | 0.78 | 0.19 | 0.78 | 1.05 | 0.80 | -0.29 | 0.81 | 1.13 |
| sub-003 | ses-02 & ses-03 | 0.82 | -0.52 | 0.68 | 0.83 | 1.01 | -0.57 | 0.64 | 0.77 | 0.93 | -0.18 | 0.79 | 1.06 |
| sub-004 | ses-02 & ses-03 | 0.99 | -0.14 | 0.84 | 1.22 | 1.05 | -0.17 | 0.86 | 1.28 | 1.01 | 0.04 | 0.84 | 1.23 |
| sub-006 | ses-02 & ses-03 | 0.59 | 0.18 | 0.70 | 0.88 | 0.48 | -0.44 | 0.44 | 0.47 | 0.78 | 0.14 | 0.79 | 1.07 |
| sub-007 | ses-02 & ses-03 | 1.07 | -0.08 | 0.79 | 1.07 | 0.78 | -0.19 | 0.85 | 1.27 | 0.77 | -0.28 | 0.80 | 1.11 |
| sub-008 | ses-02 & ses-03 | 0.74 | 0.31 | 0.85 | 1.26 | 0.89 | 0.34 | 0.94 | 1.72 | 0.87 | 0.12 | 0.92 | 1.59 |
| sub-009 | ses-02 & ses-03 | 1.12 | -0.38 | 0.81 | 1.12 | 0.93 | -0.23 | 0.88 | 1.36 | 1.05 | -0.20 | 0.80 | 1.11 |
| sub-010 | ses-02 & ses-03 | 0.76 | 0.25 | 0.91 | 1.53 | 0.55 | 1.14 | 0.83 | 1.19 | 0.75 | -0.20 | 0.88 | 1.39 |
| | Average | 0.83 | 0.05 | | 1.11* | 0.81 | 0.01 | | 1.14* | 0.87 | 0.11 | | 1.21* |
| | StDev | 0.22 | 0.30 | | 0.23 | 0.21 | 0.55 | | 0.38 | 0.11 | 0.18 | | 0.19 |

*β = coefficient of slope for best-fit line to correlation. Int = intercept for best-fit line to correlation. r = square of Pearson correlation coefficient. Z = Fisher's Z transformation of r. *indicates average Fisher's Z is significant at alpha = 0.05 (critical Z = 0.88 for N = 8).*

### 4. Insufficient $P_{ET}CO_2$ Data

***Table S8:*** *98th percentile CVR amplitude value for each reference signal in datasets with insufficient $P_{ET}CO_2$*

| | | 98th Percentile CVR Amplitude | | |
|---|---|---|---|---|
| **Subject** | **Session** | **$P_{ET}CO_2$ (%BOLD/mmHg)** | **RVT (%BOLD/a.u.)** | **GM-BOLD (%BOLD/%BOLD)** |
| sub-006 | ses-07 | 1.57 | 2.42 | 2.25 |
| | ses-08 | 1.16 | 2.16 | 2.33 |
| sub-009 | ses-08 | 0.15 | 2.07 | 2.50 |
| | ses-09 | 0.75 | 1.83 | 2.48 |
| sub-010 | ses-07 | 1.04 | 2.32 | 2.46 |
| | ses-08 | 0.91 | 1.99 | 2.50 |
| | Average | 0.93 | 2.13 | 2.42 |
| | StDev | 0.47 | 0.22 | 0.10 |

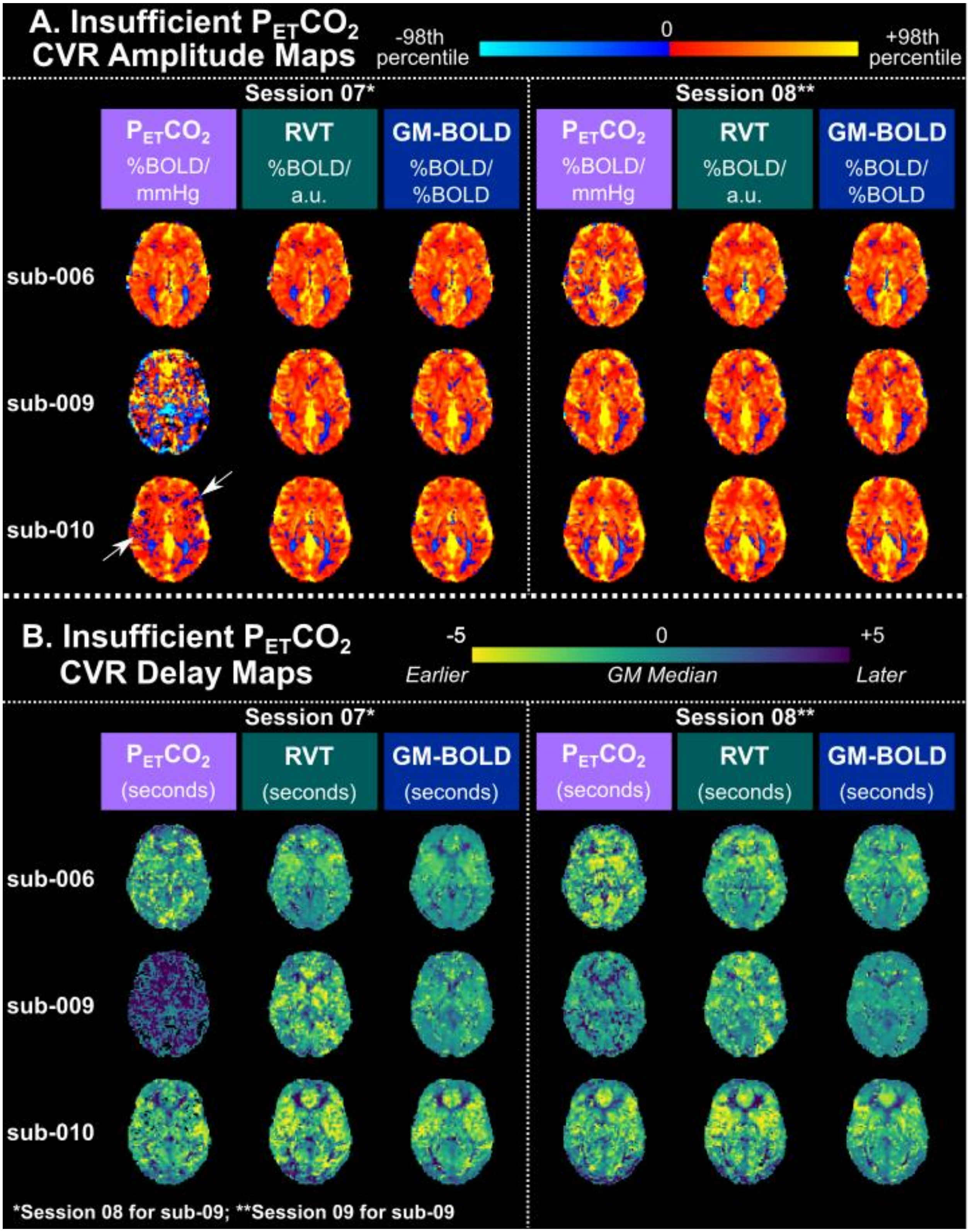

***Figure S3****: CVR maps for all datasets with insufficient $P_{ET}CO_2$ quality, transformed to the MNI152 6$^{th}$ generation template space. A) Delay-optimized CVR amplitude maps are scaled to the 98$^{th}$ percentile value across all brain voxels. White arrows indicate regions of negative CVR that could be mis-*

*characterized as the vascular "steal" phenomenon, indicating that maps with insufficient $P_{ET}CO_2$ should be interpreted carefully. B) CVR delay maps are normalized to the median delay in gray matter (GM). Voxels with delays at the boundary conditions (absolute delay = +/- 8.7s, 9s have been removed from all maps.*

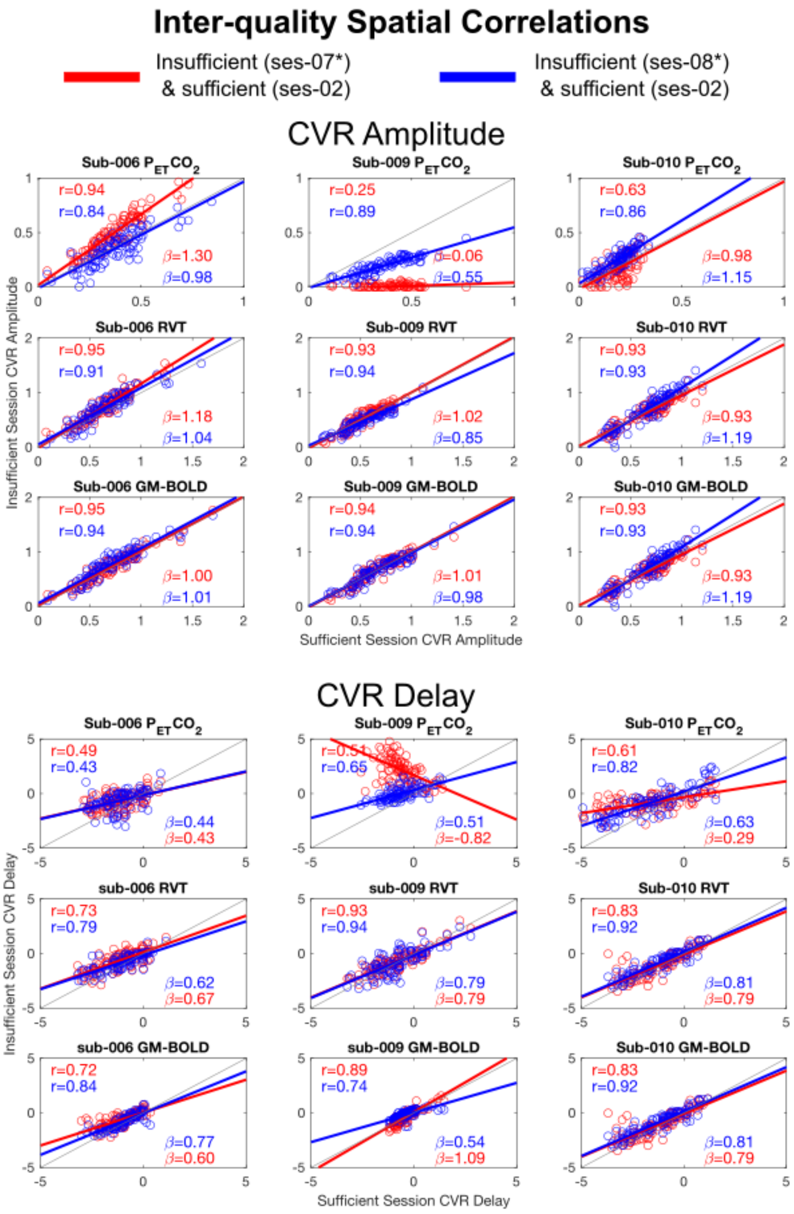

***Figure S4:*** *Inter-quality spatial correlations for all datasets with insufficient $P_{ET}CO_2$ quality. Each subplot shows the spatial correlation between CVR parameter maps from an insufficient $P_{ET}CO_2$ quality dataset and a corresponding sufficient $P_{ET}CO_2$ dataset in the same subject. A) CVR amplitude and B) CVR delay spatial correlations. Pearson correlation coefficients (r) are listed in the top right corner and slopes for the*

lines-of-best-fit (β) are displayed in the bottom right corner. *for sub-009, the insufficient sessions are ses-08 (red) and ses-09 (blue).

***Table S9:*** *Inter-reference spatial correlations between CVR parameter maps: relationship between amplitude or delay values from each reference signal in datasets with insufficient* $P_{ET}CO_2$

| Inter-Reference Spatial Correlations | | | | | | | | | | | | | |
|---|---|---|---|---|---|---|---|---|---|---|---|---|---|
| | | CVR Amplitude | | | | | | CVR Lag | | | | | |
| Subject | Session | $P_{ET}CO_2$ & RVT | | $P_{ET}CO_2$ & GM-BOLD | | GM-BOLD & RVT | | $P_{ET}CO_2$ & RVT | | $P_{ET}CO_2$ & GM-BOLD | | GM-BOLD & RVT | |
| | | β | Z | β | Z | β | Z | β | Z | β | Z | β | Z |
| sub-006 | ses-07 | 1.57 | 2.17 | 1.45 | 2.04 | 1.07 | 2.50 | 0.55 | 0.52 | 0.37 | 0.49 | 1.19 | 1.23 |
| | ses-08 | 1.66 | 1.77 | 1.71 | 1.70 | 0.95 | 2.63 | 0.48 | 0.64 | 0.23 | 0.31 | 0.91 | 1.12 |
| sub-009 | ses-08 | 2.04 | 0.31 | 2.26 | 0.29 | 0.83 | 2.77 | -0.42 | 0.56 | -0.26 | 0.58 | 1.50 | 1.51 |
| | ses-09 | 2.02 | 1.71 | 2.83 | 1.72 | 0.70 | 2.55 | 0.87 | 0.50 | 0.24 | 0.42 | 2.49 | 1.11 |
| sub-010 | ses-07 | 1.26 | 0.77 | 1.44 | 0.85 | 0.92 | 2.52 | 1.19 | 0.77 | 0.93 | 0.80 | 1.23 | 1.65 |
| | ses-08 | 2.37 | 1.90 | 2.96 | 1.86 | 0.79 | 2.38 | 1.01 | 1.50 | 0.72 | 1.34 | 1.21 | 1.44 |
| | Average | 1.82 | 1.44* | 2.11 | 1.41* | 0.88 | 2.56* | 0.61 | 0.75 | 0.37 | 0.66 | 1.42 | 1.34* |
| | StDev | 0.40 | 0.73 | 0.68 | 0.69 | 0.13 | 0.13 | 0.57 | 0.38 | 0.42 | 0.37 | 0.55 | 0.22 |

*β = coefficient of slope for best-fit line to correlation. Z = Fisher Z transformation of correlation coefficient. *Indicates Fisher's Z is significantly different from 0 at alpha = 0.05 (critical Z = 1.13 for N = 6).*

### 5. Comparison of Delays with Refined GM-BOLD Reference Signal

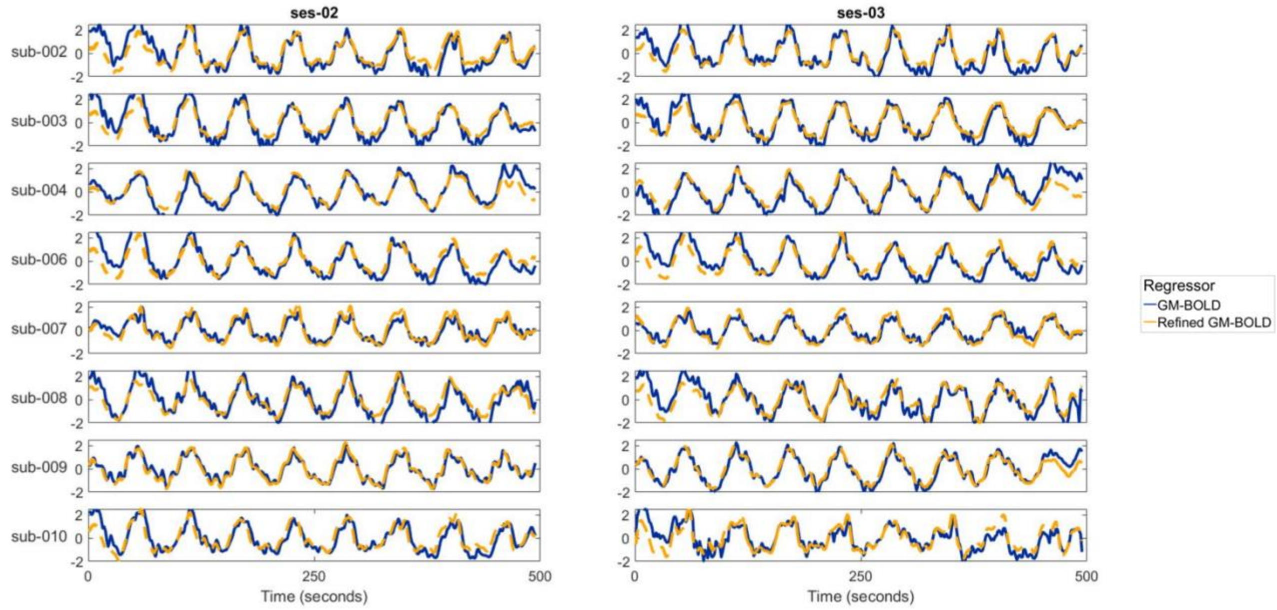

***Figure S5:*** *GM-BOLD (blue) and Refined GM-BOLD (yellow) reference signals for all subjects with sufficient* $P_{ET}CO_2$ *data quality across ses-02 and ses-03.*

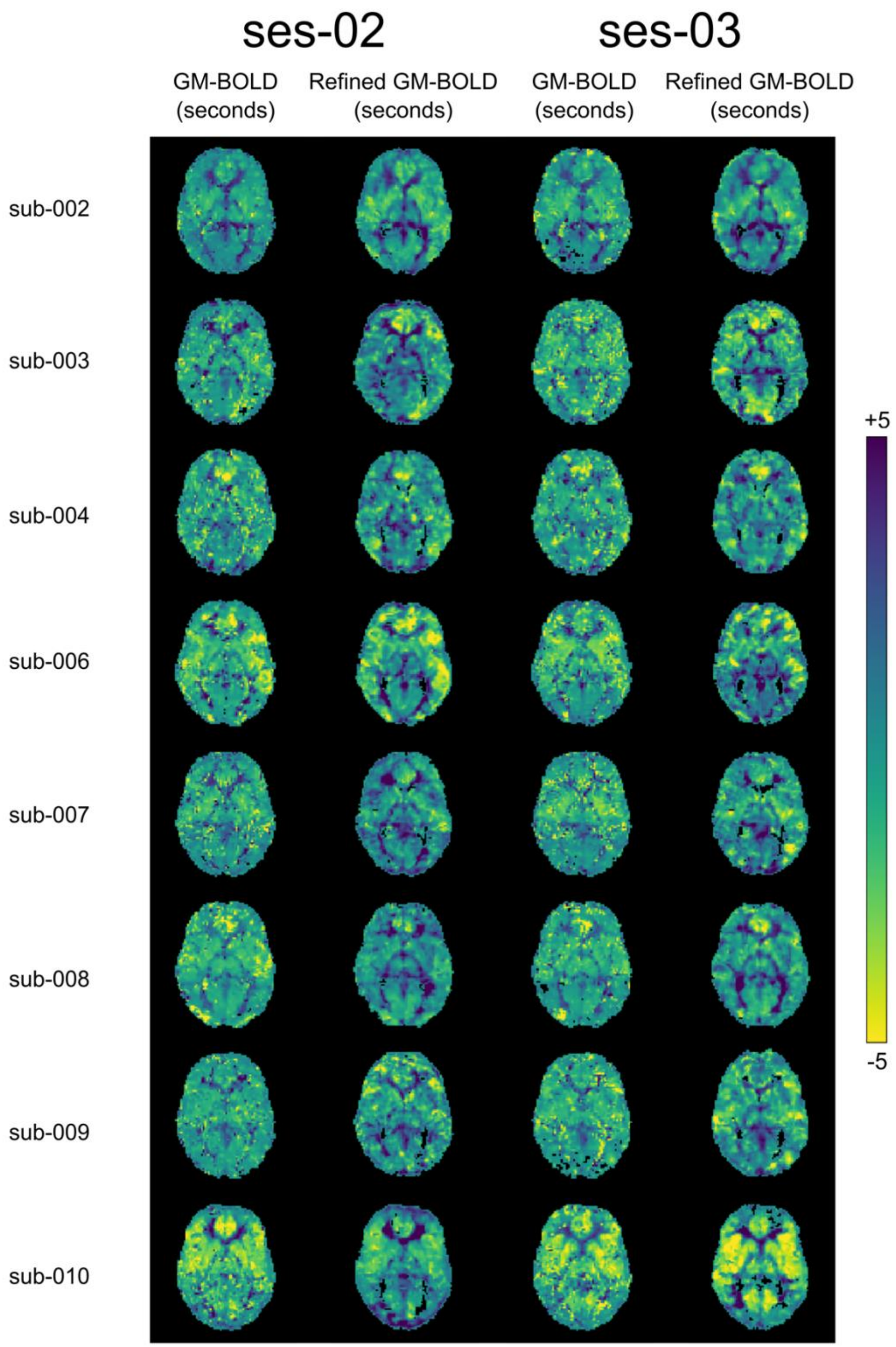

***Figure S6:*** *CVR delay maps for all subjects with sufficient $P_{ET}CO_2$ data quality across ses-02 and ses-03. Maps were processed using either the GM-BOLD or Refined GM-BOLD approaches. The names of the output maps from Rapidtide ended with “maxtime_map”.*

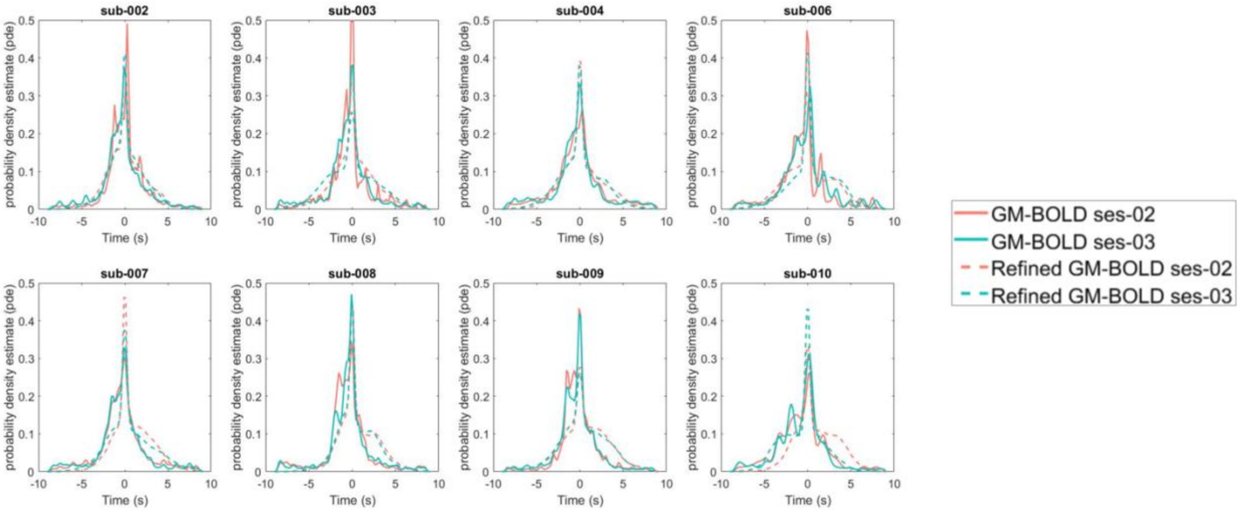

***Figure S7:*** *CVR delay distributions for all subjects with sufficient $P_{ET}CO_2$ data quality across ses-02 and ses-03.*

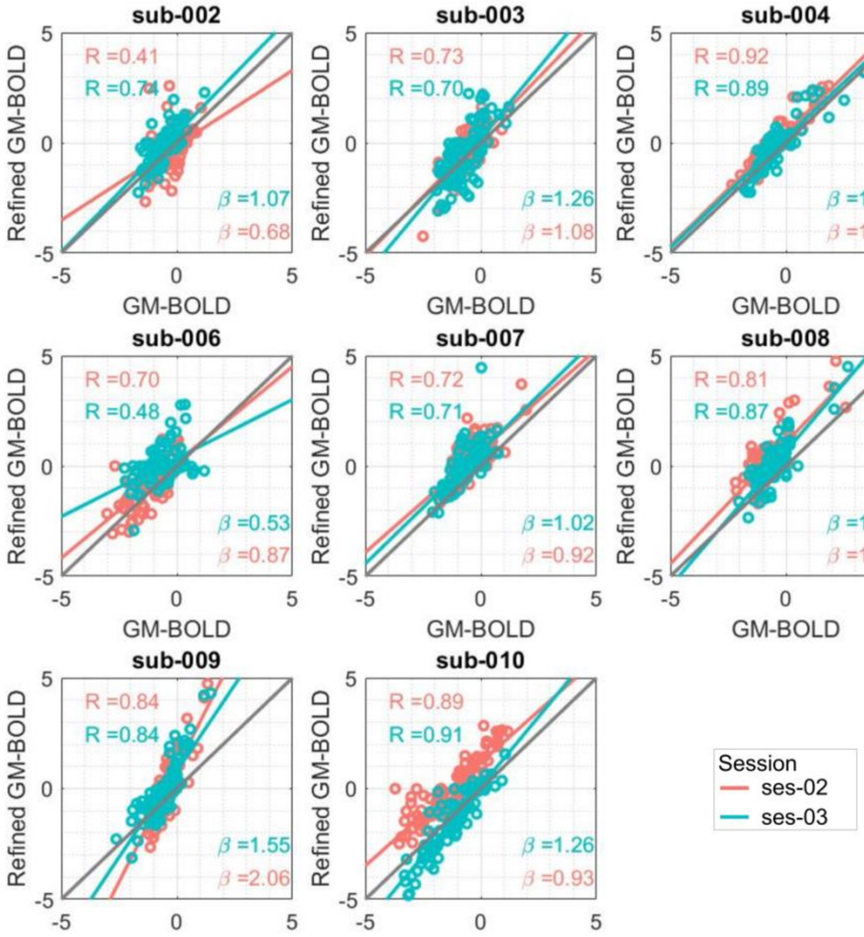

***Figure S8:*** *Inter-reference spatial correlation between GM-BOLD and Refined GM-BOLD delay maps for all subjects with sufficient $P_{ET}CO_2$ data quality across ses-02 (orange) and ses-03 (teal).*

***Table S10:*** *Inter-reference spatial correlation of CVR delay in median cortical parcels between GM-BOLD and Refined GM-BOLD across all subjects with sufficient $P_{ET}CO_2$ data quality for ses-02 and ses-03. $B_1$ is coefficient of the slope for best-fit line to correlation, with intercept at $B_0$. Pearson's correlation coefficient (R) and Fisher's Z (Z) depict the agreement between the GM-BOLD and Refined GM-BOLD approaches.*

**Inter-Reference Spatial Correlations: CVR Delay**

| | | **GM-BOLD & Refined GM-BOLD** | | | |
|---|---|---|---|---|---|
| **Subject** | **Session** | $B_1$ | $B_0$ | R | Z |
| Sub-002 | Ses-02 | 0.68 | -0.11 | 0.41 | 0.44 |
| | Ses-03 | 1.07 | 0.43 | 0.73 | 0.94 |
| Sub-003 | Ses-02 | 1.08 | 0.27 | 0.73 | 0.93 |
| | Ses-03 | 1.26 | 0.26 | 0.7 | 0.87 |
| Sub-004 | Ses-02 | 1.02 | 0.46 | 0.92 | 1.56 |
| | Ses-03 | 1.00 | 0.24 | 0.89 | 1.42 |
| Sub-006 | Ses-02 | 0.87 | 0.17 | 0.71 | 0.88 |
| | Ses-03 | 0.53 | 0.36 | 0.48 | 0.52 |
| Sub-007 | Ses-02 | 0.92 | 0.68 | 0.71 | 0.9 |
| | Ses-03 | 1.02 | 0.66 | 0.71 | 0.88 |
| Sub-008 | Ses-02 | 1.10 | 1.06 | 0.81 | 1.14 |
| | Ses-03 | 1.23 | 0.67 | 0.87 | 1.32 |
| Sub-009 | Ses-02 | 2.06 | 0.95 | 0.84 | 1.24 |
| | Ses-03 | 1.55 | 0.79 | 0.84 | 1.23 |
| Sub-010 | Ses-02 | 0.93 | 1.18 | 0.89 | 1.41 |
| | Ses-03 | 1.26 | 0.11 | 0.91 | 1.54 |
| | Average | 1.10 | 0.51 | 0.76 | 1.08 |
| | StDev | 0.35 | 0.36 | 0.15 | 0.34 |

***Table S11:*** *Rapidtide's delay maps were generated by using preprocessed fMRI data and by running Rapidtide (Frederick, Salo, & Drucker, 2022) with the settings below. Adapted from Gong et al. (2022).*

| **Rapidtide Setting** | **Choice** | **Rationale** |
|---|---|---|
| globalmeaninclude | Path to file with eroded GM mask | Only include the voxels in the mask for generating the global regressor. |
| refineinclude | Path to file with eroded GM mask | Only include the voxels in the mask in refining the global regressor. |
| motionfile | Path to file with 6 columns of motion regressors from volume registration | Denoise the signal from motion sources. |
| motderiv | True | Denoise the signal from motion derivative sources. |
| delaymapping | True | Macro setting that sets defaults that are optimized for delay mapping. |
| searchrange | -9 9 | Limit analysis of lags at lower and upper bounds from -9 to +9 seconds, respectively. |
| passes | 3 | Determines the number of processing passes in the Rapidtide procedure. 3 is the default choice given delaymapping option. |
| despeckle_passes | 4 | 4 is the default choice given delaymapping option. |
| refineoffset | True | True is the default choice given delaymapping option. |
| pickleft | True | Picks the leftmost peak in histogram of delays when setting the refine offset. True is the default choice given delaymapping option. |
| datastep | 1.5 | Set the timestep in seconds to match the TR of the fMRI input data. |
| detrendorder | 4 | Sets the order of trend removal. Consistent with 4th order Legendre polynomials used in lagged-GLM. |
| oversampfac | 5 | Oversample fMRI data by 5, to allow 0.3 second timestep increments in analysis. Consistent with 0.3 second shift increment in lagged-GLM. |
| filterband | lfo | Filter the data and regressors to remove low frequencies (0.0009-0.15 Hz). Default setting in Rapidtide. |
| spatialfilt | -1 | Spatially smooths fMRI data with a Gaussian filter using a sigma equal to half the mean voxel dimension (1.2 mm for our data). Recommended width for spatial smoothing with Rapidtide |

### *6. References*

Frederick, B. deB, Salo, T., & Drucker, D. M. (2022). bbfrederick/rapidtide: Version 2.2.8.1 - 8/29/22 deployment bug fix. https://doi.org/10.5281/ZENODO.7032879

Gong, J., Stickland, R. C., Bright, M. G., & Bright, M. (2022). Hemodynamic timing in resting-state and breathing-task BOLD fMRI. *BioRxiv*, 2022.11.11.516194. https://doi.org/10.1101/2022.11.11.516194
